# Cyclin A/B RxL Macrocyclic Inhibitors to Treat Cancers with High E2F Activity

**DOI:** 10.1101/2024.08.01.605889

**Authors:** Shilpa Singh, Catherine E. Gleason, Min Fang, Yasmin N. Laimon, Vishal Khivansara, Shanhai Xie, Yavuz T. Durmaz, Aniruddha Sarkar, Kenneth Ngo, Varunika Savla, Yixiang Li, Muhannad Abu-Remaileh, Xinyue Li, Bishma Tuladhar, Ranya Odeh, Frances Hamkins-Indik, Daphne He, Miles W. Membreno, Meisam Nosrati, Nathan N. Gushwa, Siegfried S.F. Leung, Breena Fraga-Walton, Luis Hernandez, Miguel P Baldomero, Bryan M. Lent, David Spellmeyer, Joshua F. Luna, Dalena Hoang, Yuliana Gritsenko, Manesh Chand, Megan K. DeMart, Sammy Metobo, Chinmay Bhatt, Justin A. Shapiro, Kai Yang, Nathan J. Dupper, Andrew T. Bockus, John G. Doench, James B. Aggen, Li-Fen Liu, Bernard Levin, Evelyn W. Wang, Iolanda Vendrell, Roman Fischer, Benedikt Kessler, Prafulla C. Gokhale, Sabina Signoretti, Alexander Spektor, Constantine Kreatsoulas, Rajinder Singh, David J. Earp, Pablo D. Garcia, Deepak Nijhawan, Matthew G. Oser

## Abstract

Cancer cell proliferation requires precise control of E2F1 activity; excess activity promotes apoptosis. Here, we developed cell-permeable and bioavailable macrocycles that selectively kill small cell lung cancer (SCLC) cells with inherent high E2F1 activity by blocking RxL-mediated interactions of cyclin A and cyclin B with select substrates. Genome-wide CRISPR/Cas9 knockout and random mutagenesis screens found that cyclin A/B RxL macrocyclic inhibitors (cyclin A/Bi) induced apoptosis paradoxically by cyclin B- and Cdk2-dependent spindle assembly checkpoint activation (SAC). Mechanistically, cyclin A/Bi hyperactivate E2F1 and cyclin B by blocking their RxL-interactions with cyclin A and Myt1, respectively, ultimately leading to SAC activation and mitotic cell death. Base editor screens identified cyclin B variants that confer cyclin A/Bi resistance including several variants that disrupted cyclin B:Cdk interactions. Unexpectedly but consistent with our base editor and knockout screens, cyclin A/Bi induced the formation of neo-morphic Cdk2-cyclin B complexes that promote SAC activation and apoptosis. Finally, orally-bioavailable cyclin A/Bi robustly inhibited tumor growth in chemotherapy-resistant patient-derived xenograft models of SCLC. This work uncovers gain-of-function mechanisms by which cyclin A/Bi induce apoptosis in cancers with high E2F activity, and suggests cyclin A/Bi as a therapeutic strategy for SCLC and other cancers driven by high E2F activity.

## Introduction

Small cell lung cancer (SCLC) is an aggressive form of lung cancer without highly effective targeted therapies^1^. SCLC is genomically characterized by near universal loss of function (LOF) mutations in *RB1* (pRB=protein) and *TP53* without recurrently mutated druggable oncogenic drivers, which has made finding new therapeutic strategies for SCLC more challenging^2–6^. One strategy to identify new therapeutic targets in SCLC is to identify synthetic lethal vulnerabilities with *RB1* and/or *TP53* LOF mutations. Approaches to identify synthetic lethal targets with *RB1* loss in SCLC and other cancers have identified several targets that function in mitosis including Aurora A kinase^7,8^, Aurora B kinase^9^, PLK1^9,10^, and Mps1^11^ together suggesting that cancer cells become hyper-dependent on mitotic kinases upon *RB1* loss.

Unphosphorylated pRB in the G1 phase of the cell cycle functions to bind and repress activating E2Fs including E2F1, E2F2, and E2F3^12–14^. When pRB is phosphorylated, first by Cdk4/6-cyclin D and later by Cdk2-cyclinA/E, its interaction with E2F is inhibited and E2F promotes cell cycle progression through S phase^12–14^. Although activating E2Fs are necessary for cellular proliferation, too much E2F activity paradoxically induces apoptosis^15–20^ suggesting a goldilocks phenomenon. In addition to the negative regulation of activating E2Fs by pRB, E2F1 is phosphorylated by cyclin A/Cdk2 at the end of S phase, which blocks E2Fs ability to bind DNA thereby dampening E2F activity^21–24^. The MRAIL motif of cyclin A forms a Hydrophobic Patch (HP) that serves to recognize and bind an RxL motif on E2F1 to initiate cyclin A/Cdk2-mediated E2F phosphorylation^24,25^. Blocking a cyclin A-E2F RxL interaction genetically or with cell-permeable “tool” peptides that mimic the RxL peptide motif on E2F1 promotes apoptosis selectively in transformed cells with high baseline E2F activity from a dysregulated pRB pathway^26^.

However, efforts to develop small molecule inhibitors to block cyclin/RxL-substrate interactions, and specifically cyclin A /RxL-substrate interactions have not advanced beyond the hit-finding stage due to apparent limitations imposed by synthesis, selectivity, and physicochemical properties^27–33^. Macrocyclic peptides offer a feasible strategy to block broad and shallow PPI surfaces of this type^34^, which until recently have been considered undruggable. In addition, the structural complexity of macrocycles enables significant opportunities to achieve increased selectivity to the target of interest. Furthermore, over the past decade a growing body of research has demonstrated that peptidic macrocycles can be engineered to be both passively cell-permeable^35–40^ and orally bioavailable^41–44^, thus offering promise for this modality to access intracellular targets and usher in a new era of therapeutics^45^.

Many macrocyclic therapeutics are either derived from difficult to analog natural products or have physicochemical properties which limit their application exclusively to extracellular targets^46^. Thus, it has been challenging to design peptidic macrocycles that are potent, selective, stable, cell-permeable and bioavailable to be used as cancer therapeutics. We developed an approach to overcome these obstacles and discovered a series of macrocycles that bind the HP of cyclins with diverse selectivity profiles and then optimized macrocycles that inhibit cyclin A/B RxL interactions for oral bioavailability. Here we report the development of cyclin A/B RxL macrocyclic peptide inhibitors, their activity in SCLC cell lines, cell line xenografts, and patient-derived xenografts, and uncover the mechanisms by which they potently induce apoptosis in SCLCs with high E2F activity.

## Results

### Development of Macrocyclic Peptides that Bind to Cyclin A and/or B Preventing Cyclins from Binding RxL Motifs on Their Substrates

To facilitate the structure-based design of passively permeable peptidic macrocyclic inhibitors of the RxL-motif interactions with the Hydrophobic Patch (HP) of various cyclins (Fig. 1A), we first built a model based on the available structures of a macrocycle (PDB code: 1URC) and p27-Kip1 (PDB code: 1JSU) bound to the cyclin A2/Cdk2 complex (Fig. S1A). A composite molecule (Fig. S1B) containing the lariat macrocycle occupying the larger HP (Fig. S1A) fused to the N-terminal sequences from the RxL-motif of p27-Kip1 occupying a smaller adjacent hydrophobic pocket “S” (Fig. 1A), served as the starting point for structure-based compound design. Forward design and synthesis efforts were then focused on: i) optimizing cyclin affinity; ii) identifying compounds with differing cyclin selectivity profiles to aid elucidation of the impact on biological outcomes; iii) achieving cell permeability with the goal of ultimately enabling oral bioavailability. Cyclin affinity was achieved by structure-based design guided by binding models derived from the published crystal structures. Structural differences between the S & HP pockets of cyclin A, cyclin B, and cyclin E from the corresponding crystal structures (PDB codes 1JSU, 2B9R, 1W98, respectively) were used to tune the selectivity of compounds for individual cyclins. Guided by the computational binding models, rational design approaches were taken to discover hit scaffolds and to optimize high affinity interactions with the respective cyclins by incorporating peptidic and peptidomimetic features. Cell-permeability was achieved by leveraging Circle Pharma’s proprietary MXMO^TM^ platform–an integrated set of synthetic and computational workflows, and models for permeability prediction and optimization of macrocycles based on our previous research^36,38,47,48^. This entailed targeted introduction of structural modifications predicted to improve permeability without negatively impacting cyclin affinity, and by establishing an appropriate balance of physicochemical properties. By applying this approach iteratively, we discovered the set of passively permeable macrocycles with high affinity and diverse selectivity for cyclins (Fig. 1B). A space-filling representation of the binding of compound CIRc-004 to cyclin A2/Cdk2 illustrates the location and orientation of the binding site on cyclin A2, while a stick representation shows how the compound optimally fits into both the HP and S pockets (Fig. 1C, S1C). The details of the medicinal chemistry effort will be published separately.

**Figure 1:**
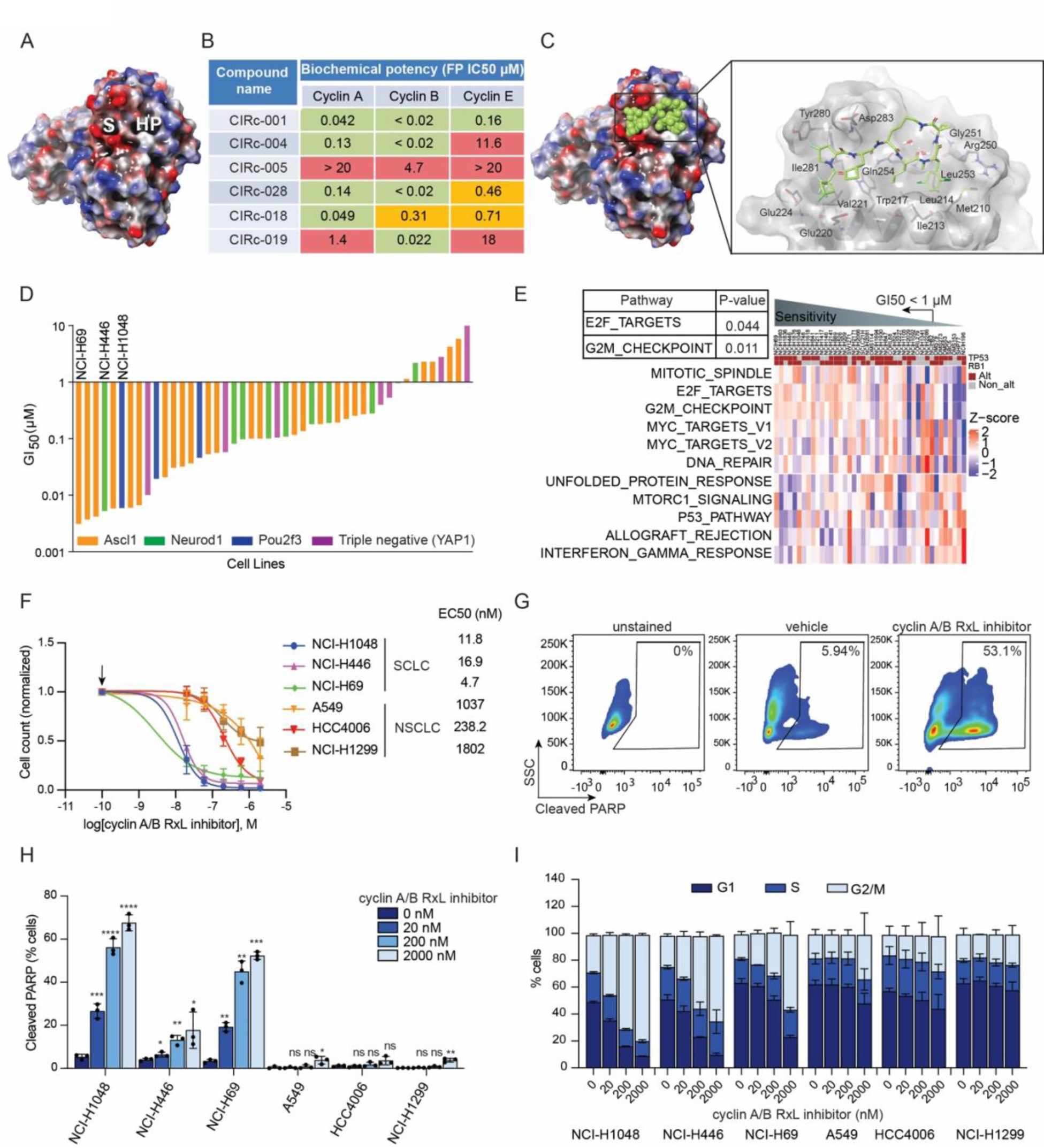
Discovery and Activities of Cyclin RxL Inhibitors. **A)** Structure of cyclin A (PDB code: 1JSU) with blue indicating basic, red acidic and white neutral surfaces. “HP” indicates the Hydrophobic Patch formed in part by the highly conserved MRAIL motif in cyclins and “S” indicates the smaller adjacent hydrophobic pocket. **B)** Docked model of CIRc-004 filled-in structure in green bound to cyclin A. Shown next to it is the detailed “stick” structure representation of CIRc-004 docked to cyclin A. **C)** Biochemical activity of cyclin RxL macrocyclic inhibitors in cyclin A1/Cdk2, cyclinE1/Cdk2 and cyclin B/CDK1 complexes measured by Fluorescence Polarization. **D**) GI_50_ waterfall plot of 46 human SCLC cell lines panel. **E**) Heatmap displays the activity patterns of various Hallmark pathways within a panel of 46 SCLC cell lines. The Gene Set Variation Analysis (GSVA) method, utilizing the MSigDb Hallmark collection of RNA-seq data, was employed to calculate scores for each pathway. Each column represents a distinct SCLC cell line ordered from most sensitive to least sensitive based on GI_50_ values for cyclin RxL A/B inhibitor (CIRc-004). p-value <0.05 are shown and calculated by Wilcoxon rank sum exact test. **F**) Dose response assays of the indicated human SCLC cell lines (NCI-H1048, NCI-H446, and NCI-H69) and human NSCLC cell lines (A549, HCC4006, and NCI-H1299) treated for 6 days with increasing doses of the cyclin A/B RxL inhibitor (CIRc-004). Average half-maximal effective concentration (EC50) for cyclin A/B RxL inhibitor (CIRc-004) shown next to respective cell line. **G**) Representative flow cytometry analysis of cleaved PARP in NCI-H1048 cells treated with CIRc-004 (200 nM) or DMSO (vehicle) for 3 days. **H**) Quantitation of cleaved PARP positive cells analyzed by flow cytometric analysis in the different SCLC and NSCLC lung cancer cell lines treated for 3 days with the indicated doses of the cyclin A/B RxL inhibitor (CIRc-004). **I**) Cell cycle distribution of the indicated SCLC and NSCLC cell lines treated with CIRc-004 (200 nM) or DMSO (vehicle) for 24 hours and then stained with propidium iodide (PI). For **F, H** and **I**, n=3 biological independent experiments and data are mean +/− SD. Arrow in **F** indicates DMSO-treated sample which was used for normalization. Statistical significance for **H** was calculated using unpaired, two-tailed students *t*-test. Where indicated, *=p<0,05, **=p<0.01, ***=p<0.001, ****=p<0.000.

Prior work demonstrated that cyclin RxL peptides could selectively induce apoptosis in transformed cell lines through hyperactivation of E2F^26^. Given the near universal loss of *RB1* leading to E2F dysregulation in SCLC^2–4,49^, we first tested these compounds for their anti-proliferative effects on a limited panel of SCLC cell lines. Compounds that inhibited both cyclin A and cyclin B had the strongest anti-proliferative activity (Fig. S1D), while the inactive enantiomer of the dual cyclin A/B RxL macrocycle CIRc-004, named CIRc-005, lacked both biochemical activity and cellular potency (Figs. 1B, S1D). Selective macrocyclic RxL site binders for either cyclin A or cyclin B were significantly less potent, and the activity on cyclin E was dispensable for the anti-proliferative effects in these cell lines (Figs. 1B, S1D). The non-transformed fibroblast cell line WI-38 was insensitive to cyclin A/B RxL inhibitors suggesting therapeutical potential for this novel class of macrocycles. Thus, we refer to this new class of cancer drugs as cyclin A/B RxL inhibitors.

### Small Cell Lung Cancer Cell Lines with High E2F Activity are Hypersensitive to Cyclin A/B RxL Macrocyclic Peptide Inhibitors

Given previous data^26^ and our data above, we hypothesized that cancers with high E2F activity, such as SCLC with near universal LOF *RB1* and *TP53* mutations^2–4,49^, would be hypersensitive to cyclin A/B RxL inhibitors. Therefore, we performed a screen of 46 SCLC cell lines with a cyclin A/B dual RxL inhibitor CIRc-004 (Fig. 1D). Strikingly, almost all SCLC cell lines were highly sensitive to the CIRc-004 at low nanomolar concentrations. Gene Set Variation Analysis (GSVA)^50^ using MSigDb hallmark gene sets^51^ of RNA-seq data from these SCLC cell lines showed high expression of E2F targets and G2/M checkpoint pathway scores correlated with CIRc-004 hypersensitivity (Fig. 1E). CIRc-004 sensitivity was observed irrespective of SCLC subtype^52^ (Fig. 1D). To identify which other cancer cell types were sensitive to cyclin A/B RxL inhibitors and to ensure that cyclin A/B RxL inhibitors were not generally toxic, we also screened the Horizon Discovery OncoSignature^TM^ cell line panel (Fig. S2A), which includes 300 cancer cell lines, with an early generation RxL inhibitor that broadly inhibits cyclins A, B, and E (CIRc-001) (Fig. 1B, S1D). ∼180 out of 300 cancer cell lines were sensitive to CIRc-001 with EC50’s<1 μM (Fig. S2A) which demonstrated cytotoxic activity in other cancer types, but also suggested selective sensitivity as ∼40% of cell lines were inherently resistant to CIRc-001 even at high micromolar concentrations. Lung, breast, and ovarian cancer lines had a relatively large number of sensitive cell lines.

Low throughput dose-response assays in 3 SCLC (NCI-H1048, NCI-H446, and NCI-H69), and 3 NSCLC (A549, HCC4006, and NCI-H1299) that we identified as inherently resistant to CIRc-004, showed that SCLC lines were hypersensitive to CIRc-004, but not its inactive enantiomer CIRc-005, all with EC50s <20 nM (Figs. 1F, S2B). Consistent with this, cyclin A/B RxL inhibition induced apoptosis in these SCLCs, but not in these NSCLCs (Fig. 1G-H). Cyclin A/B RxL inhibition induced a mitotic arrest in a dose-dependent manner (Fig. 1I, S2C). These phenotypes required both cyclin A and cyclin B RxL inhibitory activity as cyclin A selective (CIRc-018) or cyclin B selective (CIRc-019) RxL inhibitors did not affect proliferation, apoptosis, or cell-cycle distribution (Fig. S2D-F, S1D).

### Genome-Wide CRISPR/Cas9 Knockout Screen Uncovers Mechanisms by which Cyclin A/B RxL Inhibitors Induce Apoptosis

To identify downstream mechanisms by which cyclin A/B RxL inhibitors block proliferation and induce apoptosis, we performed genome-wide CRISPR/Cas9 knockout resistance screens in the highly sensitive NCI-H1048 SCLC cell line. For these screens, NCI-H1048 cells were first infected with the genome-wide Brunello sgRNA library (CP0043) containing 77,741 sgRNAs and 1000 control sgRNAs at an MOI of 0.3 (see methods). On day 12 [early timepoint (ETP)], all cells were pooled and split into 4 treatment arms: 1) cyclin A/B RxL inhibitor (CIRc-004); 2) cyclin A/B/E RxL inhibitor (CIRc-001); 3) a highly selective orthosteric Cdk2 inhibitor in clinical trials (PF-07104091); and 4) the inactive enantiomer of CIRc-004 (CIRc-005) as a negative control and comparator (Fig. 2A). Active compounds were used at their ∼EC90 concentrations in NCI-H1048 cells (Fig. S3A-C). The orthosteric Cdk2 inhibitor PF-07104091 was included as a comparison as preclinical data with PF-07104091 revealed that SCLC cell lines were also highly sensitive to PF-07104091 leading to an early phase clinical trial of PF-07104091 in patients with SCLC (NCT04553133)^53^. Once resistant populations of cells emerged [day 26=late timepoint (LTP)], cells were harvested for deep sequencing.

**Figure 2:**
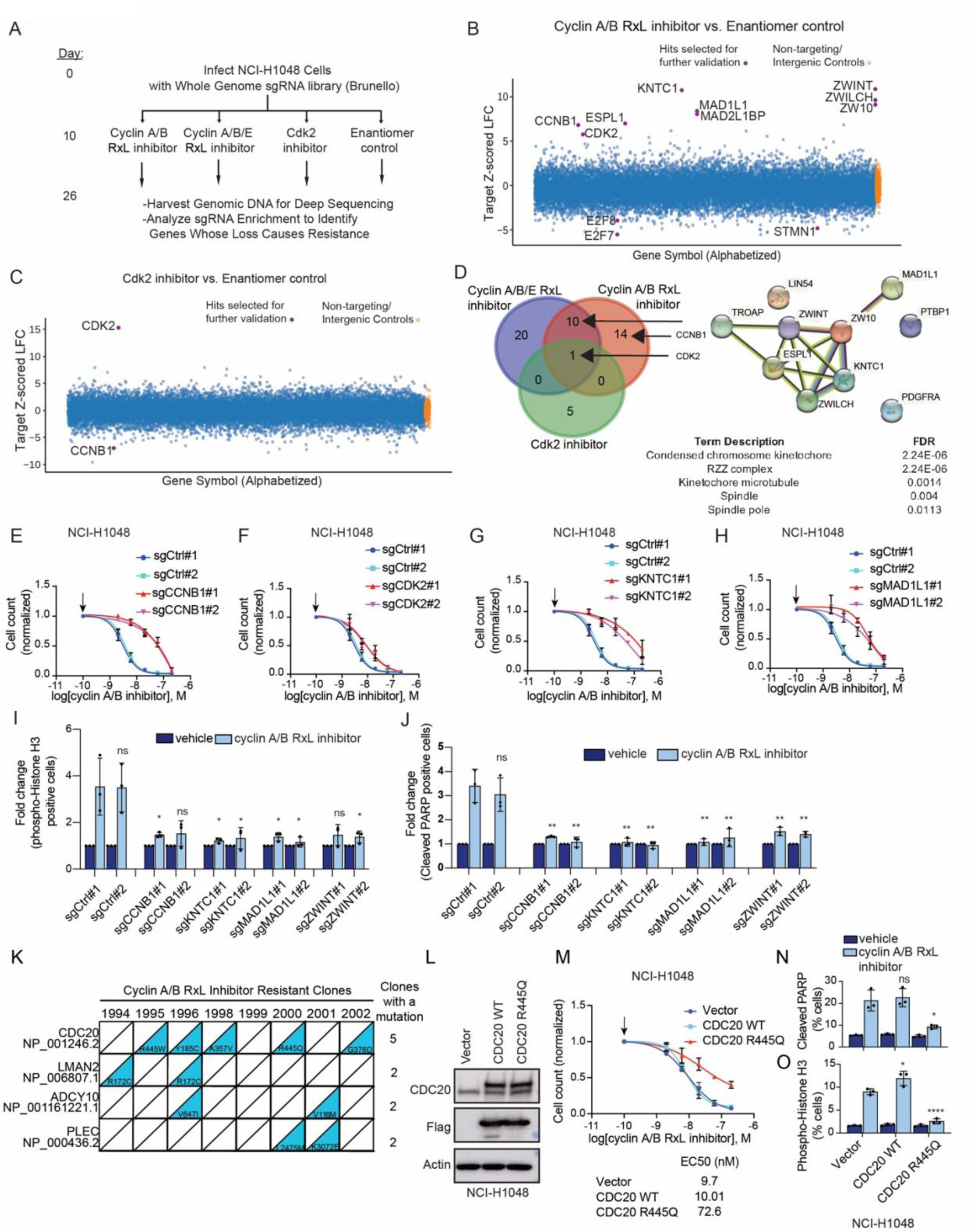
A Genome-wide CRISPR/Cas9 Knockout Screen and Forward Genetic Screen Identifies How Cyclin A/B RxL Inhibitors Kill Cancer Cells. **A**) Schema for the genome-wide CRISPR/Cas9 resistance screen in NCI-H1048 cells infected with the Brunello genome-wide sgRNA library. Following selection (day 10), cells were treated with cyclin A/B RxL inhibitor (CIRc-004, 200 nM); cyclin A/B/E inhibitor (CIRc-001, 200 nM); Cdk2 inhibitor (PF-07104091, 500 nM), or the inactivate enantiomer of CIRc-004 (CIRc-005, 200 nM) used as a negative control and then harvested at day 26 when the sgRNA-infected cells showed resistance. **B-C**) Top enriched and depleted hits determined by Apron analysis of the **B**) cyclin A/B RxL inhibitor (CIRc-004), or the **C**) Cdk2 (PF-07104091) at the end timepoint (day 26) all relative to the CIRc-005 enantiomer negative control at the end timepoint (day 26). For **B-C,** n=2 biological independent experiments. **D**) Venn diagram showing top enriched hits (q-value<0.25) among the different treatment groups indicated. Shown next to it are the protein-protein interaction network analysis (http://string-db.org/) of statistically significant hits shared between cyclin A/B, cyclin A/B/E RxL and Cdk2 inhibitors linked to the biological processes indicated and all involve in the spindle assembly checkpoint in mitosis. **E-H**) Dose response assays of NCI-H1048 cells infected with two independent non-targeting sgRNAs (sgCtrl) or two independent sgRNAs against **E**) *CCNB1*, **F**) *CDK2*, **G**) *KNTC1*, **H**) *MAD1L1*, treated for 6 days with increasing doses of CIRc-004. **I**) Quantitation of phospho-histone H3 flow cytometric data of NCI-H1048 cells infected with the sgRNAs indicated and treated with CIRc-004 at 20 nM for 24 hours. Data is plotted as fold change after cyclin A/B RxL inhibitor treatment relative to vehicle. **J**) Quantitation of cleaved PARP flow cytometric data of NCI-H1048 cells infected with the sgRNAs indicated and treated with CIRc-004 at 20 nM for 3 days. Data is plotted as fold change after cyclin A/B RxL inhibitor treatment relative to vehicle. **K**) Genes with mutations in more than one out of eight CIRc-004 resistant clones from the forward genetic screen in iHCT-116 cells (see Methods). **L**) Immunoblot of NCI-H1048 cells stably expressing vector, Flag-CDC20 WT, or Flag-CDC20 R445Q mutant. **M**) Dose response assay of NCI-H1048 cells from **L** treated with increasing doses of CIRc-004. **N**) Quantitation of cleaved PARP flow cytometric data of NCI-H1048 cells from **L** treated with CIRc-004 at 20 nM for 3 days. **O**) Quantitation of phospho-histone H3 flow cytometric data of NCI-H1048 cells from **L** treated with CIRc-004 at 20 nM for 24 hours. For **E-J**, **L-O**, n=3 biological independent experiments and data are mean +/– SD. Arrows in **E-H** indicates DMSO-treated sample which was used for normalization. Statistical significance in **E-H**, **N-O** was calculated using unpaired, two-tailed students *t*-test. Where indicated, *=p<0,05, **=p<0.01, ***=p<0.001, ****=p<0.0001.

As expected for a positive selection screen, sgRNA enrichment/depletion comparing biological replicates showed that the 2 biological replicates were modestly correlated with robust enrichment of some sgRNAs in both biological replicates for each drug condition (Fig. S3D-G). Gene level analysis comparing the drug treatment LTP vs. inactive enantiomer LTP or the shared ETP identified several highly significant “enriched hits” whose CRISPR inactivation caused resistance to each inhibitor. Enriched hits after treatment with cyclin A/B or A/B/E RxL inhibitors vs. the orthosteric Cdk2 inhibitor were surprisingly largely non-overlapping, highlighting a distinct mechanism of cell killing by cyclin A/B or cyclin A/B/E RxL inhibitors compared to orthosteric Cdk2 inhibitors (Fig. 2B-D, S3H-K). Cyclin B itself (*CCNB1*) was a top enriched hit after treatment with the cyclin A/B RxL inhibitor suggesting that cyclin B is necessary for cell killing by the cyclin A/B RxL inhibitor (Fig. 2B, 2D, S3I). While this provided evidence to confirm cyclin B being a likely target of cyclin A/B RxL inhibitors, it was also surprising and suggested that cyclin A/B RxL inhibitors act to promote a gain of cyclin B function to kill cells. The only common hit whose loss caused resistance to all 3 compounds was Cdk2 itself, which also supports Cdk2 as a common drug target for all 3 compounds as Cdk2 is the canonical kinase for cyclin A/E and the drug target for the Cdk2 inhibitor (Fig. 2B-C, S3H-K), but also suggested that cell killing may involve a gain of Cdk2 function.

Notably, many of the enriched hits in both cyclin A/B and cyclin A/B/E RxL inhibitors were involved in the spindle assembly checkpoint (SAC) during mitosis including *KNTC1*, *ESPL1*, *ZWINT*, *ZWILCH*, *ZW10*, and *MAD1L1* (Fig. 2B-C, S3H-K) demonstrating that a functional SAC complex is necessary for cell killing by cyclin A/B or A/B/E RxL inhibitors, but not by the Cdk2 inhibitor. On the other hand, repressive E2Fs, including E2F7 and E2F8, that bind and repress genes bound by activating E2Fs (E2F1-E2F3) to oppose E2F1 activity^54–57^, were among the top depleted hits that further sensitized cells to cyclin A/B or A/B/E RxL inhibition (Fig. 2B, S3H). The mitotic phospho-protein stathmin (*STMN1*)^58^ was also a top depleted hit after cyclin A/B or A/B/E RxL inhibition suggesting that *STMN1* inactivation sensitizes cells to cyclin A/B inhibition (Fig. 2B, S3H). Lastly, *CCNB1* was a top depleted hit in the orthosteric Cdk2 inhibitor screen (Fig. 2C) suggesting that cyclin B inhibition potentiates cellular killing by orthosteric Cdk2 inhibitors, which is achieved through our dual cyclin A/B RxL inhibitors.

For further validation studies, we focused on the cyclin A/B RxL inhibitor (CIRc-004) as CIRc-004 spared cyclin E and, perhaps as a result, demonstrated modestly increased cellular potency (Fig. S1D). CRISPR inactivation of *CCNB1, CDK2, KNTC1, LIN54, MAD1L1,* and *ZWINT* in NCI-H1048 cells caused robust resistance to CIRc-004 in dose-response assays thereby validating our screen results (Fig. 2E-H, S4A-B). CRISPR inactivation of select validated hits (*CCNB1, CDK2, KNTC1,* and *MAD1L1*) in NCI-H446 cells also caused CIRc-004 resistance suggesting a conserved mechanism of cell killing (Fig. S4C-F). Moreover, *CCNB1* inactivation increased sensitivity, while *CDK2* inactivation caused partial resistance to the orthosteric Cdk2 inhibitor (Fig. S4G-H), thereby validating our Cdk2 inhibitor screen. Lastly, CRISPR inactivation of *CCNB1*, *KNTC1, MAD1L1,* or *ZWINT* rescued the mitotic arrest and apoptosis induced by CIRc-004 (Fig. 2I-J, S4I-K).

### Forward Genetic Screen Identifies CDC20 Mutants that Cause Resistance to the Cyclin A/B RxL Inhibitor

The Nijhawan lab recently described a forward genetic system using an engineered colorectal cancer cell line, termed iHCT116, that identifies mutations that confer compound resistance using HCT116 cells engineered for inducible protein degradation [with the addition of indole acetic acid (IAA)] of mismatch repair protein, MLH1^59^. As a result, cells cultured without IAA have a lower mutation rate (Mut-low) than cells cultured with IAA (Mut-high). These cells are diploid and, therefore, resistance mutations are more likely to be the result of a gain of function, which is distinct from the CRISPR knockout screen above. CIRc-004 inhibited proliferation of iHCT116 cells over three days (IC50 = 0.057 μM) (Fig. S5A). After barcoded iHCT116 cells were cultured for an extended period, six and twelve clones from Mut-low and Mut-high conditions, respectively, were isolated (Fig. S5B-C), and the IC50 for all clones were at least 100-fold higher than parental cells. None of these clones tested were resistant to an unrelated toxin, MLN4924, suggesting that non-specific mechanisms were less likely explanations for CIRc-004 resistance (Fig. S5D). After evaluating the barcode sequence in all 18 clones, we found 5 and 12 founders in the Mut-low and Mut-high condition, respectively (Fig. S5E). The increased number in the Mut-high condition provides evidence of genetic resistance. 8 distinct clones were selected for exome sequencing. Only four genes were mutated in more than 1 of 8 clones. Strikingly, heterozygous *CDC20* mutations in 5 of 8 clones (Fig. 2K) and two of the five CDC20 mutations affected arginine 445. Moreover, arginine 445 and the other three altered amino acids (tyrosine 185, alanine 357, and glycine 376) all mapped to the same surface of CDC20 in the crystal structure where CDC20 interacts with the mitotic checkpoint complex (MCC) to promote SAC activation^60–63^ (Fig. S5F). Hence, these heterozygous CDC20 mutations are likely dominant negative mutations allowing cells to bypass the MCC and SAC and transition to anaphase in the face of CIRc-004. Ectopic expression of CDC20 R445Q, but not CDC20 WT, at levels similar to endogenous CDC20 in iHCT116 cells and NCI-H1048 SCLC cells led to CIRc-004 resistance (Fig. 2L-O, S5G-H) which validated our screen and demonstrated that our screen results extended to SCLC. Together our CRISPR knockout screen and gain of function forward genetic screen both converge on SAC as the downstream mechanism of cell killing by cyclin A/B RxL inhibition.

### Spindle Assembly Checkpoint (SAC) Activation by Mps1 is Required for Cell Killing by Cyclin A/B RxL Inhibitors

To better elucidate the mechanism by which CIRc-004 causes mitotic arrest and apoptosis, we next performed fluorescent live cell imaging in NCI-H1048 cells stably expressing GFP-H2B to visualize mitosis. These experiments revealed that cells failed to initiate anaphase after treatment with CIRc-004 consistent with SAC activation causing mitotic cell death (Fig. 3A-B). SAC protein components assemble upon the scaffold protein KNL1, which is phosphorylated by the mitotic kinase Mps1 in early mitosis^64^. Because KNL1 is dephosphorylated upon satisfaction of SAC, persistence of KNL1 phosphorylation can be used as a marker of SAC activation. Consistent with this, CIRc-004 robustly increased phospho-KNL1 in sensitive SCLC cell lines but not in insensitive NSCLC cell lines at the same concentration that it induced apoptosis (Fig. 3C, Fig. S6A-D, also see Fig. 1). Based on these results, we hypothesized that chemical inhibition of Mps1 activity would phenocopy genetic loss of *KNTC1, MAD1L1,* or *ZWINT* and induce CIRc-004 resistance. We first measured the anti-proliferative impact of a highly selective Mps1 inhibitor (BAY-1217389) in order to pinpoint a sublethal Mps1 inhibitor concentration (3 nM) that blocked KNL1 phosphorylation without having a significant effect on proliferation so that cell cycle or cytotoxicity changes were unlikely to confound our results (Fig. 3D, S6E-F). Mps1 inhibition rescued NCI-H1048, NCI-H69, and NCI-H446 SCLC cells from CIRc-004-induced SAC activation, proliferation defects, and apoptosis (Fig. 3D-F, Fig. S6G-J). Moreover, exogenous expression of CDC20 R445Q, but not CDC20 WT, completely abolished SAC activation induced by CIRc-004 in both iHCT116 cells and NCI-H1048 cells (Fig. S6K-L). Together these results demonstrate that cyclin A/B RxL inhibitors robustly promote SAC activation which is required for their ability to induce apoptosis.

**Figure 3:**
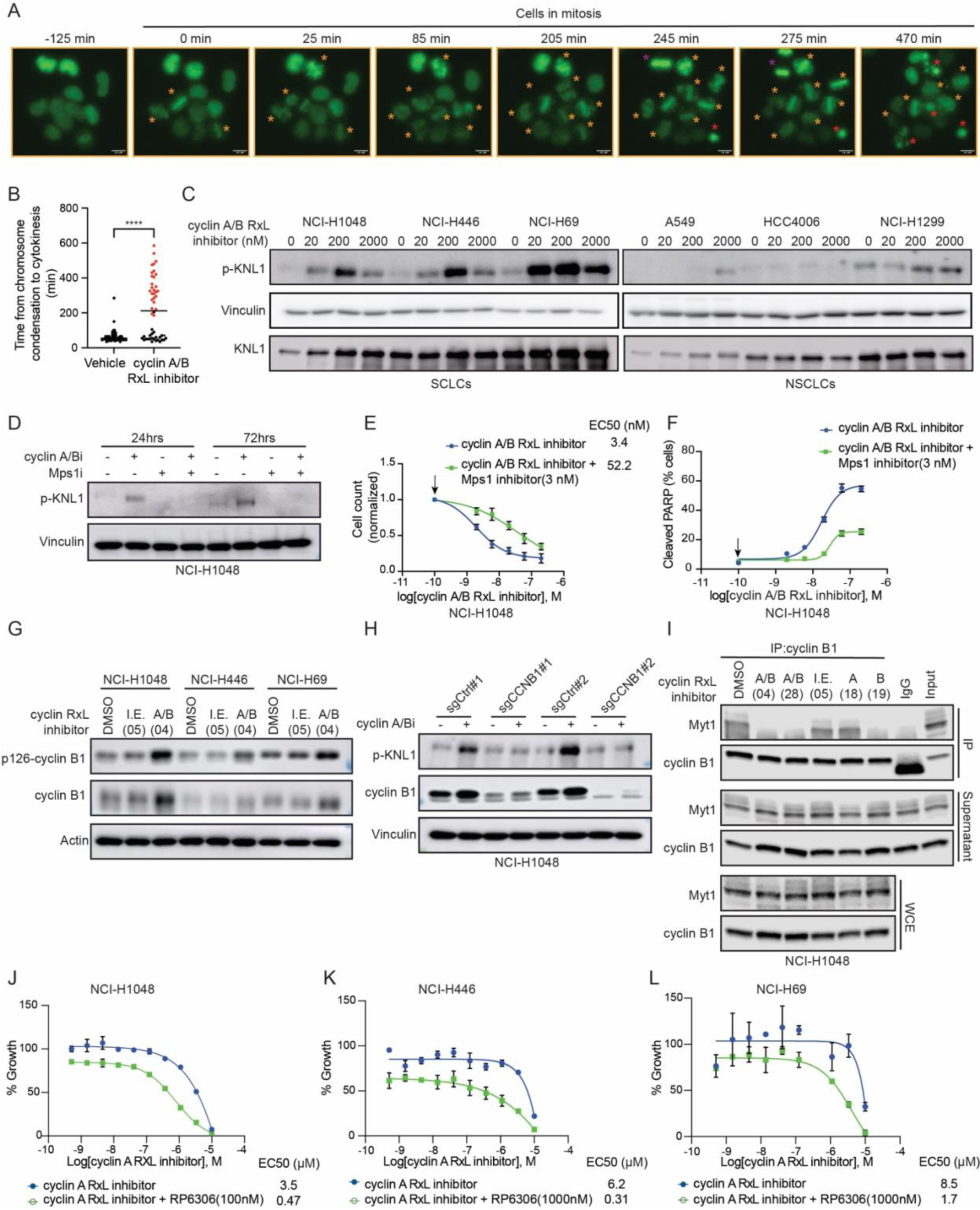
Spindle Assembly Checkpoint (SAC) Activation by Cyclin B or Mps1 is Necessary for Cyclin A/B RxL Inhibitors to Induce Apoptosis in SCLC Cell Lines. **A**) Representative images from time-lapse fluorescent microscopy of NCI-H1048 cells expressing H2B-GFP treated with 20 nM cyclin A/B RxL inhibitor (CIRc-004) over 10 hours (yellow star: arrested mitotic nuclei, purple star: normal mitotic nuclei, red star: apoptotic nuclei). Magnification=20x, scale bar 10 µm. **B)** Dot plot showing time (mins) taken by vehicle or drug treated cells to go from chromosome condensation to cytokinesis completion. Cells that failed to complete cytokinesis by experimental end point are shown in red. n=50 total mitotic nuclei per condition were counted from 2 different experiments. Statistical significance was calculated using unpaired, two-tailed students *t*-test. *=p<0.05, **=p<0.01, ***=p<0.001, ****=p<0.0001. **C**) Immunoblot analysis of the indicated human SCLC cell lines (NCI-H1048, NCI-H446, and NCI-H69) and insensitive human NSCLC cell lines (A549, HCC4006, and NCI-H1299) treated for 24 hours with increasing doses of the CIRc-004. n=3 biological independent experiments. **D)** Immunoblot analysis of NCI-H1048 cells treated with CIRc-004 at 20 nM, the Mps1 inhibitor (BAY-1217389) at 3 nM, combination of CIRc-004 and Mps1 inhibitor, or DMSO for 24 hours. **E**) Dose response assays of NCI-H1048 cells and **F**) Non-linear regression curves of cleaved PARP FACS analysis of NCI-H1048 cells treated with increasing concentrations of CIRc-004 in presence or absence of the Mps1 inhibitor (BAY-1217389) at 3 nM for 3 days. Data are mean +/− SD. n=3 biological independent experiments. **G**) Immunoblot analysis for active cyclin B using the phospho-specific antibody cyclin B phosphoserine 126 of the SCLC cell lines indicated treated with CIRc-004 for 4 hours. **H)** Immunoblot analysis of NCI-H1048 cells infected with 2 independent sgRNAs targeting *CCNB1* or 2 independent non-targeting controls (sgCtrl) treated with CIRc-004 at 20 nM or vehicle for 24 hours. **I**) Immunoblot analysis after cyclin B1 immunoprecipitation in NCI-H1048 cells treated with cyclin A/B RxL inhibitor (CIRc-004), cyclin A/B RxL inhibitor (CIRc-028), inactive enantiomer of CIRc-004 (I.E., CIRc-005), cyclin A RxL inhibitor (CIRc-018), cyclin B RxL inhibitor (CIRc-019) or DMSO. All inhibitors were used at 300 nM for 2 hours. n=3 biological independent experiments for CIRc-004 and CIRc-005, n=2 for all other inhibitors. Dose response assay of **J**) NCI-H1048, **K**) NCI-H446 and **L**) NCI-H69 cells treated with cyclin A RxL inhibitor (CIRc-018) in presence or absence of RP-6306 (Myt1 inhibitor) for 5 days. Data are mean +/− SD from two technical replicates and n=3 biological independent experiment. Arrows in **E-F** indicates DMSO-treated sample which was used for normalization.

### Cyclin B RxL Macrocycles Block an Inhibitory Cyclin B:Myt1 Interaction Thereby Increasing Cyclin B-Induced SAC Activation

Cyclin B (*CCNB1*) was a top enriched hit in our CIRc-004 resistance screen suggesting that cyclin B activity is necessary for CIRc-004’s mode of action (see Fig. 2). This is paradoxical if CIRc-004 simply blocks cyclin B function and suggests that CIRc-004 may lead to a gain, rather than a loss, of cyclin B activity. Consistent with this and our results above, increased cyclin B-Cdk1 activity itself can activate SAC^62,65^. Indeed, we observed that cyclin B activity as measured by phosphorylation of serine 126 on cyclin B^66,67^ was increased with CIRc-004 treatment (Fig. 3G). Moreover, NCI-H1048 *CCNB1* knockout cells failed to activate SAC in response to CIRc-004 demonstrating that cyclin B activity is necessary for SAC activation (Fig. 3H). We therefore hypothesized that CIRc-004 blocks a direct cyclin B RxL substrate interaction that normally functions as a negative regulator of cyclin B-Cdk1 activity. One such direct cyclin B RxL substrate is the kinase Myt1 (gene=*PKMYT1*), which acts as a negative regulator of cyclin B-Cdk1 activity by directly phosphorylating Cdk1 at threonine 14 to inhibit cyclin B/Cdk1 activity^68^. IP/Co-IP experiments for cyclin B and Myt1 in the presence of RxL inhibitors found that the interaction of cyclin B and Myt1 was disrupted by dual cyclin A/B RxL inhibitors or a selective cyclin B RxL inhibitor, but not a selective cyclin A RxL inhibitor or the inactive enantiomer of CIRc-004 (Fig. 3I). Moreover, the Myt1 inhibitor RP-6306^69,70^ potentiated cell killing by the selective cyclin A RxL inhibitor in NCI-H1048, NCI-H446, and NCI-H69 SCLC cell lines (Fig. 3J-L). This was selective for the cyclin A RxL inhibitor as RP-6306 did not potentiate cell killing by cyclin B- or cyclin A/B- RxL inhibitors (Fig. S7A-I). Together these data demonstrate that the cyclin B-Myt1 interaction is disrupted by cyclin A/B RxL inhibitors, leading to Cdk activation on the cyclin B complex to promote SAC activation and mitotic cell death.

### Cyclin A/B RxL Inhibitors Induce a Neomorphic Cyclin B-Cdk2 Interaction that Promotes Mitotic Cell Death

To further investigate the mechanism by which CIRc-004 induces mitotic cell death, we performed a CRISPR/Cas9 base editor screen in NCI-H1048 cells using both A>G and C>T Cas9 base editors^71^ with a custom designed sgRNA library that included sgRNAs tiling *CCNB1*, *CCNA2*, *CDK2*, *CDC20* (Fig. 4A, also see methods). Our sgRNA library also included positive control sgRNAs targeting splice sites of essential genes and non-targeting negative controls. At the ETP (day 10), cells transduced with the sgRNA library were treated with the cyclin A/B RxL inhibitor (CIRc-004) or the enantiomer negative control (CIRc-005) for 16 days (LTP=day 26), and then deep sequencing for the sgRNA barcodes was performed (Fig. 4A). sgRNA enrichment/depletion comparing biological replicates showed that all biological replicates were strongly correlated (Fig. S8A). Moreover, analysis of CIRc-005 at the LTP vs. the plasmid DNA showed strong depletion of positive controls relative to negative controls (Fig. S8B) demonstrating technical success. To identify candidate mutations that caused CIRc-004 resistance, sgRNA enrichment/depletion comparing CIRc-004 LTP vs. CIRc-005 LTP at *CCNB1*, *CCNA2*, *CDK2*, and *CDC20 loci* was analyzed. Consistent with our forward genetic screen in iHCT-116 cells (see Fig. 2K-O), we observed strong enrichment of several sgRNAs predicted to mutate CDC20 at amino acid (aa) residue 445 (Fig. S8C). Several sgRNAs targeting *CCNB1* were enriched including an sgRNA predicted to mutate Ala203Thr in the MRAIL motif where cyclin B interacts with RxL-containing substrates^24,25^ (Fig. 4B). Interestingly, there was a cluster of enriched sgRNAs predicted to mutate cyclin B aa’s 169-177 where it interacts with cyclin-dependent kinases, which is canonically Cdk1^72^ (Fig. 4B-C). Given these promising results and to directly confirm the specific mutations on *CCNB1* that cause CIRc-004 resistance, we then repeated the screen and performed amplicon sequencing on a focused region of *CCNB1* corresponding to aa’s 117-285. This directly validated several *CCNB1* mutants that conferred CIRc-004 resistance including Glu169Lys, Tyr170His, and Tyr177Cys (Fig. 4D). FACS-based competition assays showed that several of the candidate *CCNB1* mutants conferred CIRc-004 resistance (Fig. 4E). Amplicon sequencing of mixed populations of cells from the competition assays identified the precise base editor-induced *CCNB1* mutations and the fraction of cells carrying these mutations (Glu169Lys, Tyr170His, and Tyr177Cys) were dramatically enriched after CIRc-004 treatment (Fig. 4F).

**Figure 4:**
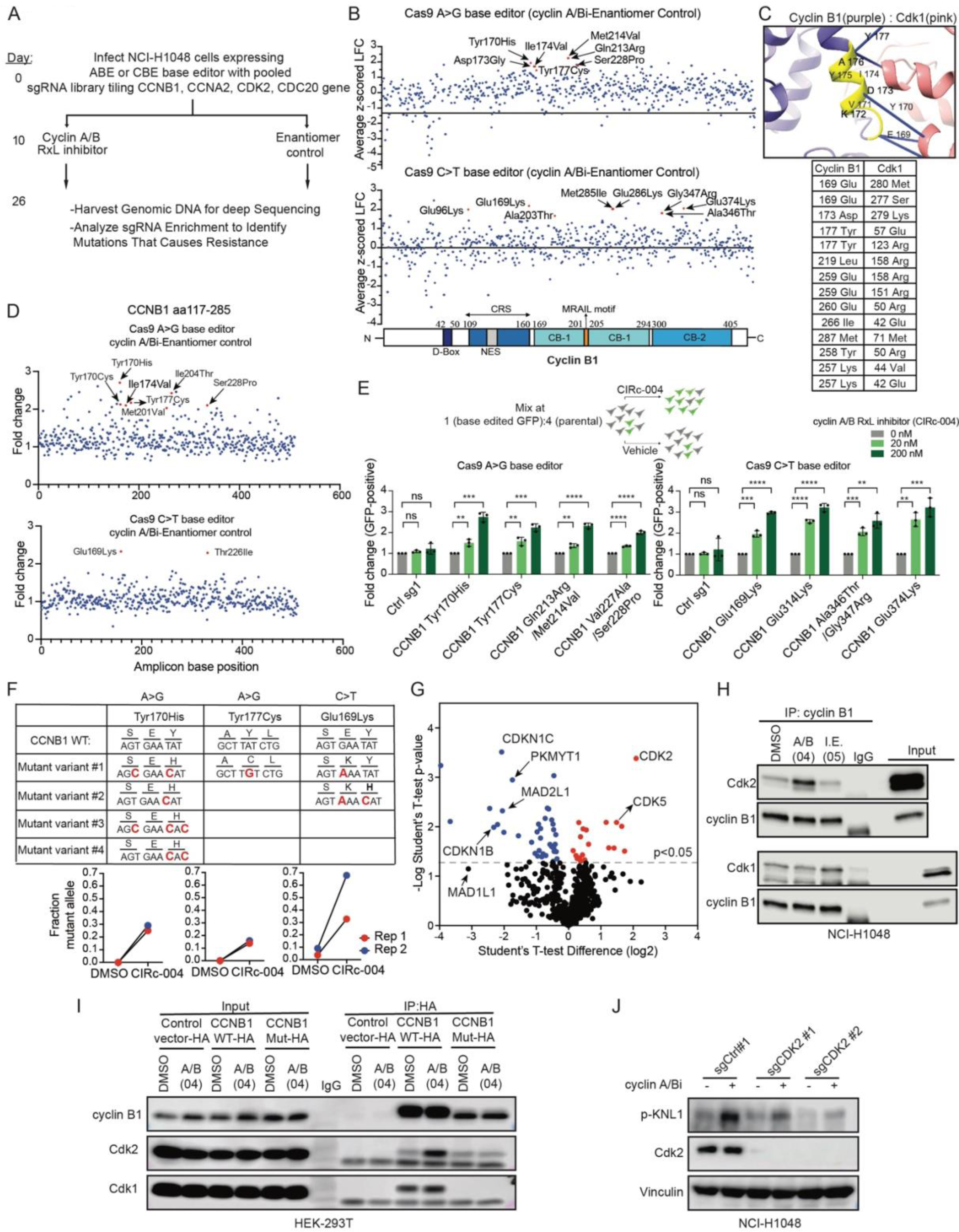
Cyclin A/B RxL Inhibitors Promote a Neomorphic Interaction Between Cyclin B and Cdk2. **A**) Schema for the base editor CRISPR/Cas9 cyclin A/B RxL inhibitor resistance screen in NCI-H1048 cells. Cells were infected with a custom-made sgRNA library including 2802 sgRNAs tiling *CCNB1*, *CCNA2*, *CDK2* and *CDC20*, and positive and negative control sgRNAs (see Methods). At day 10 [early timepoint (ETP)], cells were treated with the cyclin A/B RxL inhibitor (CIRc-004, 200 nM) or the inactivate enantiomer of CIRc-004 (CIRc-005, 200 nM) and then harvested at day 26 [late timepoint (LTP)]. **B**) Dot plot of average z-scored log-fold change (LFC) for each sgRNA tiling *CCNB1* in cells treated with CIRc-004 vs. CIRc-005 at day 26. The x-axis indicates the position of each sgRNA along *CCNB1* protein coding sequence. Variants of interest are labeled with the predicted amino acid position and mutational change. Top graph is the A>G and bottom graph is the C>T base editor screen. n=4 biological replicates. **C**) Predicted AlphaFold2 model of the cyclin B1:Cdk1 complex where high confidence interactions of cyclin B’s amino acids 169-177 with Cdk1 are highlighted in yellow. The table below lists the high confidence amino acid residues involved in the interaction between cyclin B1 and Cdk1. **D**) Dot plot of fold change variant reads at each nucleotide base position within *CCNB1* comparing NCI-H1048 Cas9 base-editor cells transduced with the sgRNA library in **A** treated with CIRc-004 vs. CIRc-005 at day 26. Variants of interest are labeled with the mutational change. n=2 biological replicates. **E**) FACS-based competition assay of GFP-positive NCI-H1048 Cas9 base-editor cells transduced with the indicated *CCNB1* sgRNA variants and GFP. GFP-positive *CCNB1* sgRNA variant cell lines were mixed with the parental base editor counterpart at a ∼1:4 ratio and then treated with CIRc-004 at the concentrations indicated for 13 days. Fold-change enrichment of GFP-positive cells comparing CIRc-004 vs. DMSO control is shown. Data are mean +/− SD and n=3 biological independent experiments. Statistical significance was calculated using unpaired, two-tailed students *t*-test. Where indicated, *=p<0,05, **=p<0.01, ***=p<0.001, ****=p<0.0001. **F**) Fraction of variant alleles from deep amplicon sequencing of genomic DNA from *CCNB1* Tyr170His, *CCNB1* Tyr177Cys, and *CCNB1* Glu169Lys cell lines from the competition assay in **E** at the experimental endpoint (Day 13) treated with CIRc-004 (200 nM) or DMSO. Top panel shows the common mutant variants observed for each sample. n=2 biological independent experiments. **G**) Volcano plot of IP of cyclin B followed by mass spectrometry (IP-MS) in NCI-H1048 cells first treated with CIRc-004 (50 nM) relative to CIRc-005 (50 nM) for 2 hours. n=3 independent experiments. **H**) Immunoblot analysis after IP of endogenous cyclin B1 in NCI-H1048 cells treated with CIRc-004 (300 nM), CIRc-005 (inactive enantiomer of CIRc-004, I.E), or DMSO for 2 hours. n=2 biological independent experiments. **I**) Immunoblot analysis after IP of exogenous cyclin B1 using an HA antibody in HEK-293T cells expressing CCNB1 WT-HA, CCNB1 triple mutant-(E169K/Y170H/Y177C)-HA, or a negative control vector, and treated with CIRc-004 (300 nM) or DMSO for 2 hours. n=3 biological independent experiments. **J**) Immunoblot analysis of NCI-H1048 cells infected with 2 independent sgRNAs targeting *CDK2* or a non-targeting control (sgCtrl) and treated with CIRc-004 at 20 nM or DMSO for 24 hours.

We then performed IP of Cyclin B1 followed by mass spectrometry (IP-MS) in NCI-H1048 cells treated with CIRc-004 or CIRc-005 as a control (Fig. 4G). Consistent with our experiments above (Fig. 3I), one of the top depleted proteins after CIRc-004 treatment was Myt1 (*PKMYT1*) (Fig. 4G, S9B) as well as known Cyclin/Cdk RxL-dependent inhibitors p27^Kip1^ (*CDKN1B*) and p57^Kip2^ (*CDKN1C*)^24^ (Fig. 4G). Surprisingly, one of the top enriched proteins after CIRc-004 treatment was Cdk2 (Fig. 4G, S9B). While cyclin B binds Cdk2 *in vitro*^73^, cyclin B canonically binds Cdk1^74^ and not Cdk2 in cells^72^. Alpha fold also identified high confidence interactions between cyclin B aa residues 169-177 with Cdk2 (Fig. S9A). IP experiments for cyclin B in NCI-H1048 and NCI-H446 cells confirmed that CIRc-004 increased the co-IP of Cdk2 without affecting Cdk1 (Fig. 4H, S9C-D). Moreover, CIRc-004 induced phosphorylation of a Cdk mitotic RxL-independent substrate stathmin (*STMN1*)^75–77^ and stathmin phosphorylation was abrogated by a selective Cdk2 inhibitor, but not a selective Cdk1 inhibitor (Fig. S9E-F). Stathmin phosphorylation inactivates stathmin disabling its ability to depolymerize microtubules which activates SAC^78^ and *STMN1* depletion strongly sensitized cells to CIRc-004 in our CRISPR knockout screen (Fig. 2B, S3H-J) together suggesting that CIRc-004-induced Cdk2-dependent stathmin phosphorylation promotes sensitivity to cyclin A/B RxL inhibitors. Based on our base editor screening results (Fig. 4B-F), we then tested whether a cyclin B triple mutant (Glu169Lys, Tyr170His, and Tyr177Cys) blocked the formation of CIRc-004-induced cyclin B-Cdk2 complexes. IP/Co-IP experiments showed that CIRc-004 induced an interaction between Cdk2 and cyclin B WT that was abrogated in cells expressing the cyclin B triple mutant (Fig. 4I, S9G). Moreover, Cdk2 inactivation in NCI-H1048 cells abrogated CIRc-004-induced SAC activation (Fig. 4J). These data, together with Cdk2 being a top enriched hit in our CRISPR knockout screen (Fig. 2B, S3H), demonstrate that cyclin A/B RxL inhibitors promote the redirection of Cdk2 to cyclin B to form cyclin B-Cdk2 complexes to promote SAC activation and mitotic cell death.

### Cyclin A RxL Peptides Block an Inhibitory Cyclin A:E2F1 Interaction Thereby Increasing E2F1 Activity Rendering Cells Hypersensitive to Cyclin A/B Inhibition

Our data above explains the functional contribution of cyclin B RxL inhibition in dual cyclin A/B RxL inhibitors mode of action (Fig. 3-4). We next sought to understand the role for cyclin A RxL inhibition in the mode of action of cyclin A/B dual RxL inhibitors. Cyclin A is expressed and canonically associates with Cdk2 in S-phase^79,80^. SAC is normally activated during mitosis when there is improper attachment of kinetochores to the mitotic spindles, which can be caused by DNA damage and replication stress that occurs during S-phase^62^. Consistent with this, pulse-labeling with EdU showed a large fraction of G2 cells undergoing active DNA synthesis after treatment with the cyclin A/B RxL inhibitor in NCI-H1048 and NCI-H446 cells (Fig. 5A). Moreover, phospho-γ-H2AX was markedly induced by cyclin A/B RxL inhibition in sensitive SCLC lines, but not in insensitive NSCLC cell lines (Fig. 5B, S10A), as was phospho-RPA2 levels (Fig. S10B). Cyclin A/B RxL inhibitors induced phospho-γ-H2AX irrespective of whether cells had the ability to form SAC complex and this required both cyclin A and cyclin B RxL inhibition (Fig. S10C-D).

**Figure 5:**
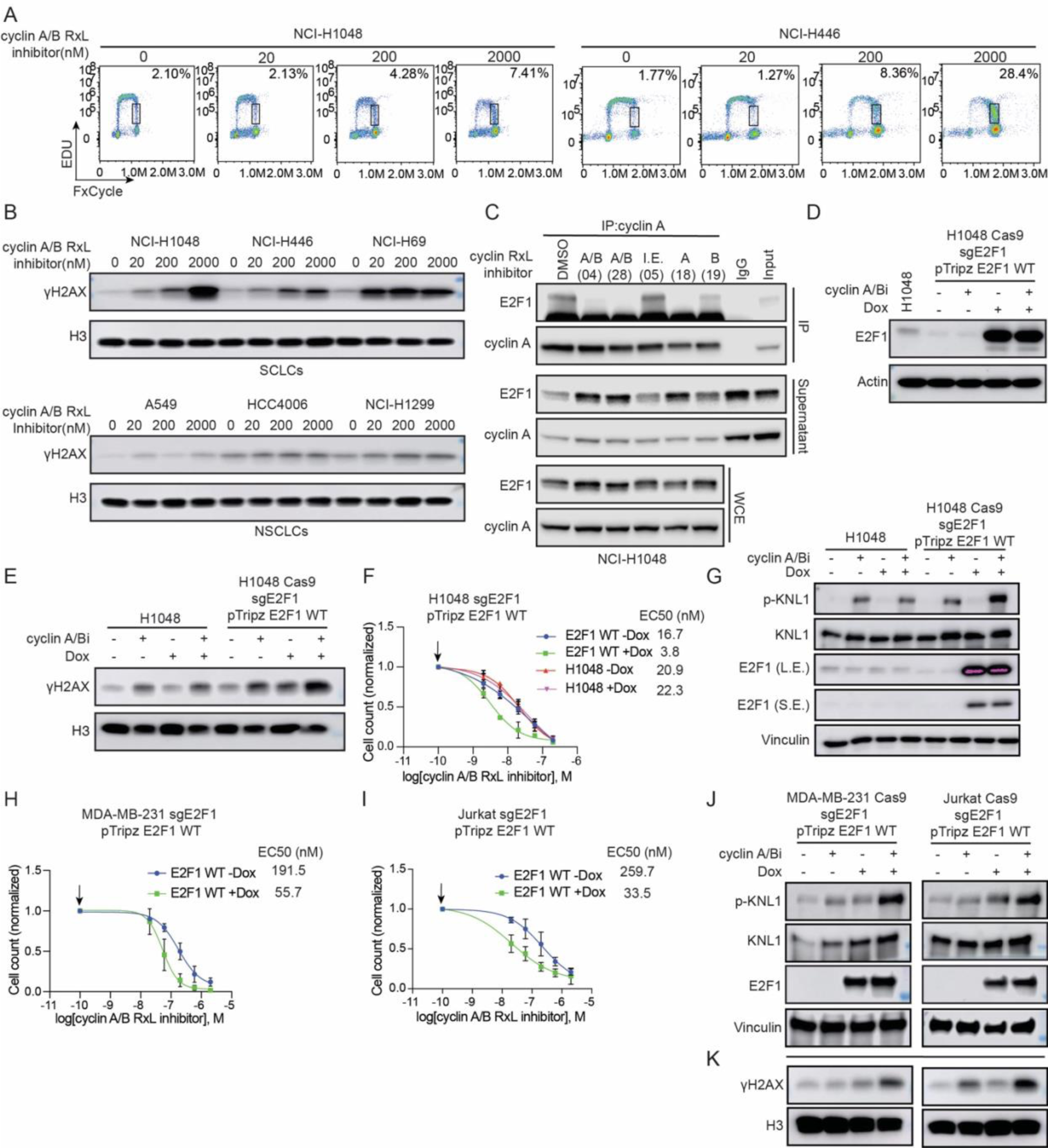
Cyclin A RxL Inhibition Leads to E2F1 Hyperactivation to Promote Sensitivity to Cyclin A/B RxL Inhibitors. **A**) FACS analysis of EdU and FxCycle in NCI-H1048 or NCI-H446 cells treated with increasing doses of cyclin A/B RxL inhibitor (CIRc-004) for 24 hours. EdU+ 4C DNA population shown for each panel in the top right corner. n=3 biological independent experiments **B**) Immunoblot analysis of the indicated human SCLC cell lines (NCI-H1048, NCI-H446, and NCI-H69) and insensitive human NSCLC cell lines (A549, HCC4006, and NCI-H1299) treated for 3 days with increasing doses of CIRc-004. n=3 biological independent experiments. **B)** Immunoblot analysis after cyclin A immunoprecipitation in NCI-H1048 cells treated with cyclin A/B RxL inhibitor (CIRc-004), cyclin A/B RxL inhibitor (CIRc-028), inactive enantiomer of CIRc-004 (I.E., CIRc-005), cyclin A RxL inhibitor (CIRc-018) cyclin B RxL inhibitor (CIRc-019) or DMSO. All inhibitors were dosed at 300 nM for 2 hours. n=3 biological independent experiments. **D, E, G**) Immunoblot analysis of NCI-H1048 cells infected with a doxycycline (DOX) inducible E2F1 sgRNA-resistant cDNA and then superinfected with an sgRNA targeted endogenous E2F1 grown in the presence or absence of DOX for 24 hours and then treated with CIRc-004 at 20 nM for another 24 hours (**D**, **G**), 72 hours (**E**). Dose response assays of **F**) NCI-H1048 **H**) MDA-MB-231 **I**) Jurkat cells grown in the presence or absence of DOX for 24 hours and then treated with increasing concentrations of CIRc-004 for 3 days (**F**) or 6 days (**H**, **I**). For **F, H, I**, data are mean +/− SD and arrows indicates DMSO-treated sample which was used for normalization. **J, K**) Immunoblot analysis of MDA-MB-231 and Jurkat cells generated as described in **D** and **E.** For **D**-**K**, n=3 biological independent experiments.

High E2F activity correlates with sensitivity to cyclin A/B RxL inhibitors in SCLC cell lines (see Fig. 1E). Repressive E2Fs, including E2F7 and E2F8, were top depleted hits that when inactivated sensitized cells to cyclin A/B RxL inhibitors in our CRISPR knockout screen (Fig. 2B, S3H) suggesting that heightened E2F activity could further enhance cyclin A/B RxL inhibitor sensitivity. Cyclin A binds E2F1 through an RxL motif, which leads to cyclin A-Cdk2-mediated inhibitory phosphorylation of E2F1 dampening E2F1 activity^21,22,24,81^ and blocking the cyclin A-E2F1 RxL interaction promotes apoptosis^15,16,26,82^. Therefore, we hypothesized that the dual cyclin A/B RxL inhibitor or the selective cyclin A RxL inhibitor blocks the cyclin A-E2F1 RxL interaction leading to heightened E2F1 activity. IP/Co-IP experiments for cyclin A and E2F1 in the presence of RxL inhibitors revealed that the interaction of cyclin A and E2F1 was only disrupted by dual cyclin A/B RxL inhibitors or a selective cyclin A RxL inhibitor (Fig. 5C). We then asked whether increasing E2F1 itself could further sensitize to cyclin A/B RxL inhibitors. Overexpression of E2F1 in E2F1 knockout NCI-H1048, MDA-MB-231, and Jurkat cells increased DNA damage, SAC activation, and enhanced sensitivity to cyclin A/B RxL inhibitors (Fig. 5D-K). Together these data demonstrate that the cyclin A: E2F1 RxL interaction is disrupted by cyclin A RxL inhibition leading to increased E2F1 activity, which increases sensitivity to cyclin A/B RxL inhibitors.

### Cyclin A/B RxL Inhibitors Have Anti-Tumor Activity in SCLC Xenograft Models

Next we performed *in vivo* studies first with a cyclin A/B RxL inhibitor CIRc-028 dosed intravenously. CIRc-028 was chosen as it has similar biochemical activity toward cyclin A/B and potency to that of CIRc-004 (Fig. 1B, S1D), but has better PK properties for use *in vivo*. Efficacy studies were performed in both NCI-H1048 and NCI-H69 xenografts treated with CIRc-028 dosed daily at 100 mg/kg for 14 days. Treatment with CIRc-028 induced tumor regressions in NCI-H69 xenografts and substantially inhibited tumor growth in NCI-H1048 xenografts (Fig. 6A-B) and was well tolerated without any weight loss (Fig. S11A-B). A pharmacodynamic (PD) study of NCI-H69 xenograft tumors from the efficacy study showed that CIRc-028 induced SAC activation and apoptosis *in vivo* (Fig. 6C-F, S11C).

**Figure 6:**
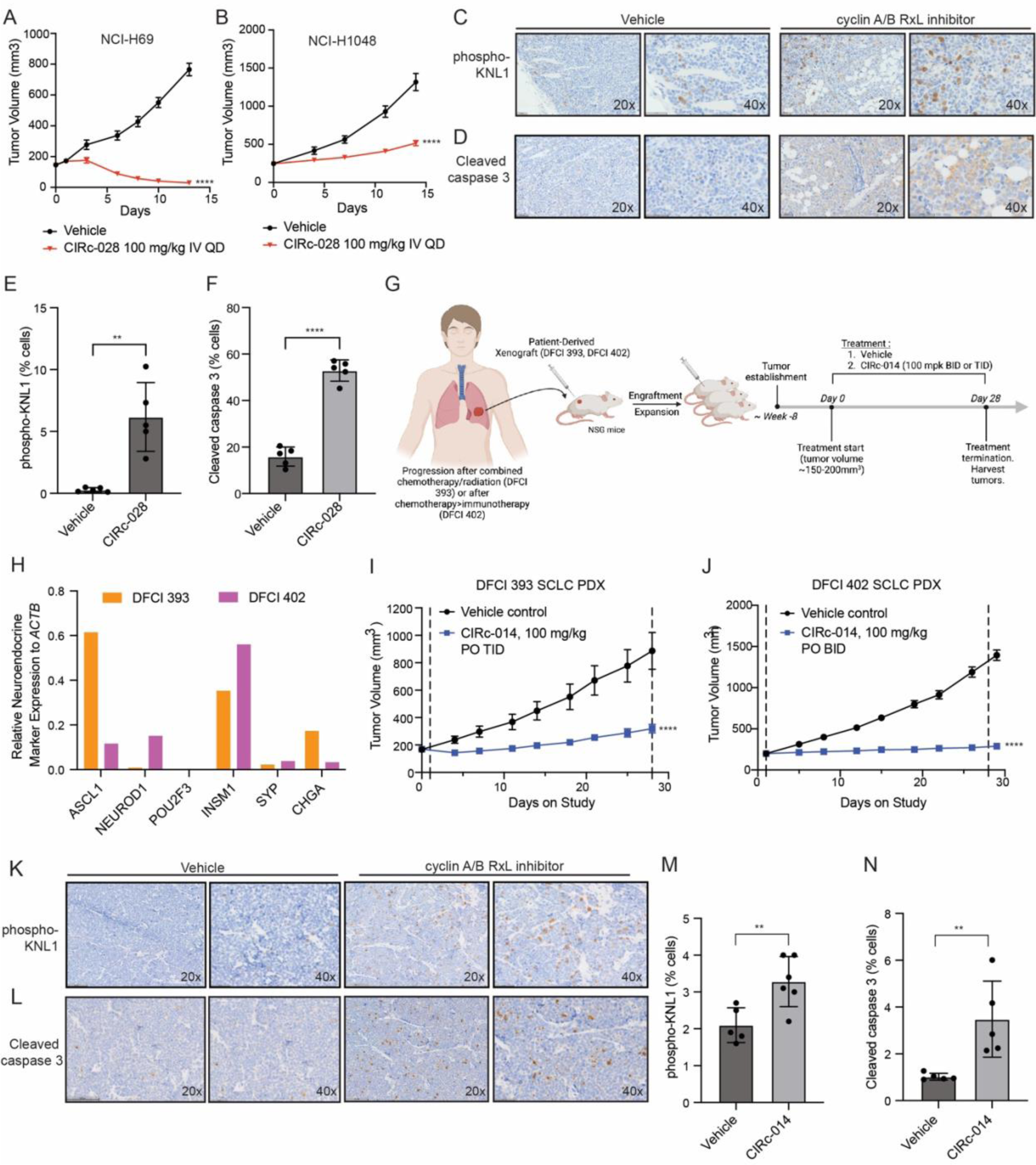
Cyclin A/B RxL Inhibitors Have On-Target and Robust Anti-Tumor Activity in SCLC Cell-Line Xenograft and Patient-Derived Xenograft Models. **A, B.** Tumor volume curves of NCI-H69 (**A**) or NCI-H1048 (**B**) cell-line xenografts in athymic nude mice treated with vehicle, cyclin A/B RxL inhibitor (CIRc-028, 100 mg/kg IV QD) for 14 days. For **A,** n=10 independent mice per arm. For **B**, n=10 independent mice for vehicle and n=8 independent mice for CIRc-028 arm. Data are mean +/− SEM. p<0.0001 by 2-way ANOVA for both NCI-H69 and NCI-H1048 comparing CIRc-028 vs. vehicle. IV=intravenous. QD=every day. **C, D)** Representative immunohistochemistry (IHC) images for phospho-KNL1 (**C**) and cleaved caspase 3 (**D**) of NCI-H69 xenografts treated with the cyclin A/B RxL inhibitor (CIRc-028) at 100 mg/kg/day by intravenous injection (IV) or vehicle for 14 days. Tumors were harvested 18 hours after the last dose. Magnification=20x or 40x as indicated, scale bar=50 µm. **E-F**) Quantification of phospho-KNL1 (**E**) and cleaved caspase 3 (**F**) IHC staining in tumors from C-D. For **E-F**, data are mean +/− SD. n=5 independent mice each with 1 tumor per mouse for each treatment group. Statistical significance was calculated using unpaired, two-tailed students *t*-test. *=p<0.05, **=p<0.01, ***=p<0.001, ****=p<0.0001. **G**) Schema for the SCLC DFCI 393 and DFCI 402 patient-derived xenograft (PDX) efficacy studies with CIRc-014 in NSG mice. **H**) mRNA expression of SCLC transcription factors and neuroendocrine markers relative to *ACTB* in DFCI 393 and DFCI 402 PDX models using FPKM values from RNA-seq data. **I-J**) Tumor volume curves of DFCI 393 (**I**) or DFCI 402 (**J**) PDX tumors in NSG mice treated with the orally bioavailable cyclin A/B RxL inhibitor CIRc-014 at 100 mg/kg PO TID (**I**) or 100 mg/kg PO BID (**J**) or vehicle continuously for 28 days. Dashed lines indicate start and end of treatment. PO=oral. For DFCI 393 PDX in **I**, n=10 independent mice for vehicle and n=9 independent mice for CIRc-014 arm. For DFCI 402 PDX in **J**, n=10 independent mice per arm. Data are mean +/− SEM. p<0.0001 by 2-way ANOVA for both DFCI 393 and DFCI 402 comparing CIRc-014 vs. vehicle. **K-N**) Representative IHC (**K,L**) and quantitation (**M,N**) for phospho-KNL1 (**K, M**) and cleaved caspase 3 (**L,N**) of DFCI 393 PDXs in NSG treated with CIRc-014 (100 mg/kg PO TID) or vehicle for 4 days. Tumors were harvested 8 hours after the last dose. Magnification=20x or 40x as indicated, scale bar=100 µm (20x) and 50 µm (40x). For **M-N**, data are mean +/− SD with 1 tumor per mouse for each treatment group. For **M**, n=5 independent mice for vehicle and n=6 independent mice for CIRc-014. For **N**, n=5 independent mice for vehicle and n=5 independent mice for CIRc-014. Statistical significance was calculated using unpaired, two-tailed students *t*-test. *=p<0.05, **=p<0.01, ***=p<0.001, ****=p<0.0001. Figure 6G was made using BioRender.

Lastly, we tested whether cyclin A/B RxL inhibition had efficacy in two patient-derived xenograft models (PDX) of SCLC (DFCI 393 and DFCI 402) derived from SCLC patients after each developed resistance to chemotherapy (Fig. 6G, also see methods). RNA-seq showed that DFCI 393 was an ASCL1+ model and DFCI 402 expressed both ASCL1 and NEUROD1 and both PDXs clustered with human neuroendocrine-high SCLC tumors^3^ (Fig. 6H, S11D). For these experiments, we used CIRc-014, which is a cyclin A/B RxL inhibitor that was further optimized for oral bioavailability (Fig. S1D, S11E). Strikingly, treatment with CIRc-014 caused near complete growth inhibition in both chemo-resistant SCLC PDX models (Fig. 6I-J, S11F) without any weight loss (Fig. S11G-H). Independent PD studies of DFCI 393 PDX tumor bearing mice treated with CIRc-014 for 4 days showed that CIRc-014 induced SAC activation and apoptosis *in vivo* (Figure 6K-N). Together these studies show on-target and robust anti-tumor activity of the orally bioavailable cyclin A/B RxL inhibitor CIRc-014 in chemo-resistant SCLC PDX models without overt toxicity.

## Discussion

Here we describe the development of cell-permeable macrocycles that bind cyclins to block their RxL-dependent interactions with RxL-containing substrates. We find that SCLCs with high E2F activity are highly sensitive to macrocycles that inhibit the ability of both cyclin A and cyclin B to interact with their corresponding RxL binding proteins leading to SAC activation and mitotic cell death. Two unbiased orthogonal genome-wide forward genetic screens in cell lines from different cancer types both show that cell killing by cyclin A/B RxL inhibitors is achieved through activation of SAC, which is distinct from the mechanism of cell killing by an orthosteric Cdk2 inhibitor PF-07104091 that is currently in a clinical trial for patients with SCLC (NCT04553133). Our data show that the efficacy of cyclin A/B RxL inhibitors is achieved by blocking the ability of cyclins to interact with at least 2 RxL-containing substrates including: 1) Disruption of a cyclin A-E2F1 inhibitory interaction which leads to increased E2F activity, priming cells for SAC-induced apoptosis; 2) Disruption of a cyclin B-Myt1 RxL inhibitory interaction which hyperactivates cyclin B-Cdk activity thereby increasing downstream SAC activation and mitotic cell death (see Fig. S12 for proposed mechanism). While disruption of these two RxL interactions is necessary for cell killing by cyclin A/B RxL inhibitors, there are almost certainly other cyclin-RxL-substrate interactions^24,25,83–85^ disrupted by these macrocycles that contribute to their ability to induce apoptosis.

Our genome-wide CRISPR/Cas9 loss of function screen surprisingly showed that both cyclin B (*CCNB1*) and Cdk2, canonically the kinase partner of cyclin A (*CCNA2*)^79,86^, were 2 of the top enriched genes that when inactivated induced resistance to cyclin A/B RxL inhibitors. Although the fact that cyclin B sgRNAs were enriched provided evidence that it is a target of the cyclin A/B RxL inhibitor, it was also surprising and suggested that cyclin A/B RxL inhibitors act to promote a gain of cyclin B function to kill cells. Unexpectedly we found that cyclin A/B RxL inhibitors promote a neomorphic interaction between cyclin B and Cdk2 in cells which promotes SAC activation and mitotic cell death (Fig. S12) which explains why both cyclin B and Cdk2 were top enriched hits in our genome-wide CRISPR/Cas9 loss of function screen. Together these data provide strong evidence that a component of the mode of action of cyclin A/B RxL inhibitors is to redirect Cdk2 to cyclin B to form cyclin B-Cdk2 complexes to phosphorylate RxL-independent substrates (i.e. stathmin), activate SAC, and consequently mitotic cell death (Fig. S12). Canonically cyclin A^72,87^ binds and activates Cdk2 in s-phase and cyclin B binds and activates Cdk1 in mitosis^72^. While cyclin B can bind Cdk2 *in vitro*^73^, it does not canonically do so during mitosis in cells. To our knowledge, this is the first demonstration of aberrant cyclin B-Cdk2 complex formation in cells that activates SAC and mitotic cell death, which in this case is achieved through a cyclin A/B RxL macrocycle. The precise mechanism by which cyclin A/B RxL macrocycles promotes the formation of cyclin B-Cdk2 neomorphic complexes in mitosis and whether the activity of Cdk2 in this complex is directly regulated by Myt1 will be an area for future investigation.

We focused this study on SCLC as SCLCs harbor near universal LOF mutations in *RB1* and *TP53*^2–6^ leading to high and dysregulated E2F activity and loss of cell cycle checkpoints, and because previous work showed that blocking cyclin:E2F1 RxL interactions selectively killed transformed cancer cell lines with high E2F activity^26^. We found that SCLCs with high E2F activity were hypersensitive to cyclin A/B RxL inhibitors and increasing E2F1 was sufficient to sensitize various other cell lines to cyclin A/B RxL macrocycles, demonstrating that high E2F activity primes cancer cells for killing by cyclin A/B RxL inhibitors. Together our data shows that cyclin A/B RxL inhibitors cause loss of cyclin A-mediated negative regulation of E2F1 activity thereby increasing E2F1 activity and sensitizing cells to cyclin A/B RxL inhibitors. Whether this increased E2F1 activity is restricted to S-phase or persists beyond S-phase into mitosis remains an open question. The mechanisms we identified in SCLC are likely broadly applicable to other cancer types as our data from HCT-116 cells, Jurkat cells, and MDA-MB-231 cells converged on a similar mechanism. Increased E2F activity can arise from *RB1* LOF mutations as in SCLC, or other genetic alterations that functionally inhibit pRB including *CDKN2A* LOF mutations or amplifications of cyclins^88^. Our cell line screening data in other cancer types (e.g. breast, ovarian) suggests that genomic alterations beyond *RB1* and/or *TP53* could correlate with cyclin A/B RxL inhibitor sensitivity. We hypothesize that high baseline E2F signatures could confer cyclin A/B RxL inhibitor sensitivity in other cancer types and potentially could be used as a biomarker to identify cancers more likely to respond to cyclin A/B RxL inhibitors.

One might expect complete inhibition of the RxL interactions of both cyclin A and cyclin B to be toxic as together cyclin A and B are essential for the proliferation of dividing cells^89,90^. However, we observed cytotoxicity at 1000-fold lower concentrations of cyclin A/B RxL inhibitors in sensitive cancer cells relative to cell lines that were inherently resistant including non-transformed fibroblasts. Moreover, these inhibitors were well tolerated in mice. This differential potency suggests that cyclin A/B RxL inhibitors do not simply block all cyclin A/B functions as combined cyclin A/B genetic inactivation is essential for survival of all cells.

Several previous studies have identified synthetic lethal targets with *RB1* loss in SCLC and other cancers. These synthetic lethal targets include Aurora A kinase^7,8^, Aurora B kinase^9^, PLK1^9,10^, and Mps1^11^ and all of these synthetic lethal targets function to regulate mitotic fidelity. Alisertib, an Aurora A kinase inhibitor showed activity in an early phase clinical trial when combined with paclitaxel in SCLCs with *RB1* pathway alterations^91^. Our data are consistent with these studies as we find that SCLCs with high E2F activity are hypersensitive to cyclin A/B RxL inhibitors, which promote SAC activation and apoptosis. A more recent study found that *CCNE1*-amplified tumors were hypersensitive to Myt1 inhibition which promotes mitotic cell death particularly when replication stress is induced with chemotherapeutic agents such as gemcitabine^69^. This study is consistent with our findings as they both converge on a mechanism where cyclin B:Cdk hyperactivation promotes mitotic cell death in tumors with genomic alterations that cause S-phase dysregulation. Cumulatively these studies suggest that cancers with *RB1* loss, often in the setting of *TP53* loss, are hyper-dependent on mitotic checkpoints for their survival.

In this work we describe cyclin A/B RxL macrocyclic inhibitors as a potential new class of therapeutics with a distinct mechanism of action from orthosteric Cdk2 inhibitors. We developed orally-bioavailable cyclin A/B RxL macrocyclic inhibitors that have strong anti-tumor activity in chemo-resistant SCLC PDXs models without overt toxicity. Our findings strongly suggest that cyclin A/B RxL inhibitors should be evaluated in patients with SCLC and other cancers driven by high E2F activity. Inspired by these and other supportive pre-clinical results, Circle Pharma is currently advancing CID-078, a further optimized, orally-bioavailable, first-in-class cyclin A/B RxL inhibitor into a Phase 1 clinical study to determine if this promising new mechanism of action can be leveraged to provide benefit to patients with SCLC and other cancers.

## Acknowledgements

We thank Dr. William G. Kaelin Jr. for insightful discussions and critical feedback and Neha Barrows for technical assistance. We are also grateful to Alan Ashworth (UCSF), Bruce Stillman (CSHL) and Jonathon Pines (ICR) for their advice. This work was supported by an SRA from Circle Pharma (M.G.O. and D.N.). M.G.O. is a William Raveis Charitable Fund Damon Runyon Clinical Investigator Award supported by the Damon Runyon Cancer Research Foundation (CI-101-19; M.G.O) and is supported by an NCI R37 grant (no. R37CA269990), and the Kaplan Family Fund. D.N. is a UT Presidential Scholar and holds the Joseph F. Sambrook, Ph.D. Distinguished Chair in Biomedical Science. D.N is supported by the Welch Foundation I-1879 and V-I-0002-20230731, and the Program in Molecular Medicine supported by an anonymous donor.

## Author Contributions

S.S.: Conceptualization, methodology, validation, formal analysis, investigation, data curation, writing. C.E.G.: Conceptualization, methodology, validation, formal analysis, investigation, data curation, writing. M.F, V.K, S.X, Y.T.D, A.S., Y.N.L., V.S., X.L., I.V., R.F.: Methodology, validation, formal analysis, investigation, data curation. M.A.: Investigation. R.O., F.H.I., D.H., M.W.M., L.H., M.P.B., B.M.L., J.F.L., D.H., Y.G., M.C., M.K.D., L.F.L., B.L., M.N., N.N.G.,S.S.F.L, B.F.W., C.B., J.A.S., K.Y., N.J.D., A.T.B.: Methodology, validation, formal analysis, investigation, data curation. D.S., A.T.B., S.M., J.G.D., B.K., J.B.A., E.W.W. C.K.: Resources, visualization, supervision, project administration. S.S: Resources, visualization, supervision, project administration. A.S: Methodology, resources, visualization, supervision, project administration. R.S., D.J.E., J.B.A., E.W.W., P.D.G., D.N., and M.G.O: Conceptualization, methodology, investigation, resources, data curation, writing – original draft, visualization, supervision, project administration, funding acquisition.

## Methods

### Cell Lines and Cell Culture

NCI-H446 (obtained 11/2016), NCI-H69 cells (10/2018), NCI-H1048 (9/2019), NCI-H526 (9/2019), WI-38 and 293T cells were originally obtained from American Type Culture Collection (ATCC). A549, HCC4006, and NCI-H1299 cells were a kind gift from Dr. Pasi Janne’s laboratory at DFCI. NCI-H1048, Jurkat and MDA-MB-231 cells with DOX-ON inducible E2F1 expression were a kind gift from Dr. William G. Kaelin’s lab.

NCI-H69, NCI-H446, HCC4006, NCI-H1299, NCI-H526, A549 and Jurkat cells were maintained in RPMI-1640 media supplemented with 10 % fetal bovine serum (FBS), 100 U/mL penicillin (P), and 100 μg/mL streptomycin (S). NCI-H1048 cells were cultured in RPMI-1640 media supplemented with 10% fetal bovine serum (FBS), P/S and Insulin-Transferrin-Selenium (Gemini). WI-38 cells were maintained in DMEM supplemented with 10% FBS. 293T cells were maintained in DMEM media with 10% FBS and P/S. MDA-MB-231 cells was maintained in DMEM/F12 with 10% FBS and P/S. Early passage cells of all the cell lines listed above were tested for mycoplasma (Lonza #LT07-218) and then were frozen using Bambanker’s freezing medium (Bulldog Bio) and maintained in culture no more than 4 months where early passage vials were thawed. All cells were iHCT116 cells previously described^59^ were cultured in 10% fetal bovine serum in DMEM media (Sigma) supplemented with 2 mM L-glutamine (Sigma).

### Pharmacological Inhibitors

Where indicated, the following chemicals (stored at −20°C or −80°C) were also added to the media: CIRc-004 (cyclin A/B RxL inhibitor; stock 10 mM in DMSO; obtained from Circle Pharma), CIRc-005 (inactive enantiomer of CIRc-004; stock 10 mM in DMSO; obtained from Circle Pharma), CIRc-001 (cyclin A/B RxL inhibitor; stock 10 mM in DMSO; obtained from Circle Pharma), Cdk2 inhibitor (PF-07104091; stock 10 mM in DMSO; chemically synthesized and validated to be the same PF-07104091 in the published patent), CIRc-018 (cyclin A selective RxL inhibitor; stock 10 mM in DMSO; obtained from Circle Pharma), CIRc-019 (cyclin B selective RxL inhibitor; stock 10 mM in DMSO; obtained from Circle Pharma), Mps1 inhibitor (BAY-1217389; Selleck chemicals #S8215; stock 10 mM), Doxycycline hydrochloride (Sigma #D3447, stock 1mg/mL), RP-6306 (MedChemExpress #HY-145817A, stock 20mM), roscovitine (Selleck Chemical #S115350MG stock 10mM), staurosporine (MedChemExpress #HY-15141 stock 20mM), and RO-3306 (ENZO, #ALX-270-463-M001, stock 10mM).

Binding potency of macrocycles for cyclin/Cdk complexes was determined by a Fluorescence Polarization (FP) competitive assay based on previously established protocols^27,29^ with the following modifications: 1) A higher affinity fluorescent labeled probe was identified referred as CIR7-2706 (Supplementary methods); 2) cyclin/CDK protein complexes were sourced as following: cyclin A2/Cdk2 (CRELUX Protein Services; www.crelux.com), cyclin B1/CDK1 (Eurofins, discovery. Cat. No. 14-450) and cyclin E1/Cdk2 (Eurofins, discovery. Cat. No. 14-475). FP binding assays were performed in 25 mM HEPES pH 7.5, 100 mM NaCl, 1mM DTT, 0.01% NP-40 and 1 mg/ml BSA for all 3 protein complexes in black 96-well plates (Costar #3356). After experimental plates are set, they were equilibrated by gentle mixing by placing them on an orbital shaker at 100 rpm for 2 hours at room temperature and then read on a SpectraMax i3X Multi-Mode Microplate Detection platform. The 50% inhibition concentration (IC_50_) of binding of a fluorescence labeled probe CIR7-2706 at 2 nM to the cyclin/Cdk complexed was determined from a 8-point three-fold serial dilution curve of CID-078. The protein concentration used were 8 nM for cyclin A2/Cdk2, and 10 nM for cyclin B1/Cdk1 and cyclin E1/Cdk2. Under these conditions, the assay dynamic range was about 120 mP units between 100 % binding of CIR7-2706 to the proteins and its complete displacement by an unlabeled competitor compound. All experiments showed a Z’ factor > 0.80. The reported IC_50_ are the average of all experiments described in Figure 1B.

### sgRNA Cloning to Make Lentiviruses

sgRNA sequences were designed using the Broad Institutes sgRNA designer tool (http://portals.broadinstitute.org/gpp/public/analysis-tools/sgrna-design) and chosen from the Brunello CP0043 sgRNA library and synthesized by IDT technologies. The sense and antisense oligonucleotides were mixed at equimolar ratios (0.25 nanomoles of each sense and antisense oligonucleotide) and annealed by heating to 100°C in annealing buffer (1X annealing buffer 100 mM NaCl, 10 mM Tris-HCl, pH 7.4) followed by slow cooling to 30°C for 3 hours. The annealed oligonucleotides were then diluted at 1:400 in 0.5X annealing buffer. For CRISPR/Cas9 knockout experiments in cells, the annealed oligos were ligated into pLentiCRISPR Puro V2 (Addgene #52961). Ligations were done with T4 DNA ligase for 2 hours at 25°C. The ligation mixture was transformed into HB101 competent cells. Ampicillin-resistant colonies were screened by restriction digestion of miniprep DNAs and subsequently validated by DNA sequencing.

The following sgRNA oligos were used for LentiCRISPRV2 Puro vector for CRISPR knockout experiments:

Non-targeting #1 sense (5’- CCGTCTCCGCATCGTCTTTT −3’)

Non-targeting #1 anti-sense (5’- AAAAGACGATGCGGAGACGG −3’)

Non-targeting #2 sense (5’- TATTTTGACTTGACGCAGGC −3’)

Non-targeting #2 anti-sense (5’- GCCTGCGTCAAGTCAAAATA −3’)

Human *CCNB1* #1 sense (5’- CATCAGAGAAAGCCTGACAC −3’)

Human *CCNB1* #1 anti-sense (5’- GTGTCAGGCTTTCTCTGATG −3’)

Human *CCNB1* #2 sense (5’- GAGGCCAAGAACAGCTCTTG −3’)

Human *CCNB1* #2 anti-sense (5’- CAAGAGCTGTTCTTGGCCTC −3’)

Human *CDK2* #1 sense (5’- CAAATATTATTCCACAGCTG −3’)

Human *CDK2* #1 anti-sense (5’- CAGCTGTGGAATAATATTTG −3’)

Human *CDK2* #2 sense (5’- CATGGGTGTAAGTACGAACA −3’)

Human *CDK2* #2 anti-sense (5’- TGTTCGTACTTACACCCATG −3’)

Human *KNTC1* #1 sense (5’- AAACATTCGGAACACTATGG −3’)

Human *KNTC1* #1 anti-sense (5’- CCATAGTGTTCCGAATGTTT −3’)

Human *KNTC1* #2 sense (5’- AAGCTAACGATGAAAATCGG −3’)

Human *KNTC1* #2 anti-sense (5’- CCGATTTTCATCGTTAGCTT −3’)

Human *LIN54* #1 sense (5’- GTTCTTTCACAGTCTACTCC −3’)

Human *LIN54* #1 anti-sense (5’- GGAGTAGACTGTGAAAGAAC −3’)

Human *LIN54* #2 sense (5’- AAGTAGTTACCATTGGAGGG −3’)

Human *LIN54* #2 anti-sense (5’- CCCTCCAATGGTAACTACTT −3’)

Human *MAD1L1* #1 sense (5’- GAAGAAGCGCGAGACCCACG −3’)

Human *MAD1L1* #1 anti-sense (5’- CGTGGGTCTCGCGCTTCTTC −3’)

Human *MAD1L1* #2 sense (5’- GAGAGACTGGACCAGACCAT −3’)

Human *MAD1L1* #2 anti-sense (5’- ATGGTCTGGTCCAGTCTCTC −3’)

Human *ZWINT* #1 sense (5’- CACCTACAGGGAGCACGTAG −3’)

Human *ZWINT* #1 anti-sense (5’- CTACGTGCTCCCTGTAGGTG −3’)

Human *ZWINT* #2 sense (5’- CAGTGGCAGCTACAACAGGT −3’)

Human *ZWINT* #2 anti-sense (5’- ACCTGTTGTAGCTGCCACTG −3’)

### cDNA Expression Plasmids

CDC20 (NM_001255.3) or CDC20 with an N-terminal 3xFlag tag were cloned in the pLVX IRES Hygro vector (Clontech). The CDC20 R445Q mutation were generated using site directed mutagenesis (Pfu polymerase). pTwist Lenti SFFV Puro WPRE lentiviral vectors encoding: 1) CCNB1 WT-HA or 2) CCNB1 Triple Mutant-HA (E169K/Y170H/Y177C) or a 3) negative control which encodes a functionally inactivated EED Mutant-HA with the following mutations to inactivate EED (F97A/Y148A/Y365A) were codon optimized and synthesized by Twist Biosciences. All plasmids were sequence verified by sanger sequencing.

### Lentivirus Production

Lentiviruses were made by Lipofectamine 2000-based co-transfection of 293FT cells with the respective lentiviral expression vectors and the packaging plasmids psPAX2 (Addgene #12260) and pMD2.G (Addgene #12259) in a ratio of 4:3:1. Virus-containing supernatant was collected at 48 and 72h after transfection, pooled together (7 mL total per 6-cm tissue culture dish), passed through a 0.45-µm filter, aliquoted, and frozen at −80°C until use.

### Lentiviral Infection

The cells were counted using a Vi-Cell XR Cell Counter (Beckman Coulter) and 2 X10^6^ cells were resuspended in 1mL lentivirus with 8 μg/mL polybrene in individual wells of a 12 well plate. The plates were then centrifuged at 931 x g [Allegra® X-15R Centrifuge (Beckman Coulter), rotor SX4750A, 2000 rpm] for 2h at 30°C. 16 hours later the virus was removed and cells were grown for 72 hours before being placed under drug selection. Cells were selected in puromycin (0.5 μg/mL), or G418 (500 μg/mL).

### Cyclin A/B RxL Inhibitor, Cdk2 inhibitor and Mps1 Inhibitor Dose Response Assays

For dose response assays, with cyclin A/B RxL inhibitor (CIRc-004), inactive enantiomer of CIRc-004 (CIRc-005) and Cdk2 inhibitor (PF-07104091), the cells were counted on day 0 using a Vi-Cell XR Cell Counter and plated in a tissue culture-treated 6-well plate at 50,000 cells/mL in 2 mL of complete media. Cells were treated with the compounds (CIRc-004, CIRc-005, PF-07104091) at indicated concentrations for 6 days. Normalized cell count was calculated as (Day 6 count for treated samples) / (Day 6 count for DMSO treated sample). The half-maximal inhibitory concentration (EC50) was calculated using GraphPad Prism 10.0.0 software.

For doxycycline inducible (DOX-ON) E2F1 overexpressing cell lines, cells were counted on day 0 and plated in a tissue culture-treated 6-well plate at 250,000 cells/mL for 3-day assays or 50,000 cells/mL in 2 mL of complete media for 6-day assays. For 3-day cyclin A/B RxL inhibitor assays, E2F1 expression was induced with DOX (1 ug/mL) for 24 hours followed by drug treatment for 3 days. For 6-day cyclin A/B RxL inhibitor assays, E2F1 expression was induced with DOX for 24 hours followed by drug treatment for 6 days where fresh DOX was replaced every 3 days.

### MTT Proliferation Assays

NCI-H1048, NCI-H446 and WI-38 cells were counted using a Countess II FL (Invitrogen) and plated at 5,000 cells/well in 96-well plates. NCI-H69 cells were grown to confluency in a T150 flask. Cells were collected by centrifugation at 283 x g (Thermo Scientific X4R Pro-MD centrifuge, Sorvall TX1000 rotor, 1100 rpm)and resuspended in 30 mL media of which 10ml was used to seed six 96-well plates. NCI-H526 cells were grown to confluency in a T75 flask, collected by centrifugation at 283 x g and resuspended in 10 mL media of which 2.5ml was used to seed six 96-well plates. The following day, cells were dosed with cyclin A/B/E RxL inhibitor (CIRc-001), cyclin A/B RxL inhibitors (CIRc-004; CIRc-028), cyclin A selective RxL inhibitor (CIRc-018), cyclin B selective RxL inhibitor (CIRc-019), or inactive enantiomer of CIRc-004 (CIRc-005). Roscovitine and staurosporine were used as plate controls to define the top and bottom of the growth inhibition curves, respectively. Inhibitors were dosed in duplicate in either an 8-(WI-38) or 10-point (SCLC cell lines) 1:3 serial dilution with 10 μM maximum concentration. Roscovitine was dosed in singlet in an 8- or 10-point 1:2 serial dilution with 100 μM maximum concentration. Staurosporine was dosed in singlet in an 8- or 10-point 1:2 serial dilution with 1 μM maximum concentration. After dosing, plates were maintained in tissue culture incubators (37°C; 5% CO_2_) for 3 days (WI-38) or 5 days (SCLC cell lines) to allow for at least 2 cell doublings before processing in an MTT proliferation assay (R&D systems, #4890-050-K) performed according to manufacturer instructions. The average absorbance value obtained with the highest two concentrations dosed for roscovitine and staurosporine was used for background subtraction. The top of the assay (100% growth) was determined by normalization with the top of the roscovitine curve. Growth inhibition 50 (GI50) was determined by nonlinear regression analysis using log(inhibitor) vs response -variable slope (four parameters) using GraphPad Prism (10.1.0) software.

For MTT proliferation assays with the Myt1 inhibitor RP-6306, NCI-H1048, NCI-H446 and NCI-H69 cells were seeded in 96-well plates as described above. The following day, cells were dosed in duplicate in a 10-point 1:3 serial dilution with CIRc-018, CIRc-019, or CIRc-004 alone or in combination with RP-6306 at 100 nM for NCI-H1048 cells or 500 or 1000 nM for NCI-H446 and 1000 nM for NCI-H69 cells. After 5 days, cells were processed in an MTT proliferation assay and GI50s determined using Graphpad Prism software (10.1.0) as described above.

### Dose response studies in iHCT-116 Cells

Viability of cells were quantified using Cell Titer Glo according to manufacturer instructions (Promega) and the IC50 values were determined using Graphpad prism (log inhibitor versus response four parameters).

### FACS-based cleaved PARP Apoptosis Assays

Cells were plated at 250,000 cells/mL density and grown in the presence or absence of cyclin A/B RxL inhibitor (CIRc-004) or Mps1 inhibitor (BAY-1217389) at indicated concentrations for 72 hours. Cells were then collected and fixed in 4% paraformaldehyde followed by a PBS wash, and then permeabilized in ice-cold 90% methanol (added dropwise) for 30 mins on ice. The cells were then centrifuged at 206 x g (Eppendorf centrifuge 5804 R, A-4-44 rotor, 2000 rpm) washed once in PBS, centrifuged again at 206 x g, and then incubated with Pacific blue conjugated cleaved PARP antibody (Cell Signaling, Asp214, D64E10, #60068) at a dilution of 1:100 for 1 hour at room temperature protected from light. Cells were then washed again in PBS containing 0.5% BSA, resuspended in PBS containing 0.5% BSA and analyzed on LSR Fortessa flow cytometer (Becton Dickinson, Franklin Lakes, NJ). Analysis for cleaved PARP positive cells was carried out using FlowJo 10.8.1.

### FACS-based phospho-histone H3 (Ser10) Analysis

Cells were plated at 500,000 cells/mL and grown in the presence or absence of cyclin A/B RxL inhibitor (CIRc-004) at indicated concentrations for 24 hours. The cells were then washed once in room temperature PBS and then fixed in 4% paraformaldehyde for 15 minutes at room temperature. The cells were then centrifuged at 206 x g (Eppendorf centrifuge 5804 R, A-4-44 rotor, 2000 rpm) for 5 mins at room temperature, washed once in PBS, centrifuged again at 206 x g, and then permeabilized with ice cold methanol at 4° C for 30 minutes. The cells were then washed again in PBS and then incubated with Alexa-647 conjugated phospho-histone H3 (Ser10) antibody (Cell Signaling, #3458) at a dilution of 1:100 for 1 hour at room temperature, then washed once in PBS containing 0.5% BSA, centrifuged at 400 x g, and then counter-stained with 4’, 6’-diamidino-2-phenylindole (DAPI) (Cell Signaling #4083) for 15 minutes at room temperature. FACS analysis was then performed to determine the % positive phospho-histone H3 (Ser 10) cells using FlowJo 10.8.1.

### FACS-based Propidium Iodide (PI) Cell Cycle Analysis

Cells were plated at 500,000 cells/mL and grown in the presence or absence of the cyclin A/B RxL inhibitor (CIRc-004) at indicated concentrations for 24 hours. Following incubation, cells were washed once in ice-cold PBS and then fixed in ice-cold 80% ethanol (added dropwise) for at least 2 hours at −20°C. The cells were then centrifuged at 206 x g (Eppendorf centrifuge 5804 R, A-4-44 rotor, 2000 rpm) for 5 mins, washed once in PBS, centrifuged again at 206 x g, and then washed again in PBS containing 0.5% BSA. Finally, cells were stained with Propidium Iodide (PI) (BD # 550825) for 15 minutes at room temperature. FACS analysis for PI was then performed on a LSR Fortessa flow cytometer (Becton Dickinson, Franklin Lakes, NJ) and analysis was carried out using FlowJo 10.8.1.

### FACS-based EdU Incorporation and DNA Content Analysis

NCI-H1048 cells were plated at 500,000 cells/ml in a 12-well plate and allowed to adhere overnight. Cells were dosed with DMSO or cyclin A/B RxL inhibitor for 24 hours. Cells were pulsed for 1h with 10μM EdU (Click-It Plus EdU Flow Cytometry Assay kit, Invitrogen #C10632) prior to collection. Following incubation, cells were washed once with ice-cold PBS and fixed by incubation in ice-cold 80% ethanol for at least 2 hours at −20°C. EdU was fluorescently tagged with Alexa Fluor^TM^ 488 by click reaction (Click-it Plus EdU flow cytometry assay kit, Invitrogen, #C10632). DNA content was monitored by FxCycle Violet stain (Invitrogen, #F10347). Flow cytometry analysis for EdU and DNA content was performed by full spectral flow cytometry on a Northern Lights cytometer (Cytek Biosciences) and analysis was carried out using FlowJo 10.9.0.

### Immunoblotting

Cell pellets were lysed in a modified EBC lysis buffer (50mM Tris-Cl pH 8.0, 250 mM NaCl, 0.5% NP-40, 5 mM EDTA) supplemented with a protease inhibitor cocktail (Complete, Roche Applied Science, #11836153001) and phosphatase inhibitors (PhosSTOP Sigma #04906837001). Soluble cell extracts were quantified using the Bradford Protein Assay. 20 µg of protein per sample was boiled after adding 3X sample buffer (6.7% SDS, 33% Glycerol, 300 mM DTT, and Bromophenol Blue) to a final concentration of 1X, resolved by SDS-PAGE using either 10% or 6% SDS-PAGE, semi-dry transferred onto nitrocellulose membranes, blocked in 5% milk in Tris-Buffered Saline with 0.1%Tween 20 (TBS-T) for 1 hour, and probed with the indicated primary antibodies overnight at 4°C. Membranes were then washed three times in TBS-T, probed with the indicated horseradish peroxidase conjugated (HRP) secondary antibodies for 1 hour at room temperature, and washed three times in TBS-T. Bound antibodies were detected with enhanced chemiluminescence (ECL) western blotting detection reagents (Immobilon, Thermo Fisher Scientific, #WBKLS0500) or Supersignal West Pico (Thermo Fisher Scientific, #PI34078). The primary antibodies and dilutions used were: Mouse Anti-β-actin AC-15 (Sigma #A3854, 1:25,000), mouse anti-Vinculin hVIN-1 (Sigma# V9131, 1:10000), rabbit anti-phospho CASC5/KNL1 Thr943/Thr1155 D8D4N (Cell Signaling #40758, 1:1000), rabbit anti-CASC5/KNL1 E4A5L (Cell Signaling #26687, 1:1000), rabbit anti-cyclin B1 D5C10 (Cell Signaling #12231, 1:1000), rabbit anti-E2F1 (Cell Signaling #3742, 1:1000), CDC20 (Cell Signaling #4823S, 1:1000), and FLAG (Sigma-Aldrich #A8592). The secondary antibodies and dilutions were: Goat Anti-Mouse (Jackson ImmunoResearch #115-035-003, 1:50000) and Goat anti-Rabbit (Jackson ImmunoResearch #111-035-003, 1:5000).

To detect phospho-Stathmin S38, total Stathmin, phospho-Cyclin B1(S126), phospho-nucleolin (T84), phospho-RPA2(S33) and phospho-FOXM1(T600), after incubation with the indicated cyclin RxL inhibitors or DMSO control, NCI-H1048 cells were washed once in ice-cold PBS and lysed in 1% Triton X-100 buffer (40 mM HEPES pH7.5, 1mM EDTA pH 8.0, 10mM sodium pyrophosphate, 10 mM beta-glycerophosphate, 50 mM sodium fluoride, 120 mM sodium chloride and 1% Triton X-100) supplemented with a protease inhibitor cocktail (cOmplete, Sigma #5892970001) and phosphatase inhibitors (PhosSTOP, Sigma #4906837001). Cell lysates were centrifuged at 9391 x g (Eppendorf centrifuge 5424R, FA-45-24-11 rotor, 10,000 rpm) for 10 min. Supernatant (cell extract) was removed and protein content estimated by performing a Bradford Assay. Cell extract (30 micrograms of protein) was boiled after adding 4X Laemmli buffer and separated by SDS-PAGE and transferred to PVDF membranes. Membranes were blocked in TBS plus 0.2% Tween-20 (TBS-T) containing 5% skim milk and incubated with relevant primary antibodies overnight at 4°C in blocking buffer containing 5% bovine serum albumin (BSA). After incubation with primary antibodies, membranes were washed in TBS-T and then incubated with HRP-labeled secondary antibodies for 1-2h at room temperature. Membranes were washed again in TBS-T, incubated with ECL reagent (LI-COR WesternSure #92695000) and scanned on an Odyssey Fc Analyzer (LI-COR Biosciences). The primary antibodies and dilutions used were: rabbit-anti-phospho-stathmin (S38) (Cell Signaling #3426S, 1:1000), rabbit-anti-total stathmin (Abcam #ab52630), rabbit-anti-phospho-Cyclin B (S126) (Abcam #ab3488, 1:1000), rabbit-anti-phospho-nucleolin (T84) (Abcam #ab155977, 1:1000), rabbit-anti-phospho-FOXM1(T600) (Cell Signaling #14655S, 1:500), rabbit-anti-phospho-RPA2(S33) (Fortis Life Sciences #A300-246A, 1:2000), rabbit-anti-tubulin-HRP (Cell Signaling #9099S 1:5000). The secondary antibodies used and dilutions were: goat-anti-rabbit-IgG-HRP (Cell Signaling #7074S, 1:5000).

### Immunoprecipitations

For immunoprecipitation (IP) of endogenous cyclin A and cyclin B in Fig. 3I and 5C, NCI-H1048 cells were seeded at 2.5 × 10^7^ cells/15 cm plate and allowed to adhere overnight. The following day, cells were incubated with the indicated cyclin RxL inhibitors at 300 nM or DMSO for 2 hours before rinsing once in ice-cold PBS and lysing in 1% NP-40 buffer (40 mM HEPES pH7.5, 1mM EDTA pH 8.0, 10 mM sodium pyrophosphate, 10 mM beta-glycerophosphate, 50 mM sodium fluoride, 120 mM sodium chloride and 1% NP-40) supplemented with a protease inhibitor cocktail (cOmplete, Sigma #5892970001) and phosphatase inhibitors (PhosSTOP, Sigma #4906837001). Cell lysates were centrifuged at 9391 x g for 10 min. Supernatant (cell extract) was removed and protein content estimated by performing a Bradford Assay. 2 mg cell extract was rotated overnight at 4°C with 2 μg anti-cyclin A [Santa Cruz Biotechnology (SCBT) sc-271682] or 2 μg anti-cyclin B antibody (SCBT sc-245). The next day, 20 μl packed protein A and protein G coated magnetic beads (Dynabeads^TM^ Protein A and Dynabeads^TM^ Protein G, Invitrogen, #10002D and #10003D, respectively) were added and lysates were rotated for an additional 1 hour at 4°C. The protein A/G beads were collected using a magnet, washed three times with 1% NP-40 cell lysis buffer, boiled and denatured in 1X Laemmli sample buffer, separated by SDS-PAGE, and immunoblotting was performed as described above. The primary antibodies and dilutions used were: anti-Myt1 (Fortis Life Sciences A302-424A 1:1000), mouse-anti-cyclin B (Cell Signaling #4135 1:1000), mouse-anti-E2F1 (SCBT #sc-251, 1:1000), mouse-anti-cyclin A (Cell Signaling #4656, 1:1000), mouse anti-CDK2 (Origene TA502935), rabbit anti-Cdc2 (Cell Signaling 28439). The secondary antibodies used and dilutions were: mouse-anti-rabbit (confirmation specific)-IgG HRP (Cell Signaling #5127, 1:2000), goat-anti-rabbit IgG-HRP (Cell Signaling #7074S, 1:5000), horse-anti-mouse IgG-HRP (Cell Signaling #7076S, 1:5000), goat-anti-mouse-IgE-HRP (Southern Biotech #1110-05, 1:5000).

For IPs of cyclin B-HA and Cdk2 in Fig. 4I and S9G, HEK-293T cells were stably infected with pTwist Lenti SFFV Puro WPRE lentiviral vectors (Synthesized by Twist Biosciences) encoding: 1) CCNB1 WT-HA or 2) CCNB1 Triple Mutant-HA (E169K/Y170H/Y177C) or a 3) negative control which encodes a functionally inactivated EED Mutant-HA with the following mutations to inactivate EED (F97A/Y148A/Y365A). 20 × 10^6^ cells were seeded at 1 × 10^6^ cells/ml density and next day treated with the cyclin A/B RxL inhibitor (CIRc-004) at 300 nM or DMSO for 2 hours before harvesting cells. Cells were lysed in 700 µl lysis buffer (Cell Signaling, #9803) supplemented with protease inhibitor cocktail (Complete, Roche Applied Science, #11836153001) and phosphatase inhibitors (PhosSTOP Sigma #04906837001). Samples were sonicated (Branson Digital Sonifier) 3 times for 5 seconds each to ensure homogenization and centrifuged at 14000 x g for 10 minutes at 4° C (Eppendorf centrifuge 5424 R, FA-45-24-11 rotor). Supernatant was precleared by rotating 20 µl Protein A agarose beads (Cell Signaling #9863) for 1 hour at 4° C. Samples were then briefly vortexed to remove the beads and protein concentration was determined. 60 µl of lysate was removed as input (∼10% total for ∼2.5% input per immunoblot) and boiled after adding 3X sample buffer (6.7% SDS, 33% Glycerol, 300 mM DTT, and Bromophenol Blue) to a final concentration of 1X. Then 300 µl of lysate was immunoprecipitated by rotating overnight at 4° C with rabbit anti-HA epitope (Cell Signaling #3724, 1:100) for exogenous cyclin B, rabbit anti-Cdk2 E8J9T (Cell Signaling #18048, 1:100) or equivalent amount of Rabbit (DA1E) mAb IgG XP Isotype Control (Cell Signaling #3900). The following day, 20 µl of Protein A agarose beads were added to each sample tube and incubated for 1 hour at 4° C with rotation. The beads were collected by centrifugation and washed five times with cold lysis buffer followed by boiling in 3X sample buffer. Samples were run on SDS-PAGE and immunoblotting was performed as described above. Primary antibodies used for immunoblotting were; rabbit anti-Cyclin B1 D5C10 (Cell Signaling #12231, 1:1000), rabbit anti-Cdk2 E8J9T (Cell Signaling #18048, 1:1000) and mouse anti-HA. 11 epitope tag (BioLegend #901501, 1:1000).

### Histone Extractions and Histone Immunoblot Analysis

For histone extractions in Fig. 5, cells were counted on day 0 using the Vi-Cell XR Cell Counter and plated in 6 cm plates at 250,000 cells/mL in 4 mL of media per plate for all cell lines along with indicated concentration of cyclin A/B RxL inhibitor. Three days later, cells were pelleted by centrifugation at 400 x g for 3 minutes at 4°C in a 15 mL conical tube. The cell pellets were then washed once in 1 mL of ice-cold PBS, transferred to a 1.5 mL Eppendorf tube, and pelleted by centrifugation at 400 x g for 3 minutes at 4°C. The PBS was removed by gentle aspiration and the soluble proteins from the cell pellets were extracted by incubation in Nucleus Lysis Buffer (250 mM Sucrose, 60 mM KCL, 15 nM NaCl, 15 mM Tris pH 7.5, 5 mM MgCl_2_, 1 mM CaCl_2_, 1 mM DTT, 0.3% NP-40 supplemented with a cocktail of protease inhibitor cocktail and phosphatase inhibitors) for 10 min at 4°C. The extracts were then centrifuged at 10,000 x g for 1 minutes at 4°C and the supernatant was removed by aspiration. The cell pellets were again resuspended in Nucleus Lysis Buffer and incubated for 10 minutes at 4°C and then centrifuged at 10,000 x g for 1 minute at 4°C. The supernatant was removed by aspiration and the histones in the insoluble pellet were extracted by overnight incubation in 60 μl of 0.2 N HCl followed by centrifugation at 16,800 x g for 15 minutes at 4°C. The supernatant was transferred to a fresh 1.5 mL Eppendorf tube and quantified using the Bradford Protein Assay. The pH of the supernatant was neutralized by adding 1/5 volume of 1 N NaOH, as confirmed by the color of the samples after the addition of sample buffer containing Bromophenol Blue. 1 μg of protein in sample buffer was boiled and resolved by SDS-polyacrylamide gel electrophoresis on a 15% SDS-PAGE gel. Immunoblot analysis was performed as described above. The primary antibodies were: rabbit anti-Histone 3 DH12 (Cell Signaling #4499, 1:5000) and rabbit anti-phospho Histone H2A.X Ser 139 (Cell signaling #2577, 1:1000).

### Cancer Cell Line Sensitivity Screening with Cyclin A/B RxL Inhibitors

A panel of 46 small cell lung cancer cell lines were tested for their sensitivity to Cyclin A/B RxL inhibition in a Cell Titer Glo (CTG) proliferation assay performed at Shanghai ChemPartner (Pudong, Shanghai, China) (see Fig. 1D). Cells were plated in 384-well format and dosed in duplicate with CIRc-004 in a 10-point, 1:3 serial dilution dose-response assay with 10 μM starting concentration. Staurosporine was used as an assay control per cell line and was dosed in duplicate from 2 μM maximum concentration in a 10-point 1:3 serial dilution. Cells were exposed to inhibitor for 4-8 days depending on the length of time required for at least two cell doublings to occur. A time ‘0 hour’ read and an endpoint DMSO control read were obtained per cell line for calculation of GI50s.

CIRc-001 was profiled in Horizon Discovery’s High Throughput OncoSignature screening platform containing 300 cell lines across 18 indications (see Fig. S2A). Cells were seeded in 384-well tissue culture plates at 250-1500 cells per well. Assay plates were dosed in triplicate with inhibitor in a 9-point, 1:3 serial dilution dose-response assay with 10 μM maximum concentration. Cells were incubated for 3 days and then analyzed using CellTiter-Glo 2.0 (Promega). At the time of treatment, a set of assay plates which do not receive treatment were collected and ATP levels measured by adding CellTiter-Glo 2.0 (Promega). All data points were collected via automated processes and were subject to quality control. Potency and efficacy metrics were derived from logistics curves fitted to growth inhibition using Horizon’s Chalice software.

### Gene Set Expression Analyses

The Gene Set Variation Analysis (GSVA) method^49^, utilizing the MSigDb Hallmark collection^50^ of RNA-seq data, was employed to calculate GSVA scores for E2F targets and G2/M checkpoint pathway using the GSVA Bioconductor package (version 1.50.5). To visualize pathway-level alterations, gene expression data was first aggregated to hallmark pathways as GSVA score based on gene set annotations obtained from MSigDB. For each pathway, pathway scores were transformed to z-scores across samples to standardize the data. The heatmap was constructed using the ComplexHeatmaps package (version 2.18.0) in R (version 4.3.2), where rows represented hallmark pathways, columns represented samples, and the color intensity reflected the mean z-score of each pathway for each sample. The heatmap was hierarchically clustered by pathway to reveal patterns of enrichment (Fig.1E).

### CRISPR/Cas9 Resistance Screen with the Cyclin A/B RxL inhibitor, the Cyclin A/B/E RxL Inhibitor, or the Cdk2 Inhibitor

On day 0, NCI-H1048 cells were counted using the Vi-Cell Counter. 150 × 10^6^ cells (which would yield representation of at least 500 cells/sgRNA if MOI=0.3) were pelleted and resuspended at 2 × 10^6^ cells/mL in complete media containing 50 µl/mL of human whole genome Brunello sgRNA library (CP0043) (Purchased from the Broad Institute) lentivirus which contains both the sgRNA and Cas9, and 8 µg/mL polybrene. The lentiviral titer was determined empirically in pilot experiments with a goal multiplicity of infection (MOI) of 0.3-0.5. CP0043 contains 77,441 sgRNAs targeting 4 sgRNAs per gene with 1000 non-targeting sgRNAs controls. The cells mixed with polybrene and lentivirus were then plated in 1 mL aliquots onto 12 well plates and centrifuged at 931 x g [Allegra® X-15R Centrifuge (Beckman Coulter), rotor SX4750A, 2000 rpm], for 2 hours at 30°C. In parallel, 2 × 10^6^ cells were also centrifuged under same conditions but without lentivirus as a control for puromycin selection (Mock). ∼16 hours later (day 1), the cells were collected, pooled, and centrifuged again at 448 x g to remove the lentivirus and polybrene, and the cell pellet was resuspended in complete media and plated into 23 15 cm tissue culture treated plates at 0.3 × 10^6^ cells/mL or in 6 well plates at the same density for the control (Mock) cells. The cells were then cultured for 72 hours at which time (day 4) the cells were counted and plated at 0.4 × 10^6^ cells/mL in 15 cm tissue culture treated plates with fresh media containing puromycin (0.25 µg/ml) to select for puromycin-transduced cells. A parallel experiment was performed on day 4 to determine the MOI of the screen. To do this, cells infected with the sgRNA library or mock-infected cells were plated at 0.4 × 10^6^ cells/mL in 6 well plates in the presence or absence of puromycin (0.25 µg/ml). After 72 hours (day 7), cells were counted using the Vi-Cell XR Cell Counter and the MOI was calculated using the following equation: (# of puromycin-resistant cells infected with the sgRNA library/# total cells surviving without puromycin after infection with the sgRNA library) – (# of puromycin-resistant mock-infected cells/ # total mock-infected cells). The calculated MOI was 0.33. On the same day (day 7), puromycin containing media was exchanged with fresh complete media not containing puromycin.

On day 10, cells were pooled, counted and split into two replicates. Each replicate was split into four arms: 1. CIRc-004 (cyclin A/B RxL inhibitor, 200 nM); 2. CIRc-001 (cyclin A/B/E RxL inhibitor, 200 nM), 3. Cdk2 inhibitor (PF-07104091, 500 nM); or CIRc-005 (inactive enantiomer of CIRc-004, 200 nM). CIRc-005 had no effect on cellular proliferation (see Fig. S2B) and therefore was used as a negative control comparison for all drug treated arms. A total of 4 × 10^7^ cells were plated at 0.2 × 10^6^ cells/mL in 15cm tissue culture plates in complete media containing the respective drugs listed above maintaining 500 cells/sgRNA. At the same time, the remaining cells were collected and divided in aliquots of 4 × 10^7^ (again to maintain representation of 500 cells/sgRNA), washed in PBS, and cell pellets were frozen for genomic DNA isolation for the initial timepoint prior drug selection. Mock-infected cells were maintained at the same density for all drug conditions and carried in parallel for the entire screen described below.

The media on the plates under drug selection was exchanged every 3 days with complete media containing fresh drug. Treatment arms were counted and passaged when cells were ∼80% confluent and split back to a representation of 4 × 10^7^ cells at a density of 0.2 × 10^6^ cells/mL onto 15 cm tissue culture plates. This occurred for the CIRc-005 inactive enantiomer every 3 days as CIRc-005 had no effect on cellular proliferation. This occurred only once or twice during the screen for all other drug treatment conditions as the drugs exhibited strong cytotoxicity. When cells were passaged, the cells were pooled, counted, and replated maintaining 500 cells/sgRNA. The remaining cells were divided in aliquots of 4 × 10^7^ cells, washed in PBS and cells pellets were frozen for genomic DNA isolation. The screen was terminated on Day 26 when there was an obvious growth advantage in the CIRc-004, CIRc-001, Cdk2 inhibitor arms infected with the sgRNA library relative to the mock infected cells treated with the same compounds.

Following completion of the screen, genomic DNA was isolated using the Qiagen Genomic DNA maxi prep kit (cat. # 51194) according to the manufacturer’s protocol. Raw Illumina reads were normalized between samples using: Log2[(sgRNA reads/total reads for sample) × 1e6) + 1]. The initial common time point data (day 10) was then subtracted from the end time point after drug treatment (day 26) to determine the relative enrichment of sgRNAs in the CIRc-004, CIRc-001, and Cdk2 inhibitor arms relative to the CIRc-005 inactive enantiomer control arm for each replicate. Apron analysis, hypergeometric analysis, and STARS analysis (GPP web portal, https://portals.broadinstitute.org/gpp/screener/) were performed using the normalized counts of the drug treated arms-early timepoint (Apron) or the log normalized counts of the drug treated arms-early timepoint (Hypergeometric, STARS) comparing the drug treated condition to either the CIRc-005 enantiomer control at the end timepoint or compared to the early timepoint. The averaged data from the 2 replicates were used for all analyses shown in the manuscript.

### Dose Response Assays for Screen Hit Validation with CRISPR/Cas9 sgRNA Knockout

NCI-H1048 and NCI-H446 cells were first infected with pLentiCRISPR V2-Puro lentiviruses encoding sgRNAs targeting the selected screen hits and selected with puromycin. Stably infected cells were counted on day 0 using a Vi-Cell XR Cell Counter and plated on tissue culture-treated 6 well plates at 50,000 cells/mL in 2 mLs of complete media. For dose response assays with cyclin A/B RxL inhibitor (CIRc-004), and Cdk2 (PF-07104091) inhibitor, the cells were treated with the compounds at indicated concentrations for 6 days and counted using the Vi-Cell. Normalized cell counts were calculated as Day 6 count for treated samples/Day 6 count for DMSO treated sample. Analysis was performed using GraphPad Prism 10.0.0 software.

### Forward Genetic Screen with Cyclin A/B RxL Inhibitor in iHCT-116 Cells

Five million iHCT116 cells^59^ that had previously been treated with vehicle or indole acetic acid were cultured in 10 μM of cyclin A/B RxL inhibitor (CIRc-004) for 17 days replenishing the media every three days. The surviving clones were analyzed for resistance to CIRc-004 in a three day proliferation assay where CIRc-004 was added in dose response with a final concentration of DMSO of 0.5%. Barcode sequences were PCR amplified and sequenced as described previously^59^. Genomic DNA was collected from select clones (Qiagen) and processed for whole exome sequencing. Mutations unique to individual clones were identified using the previously described analysis^59^.

### Rescue Experiments with Mps1 Inhibitor

For dose response assays with Mps1 inhibitor (BAY-1217389) to empirically identify concentrations of the Mps1 inhibitor that effectively blocked Mps1 activity measured by immunoblotting for phospho-KNL1 without gross effects on cellular proliferation, the cells were counted on day 0 using a Vi-Cell XR Cell Counter and plated in a tissue culture-treated 6-well plate at 250,000 cells/mL in 2 mL of complete media. Cells were treated with the drug (BAY-1217389) at indicated concentrations for 3 days. The normalized cell count was calculated as (Day 3 count for treated samples)/(Day 3 count for DMSO treated sample). Analysis was performed using GraphPad Prism 10.0.0 software.

For rescue experiments with the Mps1 inhibitor, cells were counted using the Vi-Cell XR Cell and plated at 250,000 cells/mL for NCI-H1048 cells or at 50,000 cells/ml for NCI-H69 and NCI-H446 cells and then treated with cyclin A/B RxL inhibitor (CIRc-004) at indicated concentrations in presence or absence of Mps1 inhibitor (BAY-1217389 at 3 nM) and harvested after 3 days for dose-response assays using the Vi-Cell. For FACS based cleaved PARP apoptosis assay, cells were counted and plated at 250,000 cells/mL for all the cell lines and then treated with cyclin A/B RxL inhibitor (CIRc-004) at indicated concentrations in presence or absence of Mps1 inhibitor (BAY-1217389 at 3 nM), and harvested after 3 days, stained and analyzed as described above.

For immunoblot analysis of phospho-KNL1 levels after treatment with BAY-1217389, ∼2 million cells were counted using the Vi-Cell XR Cell and plated at 500,000 cells/mL and then treated with cyclin A/B RxL inhibitor (CIRc-004) at indicated concentrations in presence or absence of BAY-1217389 at 3 nM, and harvested after 24 hours for immunoblot analysis.

### Immunofluorescence

Cells were seeded onto glass coverslips at 250,000 cells/mL and treated with CIRc-004 as indicated. Coverslips were fixed with ice cold 100% methanol for 15 min on ice. Cells were washed with PBS and incubated in blocking buffer (5% BSA in PBS) for 1 hour. Coverslips were then incubated with the following primary antibodies diluted in fresh blocking buffer: rabbit anti-phospho Histone H2A.X Ser 139 (Cell Signaling #2577, 1:400); rabbit anti-phospho CASC5/KNL1 Thr943/Thr1155 D8D4N (Cell Signaling #40758, 1:400); ACA (Antibodies inc. #15-235, 1:500); and incubated overnight at 4° C. Cells were rinsed three times with PBS + 0.05% Triton X-100 and then incubated with secondary antibodies: Alexa Fluor 568 Goat anti-rabbit (Thermo Fisher Scientific #A11011, 1:300) and Alexa Fluor 647 Goat anti-human (Thermo Fisher Scientific #A21445, 1:300) for 1 hour at room temperature. The coverslips were then washed 3 times in PBS containing 0.05% Triton X-100. Cells were then incubated with 1 μg/ml DAPI (4,6-diamidino-2-phenylindole, Cell Signaling) for 5 min. After rinsing with PBS coverslips, coverslips were mounted onto glass slides with ProLong Glass mounting reagent (Invitrogen, cat. no. P36930). Images were acquired using a Zeiss LSM980 laser-scanning microscope (Oberkochen) with ZEN 2.3 SP1 software. Image analysis was performed Image J2 v2.9.0 software.

### Fluorescence Quantification of Phospho-KNL1 Confocal Images

Image quantification was performed using Fiji (Image J2 v2.9.0) software. Briefly, regions of interest were defined by identifying mitotic cells and drawing an area centered on the DAPI stain. Background values measured in an adjacent region were subtracted, and the resulting corrected total cell fluorescence (CTCF) was calculated using the formula; CTCF = (Integrated density) – (Area of selected cell x Mean fluorescence of background). Each individual mitotic cell CTCF was plotted in a dot plot and CIRc-004 treated cells were compared vehicle treated cells.

### Live-cell imaging

Live cell imaging (LCI) was performed on a Ti2 inverted microscope fitted with a CSU-W1 spinning disk system (Nikon). To perform LCI, NCI-H1048 cells were stably expressed with both mCherry-PCNA and PGK-H2B-eGFP (Addgene #21210). Two rounds of FACS sorting were performed to obtain a pure population of eGFP and mCherry double-positive cells. To capture mitotic events for these experiments, we focused our live cell imaging experiments on H2B-eGFP. Cells were seeded onto 35 mm µ-Dish (iBiDi, #80137) at 500,000 cells/mL and allowed to attach and grow for 48 hours. CIRc-004 (20 nM) was added 2 hours prior to LCI. Z-stacks (+6 (above) and −4 (below) planes at 0.5-μm spacing) were captured every 5 minutes for 10 hours, using a Zyla 4.2 sCMOS camera (Andor), and a 20×/0.95 NA Plan Apochromat Lambda objective with the correction collar set to 0.17. An environmental enclosure was used to maintain cell culture conditions (37°C and humidified 5% CO2) for all live-cell confocal imaging. Images were analyzed on NIS Elements Viewer 4.2 (Nikon). The timing of mitosis was measured from DNA condensation (beginning timepoint) to cytokinesis completion (end timepoint).

### CRISPR/Cas9 Base Editor Screens with the Cyclin A/B RxL inhibitor

In order to establish the adenosine or cytosine base editor Cas9 expressing cell line for the screen with the tiling library, NCI-H1048 cells were transduced with pRDA_867 (A>G base editor Cas9) and pRDB_092 (C>T base editor Cas9), respectively. Successfully transduced cells were selected with blasticidin (5 µg/ml) to make stable adenosine or cytosine base editor cell lines.

The base editor library (CP1985) was custom designed using the Broad Institute’s CRISPR base editor library designer Beagle (https://portals.broadinstitute.org/gppx/beagle/screener) to include a total of 2802 sgRNAs tiling *CCNB1,* NM_031966.4 (572 sgRNAs), *CCNA2* NM_001237.5 (577 sgRNAs), *CDK2* NM_001798.5 (476 sgRNAs) and *CDC20* NM_001255.3 (897 sgRNAs). As controls the library contained 56 guides against essential genes (*EEF2*, *DBR1*, *PLK1*, *KIF11*, *GAPDH*, *PCNA*, *RPA3*, *RPL3*, *POLR2B* and *PSMB1*), 112 guides against intergenic sites and 112 guides as non-targeting controls.

First a pilot experiment was performed to empirically determine the lentiviral titer needed for a goal multiplicity of infection (MOI) of 0.3-0.6. Cells were mixed with polybrene and increasing amounts of the lentiviral CP1985 sgRNA library and plated in 1 mL aliquots onto 12 well plates and centrifuged at 931 x g [Allegra® X-15R Centrifuge (Beckman Coulter), rotor SX4750A, 2000 rpm], for 2 hours at 30°C. In parallel, 2 × 10^6^ cells were also centrifuged under same conditions but without lentivirus as a control for puromycin selection (Mock). ∼16 hours later (day 1), the cells were collected, pooled, and centrifuged again at 448 x g to remove the lentivirus and polybrene, and the cell pellet was resuspended in complete media and plated into 15 cm tissue culture treated plates at 0.4 × 10^6^ cells/mL or in 6 well plates at the same density for the control (Mock) cells. The cells were then cultured for 72 hours at which time (day 4) the cells were counted and plated at 0.4 × 10^6^ cells/mL in 15 cm tissue culture treated plates with fresh media containing puromycin (0.5 µg/ml) to select for puromycin-transduced cells. After 72 hours (day 7), cells were counted using the Vi-Cell XR Cell Counter and the MOI was calculated using the following equation: (# of puromycin-resistant cells infected with the sgRNA library/# total cells surviving without puromycin after infection with the sgRNA library) – (# of puromycin-resistant mock-infected cells/ # total mock-infected cells). These experiments demonstrated that 25 µl/mL of CP1985 virus was needed to achieve at MOI of ∼0.3-0.6.

For the screen, adenosine or cytosine base editor Cas9 expressing NCI-H1048 cell lines were counted using the Vi-Cell Counter on day 0. Two replicates of 2.4 × 10^7^ cells (which would yield representation of at least 2000 cells/sgRNA if MOI=0.3) were pelleted and resuspended at 2 × 10^6^ cells/mL in 1 mL of complete media containing 25 µl/mL of the pooled sgRNA library (CP1985) lentivirus and 8 µg/mL polybrene. A parallel experiment was performed on day 4 to determine the actual MOI of the screen as described above, which showed that the MOI of the screen was ∼0.6. On the same day (day 7), puromycin containing media was exchanged with fresh complete media not containing puromycin for all cells in the screen.

On day 10, cells from each replicate were pooled, counted, and then split into two additional replicates (so four replicates in total). 6 × 10^6^ cells from each replicate were harvested for an early timepoint (ETP) for representation of 2000 cells/sgRNA. This was done by washing cells in PBS and freezing cell pellets at −80° C for genomic DNA isolation. Then the remaining cells from each replicate were split into two treatment arms: 1) CIRc-004 (cyclin A/B RxL inhibitor, 200 nM); 2) CIRc-005 as a negative control and comparator (inactive enantiomer of CIRc-004, 200 nM) with 4.5 × 10^6^ cells in each treatment arm plated in 15 cm tissue culture plates at 0.2 × 10^6^ cells/mL in complete media containing the respective drugs listed above to maintain representation of 1500 cells/sgRNA. Mock-infected cells were maintained at the same density for all drug conditions and carried in parallel for the entire screen described below.

The media on the plates under drug selection was exchanged every 3 days with complete media containing fresh drug at the concentrations above. Treatment arms were counted and passaged when cells were ∼80% confluent and split back to a representation of 4.5 × 10^6^ cells at a density of 0.2 × 10^6^ cells/mL onto 15 cm tissue culture plates. This occurred for the CIRc-005 inactive enantiomer every 3 days as CIRc-005 had no effect on cellular proliferation. As the CIRc-004 exhibited strong cytotoxicity, this treatment arm only required media exchange every 3 days. When cells were passaged, the cells were pooled, counted, and replated maintaining 1500 cells/sgRNA. The remaining cells were divided in aliquots of 6 × 10^6^ cells, washed in PBS and cells pellets were frozen for genomic DNA isolation. The screen was terminated on Day 26 when there was an obvious growth advantage in the CIRc-004, arm infected with the sgRNA library relative to the mock infected cells treated with the same compounds.

Following completion of the screen, genomic DNA was isolated using the Qiagen Genomic DNA midi prep kit (cat. # 51183) and mini prep kit (cat. #51106) according to the manufacturer’s protocol. Raw Illumina reads were normalized between samples using: Log2[(sgRNA reads/total reads for sample) × 1e6) + 1]. Log Fold Change (LFC) in Fig. S8A-B was calculated by subtracting the plasmid DNA (pDNA) log normalized value from the CIRc-005 late timepoint (LTP, day 26) log normalized value and this was used for replicate reproducibility and dropout of essential sgRNA vs. negative-control sgRNAs. To calculate the z-scored LFC in Fig. 4B and S8C-E, we first subtracted the log normalized value of the negative enantiomer control (CIRc-005) at the LTP (day 26) from the log normalized value of the cyclin A/B RxL inhibitor (CIRc-004) at the LTP (day 26). This LFC value (CIRc-004 minus CIRc-005) for each sgRNA was z-scored, a transformation which factors in the performance of the control guides, using the following equation: z-score = (x - μ)/σ, where x, μ, and σ correspond to LFC of an individual guide, the mean LFC of all negative control guides, and standard deviation of all control guides, respectively. These z-scored LFC values for each sgRNA were then matched to the annotated CP1985 library where Beagle (Broad Institute) was used to predict the expected nucleotide edit and plotted for each tiled gene as indicated in each figure panel listed above. The averaged data from the 4 replicates were used for all analyses shown in the manuscript.

### CRISPR/Cas9 Base Editor Screen with the Cyclin A/B RxL inhibitor with Focused *CCNB1* Target Validation

To validate the exact mutations enriched after CIRc-004 treatment in *CCNB1* corresponding to the amino acids 117-285, which contained several enriched *CCNB1* variants from our original base editor screen in Fig. 4B, we repeated the screen as in Fig. 4A at scale with a representation of 1500 cells/sgRNA, and at the late timepoint (LTP, day 26), mRNA was isolated from the cyclin A/B RxL inhibitor (CIRc-004) and enantiomer control (CIRc-005) treated arms using Quick-RNA™ Miniprep kit (Zymo Research, CA, USA) according to the manufacturer’s instructions. RNA concentration was determined using the Nanodrop 8000 (Thermofisher Scientific). A cDNA library was synthesized using iScript Reverse Transcription Supermix for RT-qPCR (Biorad #1708841) according to the manufacturer’s instructions with 3000 ng of RNA. Target site on *CCNB1* was amplified by a 2-step PCR using KOD XTREME HOT START DNA POLYMERASE (Sigma Aldrich #71975-3) using 25% of the total cDNA reaction above. For the first PCR step, *CCNB1* #1F: 5’-GGAACGGCTGTTGGTTTCTG-3’ and *CCNB1* #1R: 5’-AAGTAAAAGGGGCCACAAGC-3’ primers were used to amplify a 1514 bp long region. Then 0.2 microliters of the first PCR reaction was used for a second round of PCR amplification using the following primers: #2F: 5’- CGCCTGAGCCTATTTTGGTTG-3’ and *CCNB1* #2R: 5’-CCATCTGTCTGATTTGGTGCTTAG-3’ to yield a 510 bp product. This final PCR product was column purified using QIAquick Gel Extraction kit (Qiagen #28704) and sent for amplicon sequencing using next-generation sequencing by the MGH DNA Core Facility and analyzed using MGH DNA Core’s complete amplicon sequencing pipeline for data analysis of >600 bp amplicons using the following CCNB1 NM_031966.4 refseq amplicon: CGCCTGAGCCTATTTTGGTTGATACTGCCTCTCCAAGCCCAATGGAAACATCTGGATGTGCCCCTGCAGAAGAAGACCTGTGTCAGGCTTTCTCTGATGTAATTCTTGCAGTAAATGATGTGGATGCAGAAGATGGAGCTGATCCAAACCTTTGTAGTGAATATGTGAAAGATATTTATGCTTATCTGAGACAACTTGAGGAAGAGCAAGCAGTCAGACCAAAATACCTACTGGGTCGGGAAGTCACTGGAAACATGAGAGCCATCCTAATTGACTGGCTAGTACAGGTTCAAATGAAATTCAGGTTGTTGCAGGAGACCATGTACATGACTGTCTCCATTATTGATCGGTTCATGCAGAATAATTGTGTGCCCAAGAAGATGCTGCAGCTGGTTGGTGTCACTGCCATGTTTATTGCAAGCAAATATGAAGAAATGTACCCTCCAGAAATTGGTGACTTTGCTTTTGTGACTGACAACACTTATACTAA GCACCAAATCAGACAGATG.

To analyze variants from the *CCNB1* refseq above, the percentage of mismatched reads at individual base positions were normalized to the percentage of sequencing reads to get the %variants at each position and then averaged across 2 independent experiments. Then the enrichment of %variants in the CIRc-004 treated arm was compared to the variants in the CIRc-005 treated arm to determine the fold change in CIRc-004 vs CIRc-005 at the LTP.

### AlphaFold Protein Co-Folding Prediction

Co-folding of proteins were done by AlphaFold 2 implemented in ColabFold^92^. Default parameters were used to predict their relative positions. Specifically, those parameters are: msa_mode (MMseqs2_UniRef_Environmental), pair_mode (unpaired_paired), model_type (auto), number of cycles (3), recycle_early_stop_tolerance(auto), relax_max_iterations (200), and pairing_strategy (greedy). The resulting co-folding structure along with the PAE files were analyzed using ChimeraX software^93^. The command “alphafold contacts /A to /B distance 3” was employed to identify residues potentially interacting within a putative predicted distance of less than 3.0 Å.

### Generation of Endogenous *CCNB1* Mutations using Cas9 Base Editing

sgRNA sequences targeting specific base edits in *CCNB1* were selected from the CP1985 sgRNA library based on z-scored LFC enrichment after CIRc-004 treatment in our primary screens, synthesized by IDT technologies and cloned using the same approach as sgRNA oligo cloning described above. These sgRNAs were integrated into the GFP-expressing pMV_AA013 lentiviral vector (Broad Institute) using sgRNA Cloning method described above. NCI-H1048 cells, expressing either pRDA_867 (A>G base editor Cas9) or pRDB_092 (C>T base editor Cas9), were transduced with the pMV_AA013 plasmid carrying the sgRNA sequences and subsequently subjected to puromycin selection to establish stable edited cell lines.

The following sgRNA oligos were used for A>G edits:

Non-targeting sense (5’- CAACGTCGCGAACGTCGTAT −3’)

Non-targeting anti-sense (5’- ATACGACGTTCGCGACGTTG −3’)

Human *CCNB1* Tyr170His sense (5’- ACATATTCACTACAAAGGTT −3’)

Human *CCNB1* Tyr170His antisense (5’- AACCTTTGTAGTGAATATGT −3’)

Human *CCNB1* Tyr177Cys sense (5’- TATGCTTATCTGAGACAACT −3’)

Human *CCNB1* Tyr177Cys antisense (5’- AGTTGTCTCAGATAAGCATA −3’)

Human *CCNB1* Gln213Arg/Met241Val sense (5’- GGTTCAAATGAAATTCAGGT −3’)

Human *CCNB1* Gln213Arg/Met241Val antisense (5’- ACCTGAATTTCATTTGAACC −3’)

Human *CCNB1* Val227Ala/Ser228Pro sense (5’- ATGGAGACAGTCATGTACAT −3’)

Human *CCNB1* Val227Ala/Ser228Pro antisense (5’- ATGTACATGACTGTCTCCAT −3’)

The following sgRNA oligos were used for C>T edits:

Non-targeting sense (5’- CAACGTCGCGAACGTCGTAT −3’)

Non-targeting anti-sense (5’- ATACGACGTTCGCGACGTTG −3’)

Human *CCNB1* Glu169Lys sense (5’- CATATTCACTACAAAGGTTT −3’)

Human *CCNB1* Glu169Lys antisense (5’- AAACCTTTGTAGTGAATATG −3’)

Human *CCNB1* Glu314Lys sense (5’- TGTACCTCTCCAATCTTAGA −3’)

Human *CCNB1* Glu314Lys antisense (5’- TCTAAGATTGGAGAGGTACA −3’)

Human *CCNB1* Ala346Thr/ Gly347Arg sense (5’- CTCCTGCTGCAATTTGAGAA −3’)

Human *CCNB1* Ala346Thr/ Gly347Arg antisense (5’- TTCTCAAATTGCAGCAGGAG −3’)

Human *CCNB1* Glu374Lys sense (5’- GAGATTCTTCAGTATATGAC −3’)

Human *CCNB1* Glu374Lys antisense (5’- GTCATATACTGAAGAATCTC −3’)

### FACS based competition assay

The GFP-expressing stable base-edited cells were combined with their parental base editor counterpart cells (without the sgRNA and GFP) at a ratio of 1:4 with the exact ratio confirmed by FACS analysis for GFP and seeded at a density of 0.1 × 10^6^ cells/ml in 10 cm dishes. The cells were cultured in the presence of the cyclin A/B RxL inhibitor (CIRc-004) at concentrations of 20 nM or 200 nM or DMSO. Cells were collected for FACS analysis on days 6 and 13. Following each time point, cells were replated and continued to be cultured in the presence of the drug, with fresh drug added every 3 days. On day 13, the remaining cells post-FACS were collected, and genomic DNA was extracted from the harvested cells. Data from the FACS-based competition assay at day 13 is shown in Fig. 4E.

To verify the exact mutations in CCNB1 made by the base-editing sgRNAs and to determine if CIRc-004 caused enrichment of these mutations, genomic DNA from the competition assay above at day 13 was harvested andPCR amplification of the *CCNB1* target region (forward #3F: 5’- GCCCAATGGAAACATCTGGATG −3’, reverse #3R: 5’- GCGATCTCTTAAGAAATGCTGCCC −3’) was performed followed by next-generation sequencing using CRISPR amplicon sequencing conducted at the MGH DNA Core Facility as previously described^94^.

### Mass Spectrometry Cyclin B1 CIRc-004-Dependent Interactome Sample Preparation

To identify proteins that are modulated by the cyclin A/B RxL inhibitor CIRc-004, we performed cyclin B immunoprecipitation followed by mass spectrometry (IP-MS) in NCI-H1048 cells treated with CIRc-004 (50 nM) or CIRc-005 (50 nM) or DMSO for 2 hours. The Cyclin B1 IP was performed as described above with the following modifications: 10 mg cell extract was rotated overnight at 4°C with 10 μg anti-cyclin B antibodies. After Cyclin B1 IP, beads were washed with PBS before freezing at −80 °C. Dry beads were resuspended with two hundred microliters of Smart digest buffer (ThermoFisher Scientific, 60113-101) before adding three microliters of Smart trypsin. Beads were digested in a Thermo block (Fischer Scientific) for 1 hour 30 minutes at 70° C with 1400 rpm (∼250 RCF) agitation. Trypsin digestion was then stopped by adding trifluoroacetic acid at 1 % final concentration. After that, samples were desalted using SOLA HRP SPE cartridges (ThermoFischer Scientific; 60109-001) following manufacturer instructions. Purified peptide eluates were dried by vacuum centrifugation and re-suspended in 0.1% formic acid and stored at −20 °C until analysis.

### Mass Spectrometry-LC_MS/MS Analysis

LC-MS/MS analysis was performed using an Ultimate 3000 HPLC connected to an Orbitrap Ascend Tribrid instrument (Thermo Fisher) and interfaced with an EASY-spray source. 0.5% of the tryptic peptides were loaded onto an AcclaimPepMap100 trap column (100µm x 2cm, PN164750; ThermoFisher) and separated on a 50cm EasySpray column (ES903, ThermoFisher) using a 60 min linear gradient from 2 to 35% B (acetonitrile, 0.1% formic acid) and at 250nl/min flow rate. Both trap and column were kept at 50C. The Orbitrap Ascend was operated in Data-Independent mode (DIA) to automatically switch between MS1 and MS2, with minor changes from previously described method^95–97^. In brief, MS1 scans were collected in the Orbitrap at a resolving power of 45K at m/z 200 over m/z range of 350 – 1650m/z. The MS1 normalised AGC was set at 125% (5e5ions) with a maximum injection time of 91 ms and a RF lens at 30%. DIA MS2 scans were then acquired using the tMSn scan function at 30K orbitrap resolution over 40 scan windows with variable width, with a normalized AGC target of 1000%, maximum injection time set to auto and a 30 % collision energy.

### Mass Spectrometry-Data Analysis

DIA-NN software (V8.1) was used to analyse the raw data in library-free mode and using the recommended default settings^97^. Briefly, a library was created from human UniProt SwissProt database (downloaded 20220722 containing 20,386 sequences) using deep learning. Trypsin was selected as the enzyme (1 missed cleavage), N-term M excision and Methionine were enabled. The identification and quantification of raw data were performed against the in-silico library applying 1% FDR at precursor level and match between runs (MBR). Perseus (v1.6.2.2) was used to further analyse the DIA-NN protein group output. Protein intensities were log2 transformed and normalised by median subtraction (by column) after applying a pre-processing filter based on valid numbers (3 out of 3 in at least one group). Subsequently, missing values were imputed following the normal distribution down shifted. A two-sample student t-test combined with multiple testing correction using Permutation-based FDR (5%) was applied. For ranking the proteins based on intensity, the average intensity was calculated for all values in the 3 treatment groups, log10 transform and rank.

The raw data included in this study will be submitted to the Proteome eXchange Consortium via the PRIDE (PRoteomics IDEntifications - https://www.ebi.ac.uk/pride/) partner repository^98^.

### Animal Studies

All mouse experiments complied with National Institutes of Health guidelines. The patient-derived xenograft studies were approved and performed at the Dana-Farber Cancer Institute Animal Care and Use Committee. The NCI-H69 and NCI-H1048 xenograft studies were approved and performed at Labcorp Early Development Laboratories, Inc. and Pharmaron (Ningbo) Technology Development Co., Ltd., respectively. Housing conditions for mice include a 12 h/12 h day-night cycle where temperature is maintained at 72 Fahrenheit.

### SCLC Cell Line Xenograft Studies

For NCI-H69 studies, 5 × 10^6^ NCI-H69 cells were suspended in serum-free RPMI1640 media, combined with an equal volume of ECM gel (Sigma-Aldrich #E1270), and implanted subcutaneously into the flanks of 7-8 week old female Nude mice (Envigo, Hsd:Athymic Nude-*Foxn1^nu^*). After 13 days, when tumors were between 88-200 mm^3^ in size, mice were randomized and put into treatment groups of CIRc-028 or vehicle (5% DMSO, 10% Solutol HS15, in 85% D5W). Mice were treated daily with intravenous (IV) injections of 100 mg/kg CIRc-028 or vehicle for 14 days. Tumors were measured 3 times per week for the duration of the study. Tumors were collected 18 hours after the last dose, formalin fixed for 24 hours, and paraffin embedded for IHC analysis.

For NCI-H1048 studies, 3 × 10^6^ NCI-H1048 cells were suspended in a 1:1 mixture of HITES media and Matrigel (Corning #354234) and implanted subcutaneously into the flanks of 6-8 week old female BALB/c nude mice (GemPharmatech Biotech Co., Ltd., #D000521). After 16 days, when tumors were between 180-315 mm^3^ in size, mice were randomized and put into treatment groups. Treatment with CIRc-028 and vehicle was carried out as described above for NCI-H69 xenografts. Tumors were measured with calipers twice per week for the duration of the study.

### RNA-Sequencing and Principal Component Analysis

RNA-seq was performed on DFCI 393 and DFCI 402 PDX tumors that were used for the PDX studies as described previously^99^. Briefly, RNA was extracted using RNeasy mini kit (Qiagen #74106) including a DNase digestion step according to the manufacturer’s instructions. Total RNA was submitted to Novogene Inc. The libraries were prepared using NEBNext Ultra II non-stranded kit. Paired end 150bp sequencing was performed on Novaseq6000 sequencer using S4 flow cell. Sequencing reads were mapped to the hg38 genome by STAR. Principal component analysis (PCA) in Figure S11 was performed using the top 500 genes with the largest standard deviation of log transformed FPKM values and these genes were subjected to PCA using the removeBatchEffect function in the limma package (version 3.58.1) and prcomp function of R software (version 4.3.3). Log transformed sum of *ASCL1, NEUROD1, CHGA, CHGA, INSM1,* and *SYP* FPKM values were calculated as the neuroendocrine (NE) score for each sample.

### Patient Derived Xenografts (PDXs)

Patient derived xenografts were generated from either tumor biopsies or pleural effusions of SCLC patients. All patient samples were collected after obtaining written informed consent. PDX tumor, DFCI 393 originated from surgical biopsy at resistance from a patient that received prior chemotherapy and radiation. The surgical biopsy of DFCI 393 was directly implanted directly into the sub-renal capsule of NSG mice for expansion. DFCI oncopanel genomic DNA sequencing of DFCI 393 showed an *RB1* E280* mutation with Single copy deletion of *RB1* at 13q14.2, and a *TP53* K132N mutation with single copy deletion of *TP53* at 17p13.1 together showing biallelic inactivation of *RB1* and *TP53*. DFCI 402 was derived from pleural effusion of a patient that was treated with first-line Carboplatin/Etoposide and then 2^nd^-line Nivolumab and the pleural effusion was removed as part of routine clinical care. The pleural fluid from the effusion was immune depleted and implanted subcutaneously to make DFCI 402. DFCI oncopanel genomic DNA sequencing of DFCI 402 showed an RB1 R358* mutation with copy loss of 13 likely contributing to biallelic loss of RB1. It also showed a *TP53* G154Afs*16 mutation. RNA-sequencing of DFCI 393 and DFCI 402 with confirmation by immunoblot analysis showed that both PDX models are neuroendocrine-high SCLCs and DFCI 393 dominantly expresses ASCL1 while DFCI 402 expresses both ASCL1 and NEUROD1.

### PDX Model Generation

Studies and procedures were performed at Dana Farber Institute in accordance with Institutional Animal Care and Use Committee (IACUC) guidelines in an AAALAC accredited vivarium. PDX models were generated using 7- to 8-week-old female NSG (NOD.Cg-Prkdcscid Il2rgtm1WjI/SzJ) mice purchased from Jackson Laboratories. Following initial implantation of patient samples, all PDX tumors were expanded and passaged continuously in NSG mice as subcutaneous tumors.

### In Vivo Efficacy, Pharmacodynamic, and Pharmacokinetic Studies in SCLC PDX Models

Female NSG mice, 7-8 weeks old were used for all in vivo studies. PDX tumor fragments were dipped in Matrigel and implanted subcutaneously in NSG mice. Tumor growth was measured twice weekly when palpable. Once tumor volumes reached approximately 200 ± 50 mm^3^, mice were randomized using Studylog software (San Francisco, CA) into the treatment groups as: vehicle (30% PEG400, 20% Solutol HS15, 50% Phosal 53 MCT) or CIR-7321 (100 mg/kg, Circle Pharma). In the DFCI-393 model, CIRc-014, was administered orally (PO)_three times a day (TID) for 28 days continuously and in the DFCI-402 model it was administered orally (PO) twice a day (BID) for 28 days continuously. In the DFCI-393 model, tumor samples were collected on day 28 after 8 hours and for DFCI-402, tumors were monitored for re-growth after treatment withdrawal. Animals were euthanized if tumor was necrotic/ulcerated or if tumor volume exceeded 2000 mm^3^. For pharmacodynamic (PD) study in DFCI 393, mice bearing PDX tumors were treated with either vehicle or CIRc-014 (100 mg/kg PO TID) for 4 days. All tumor samples were collected 8 hours after the morning dose on the last day of treatment. Tumor samples were fixed and snap frozen for further downstream IHC analysis (see below). Plasma timepoint sampling was conducted during the last dose of DFCI 393 pharmacodynamic study in CIRc-014 treatment group (N=6). Animals 1-3 were bled at 0.5 and 2 hours and animals 4-6, 1 and 4 hours respectively followed by terminal bleed via cardiac puncture at 8 hours. Blood was collected in K2EDTA tubes, centrifuged and plasma collected and sent for analysis.

### CIRc-014 Pharmacokinetic Plasma Analysis

The concentration of CIRc-014 was quantified in the mouse plasma samples via LC-MS/MS at Charles River Laboratories. CIRc-014 was used as a reference standard and glyburide (Sigma-Aldrich) was used as the internal standard. Blank matrix CD-1 mouse plasma, pooled/mixed gender with K_2_EDTA anticoagulant (BioIVT) was used to prepare calibration standards and quality control samples.

Sample extraction was performed on ice. A 10 µL aliquot of sample (calibration standards, quality control samples, blanks, and study samples) was added to a 96-well plate. 60 µL of internal standard solution (100 ng/mL glyburide in acetonitrile) was added to each well, except wells designated for matrix blanks to which 60 µL of acetonitrile was added. Samples were covered, vortexed, and centrifuged for 5 minutes. 50 µL of supernatant was transferred to a new 96 -well plate and the samples were diluted with 50 µL of MilliQ water before analysis. A Waters Acquity UPLC BEH C18, 50×2.1 mm (1.7 µm) column was used and maintained at 50°C during analysis. 5% acetonitrile/water with 0.1% formic acid was used as mobile phase A and 5% acetonitrile/methanol with 0.1% formic acid was used as mobile phase B. The chromatographic condition with flow rate at 0.8 mL/min was started with 60% B, ramped to 90% B in 1 min, and further increased to 95% B in 0.05 min and held for 0.25 min and the column was re-equilibrated at initial conditions for 0.4 min. Analyte and internal standard were detected using an Applied Biosystems Sciex API 6500 triple quadrupole mass spectrometer. The instrument was equipped with an electrospray ionization source (550°C) operated in the positive-ion mode. The analyte and internal standard were monitored in the multiple-reaction-monitoring (MRM) scan mode. The selected [M+H]+ precursor ions were m/z 962.4 for CIRc-014, and 494.1 for glyburide, and the product ions monitored were at m/z 423.4 and 169.0 for CIRc-014 and glyburide, respectively. Linear fitting with 1/x2 weighting for calibration range from 1.00 to 10000 ng/mL was used for data analysis. Analytical results for all samples met acceptance criteria.

### Immunohistochemistry Staining

Immunohistochemistry (IHC) staining was performed with a Bond RX Autostainer (Leica Biosystems) using a Polymer Refine Detection kit (Leica Biosystems). Four-micrometer thick sections were serially cut from formalin-fixed paraffin-embedded mouse tumor samples to perform single IHC studies using antibodies recognizing: Phospho-KNL1 (1:100, rabbit monoclonal antibody, Cell Signaling, Cat# 40758), and Cleaved Caspase-3 (1:400, rabbit monoclonal antibody, Cell Signaling Cat# 9664). The immunostained slides were scanned at (40X) using the PhenoImager HT instrument (Akoya Biosciences). For each slide, tumor areas were identified by a pathologist (YNL) and manually annotated using the HALO system. The number of positive and negative tumor cells was then determined using the HALO platform multiplex-IHC v3.2.5 algorithm (Indica lab) and the percentage of positive tumor cells was subsequently calculated.

### Statistical Analysis

For the CRISPR/Cas9 screen in Figure 3, the data was analyzed using the Apron, STARS, and Hypergeometric analysis algorithms on the GPP portal (https://portals.broadinstitute.org/gpp/screener/). Genes were considered “hits” if their q-value was <0.25 using the STARS analysis.

For all experiments with statistical data, the number of independent biological experiments is described in each figure legend. For immunoblots shown in Figs. 2L, 3C-D, G-I, 4H-J, 5B-E, G, J-K and Figs. S5G, S6D, K-L, S9C-G, S10B-D, at least three biological independent experiments were performed unless specified otherwise and representative immunoblots are shown. For dose response assays in Figs. 1F, 2E-H, 3E, 5F, 5H-I, and Figs. S2B-D, S4A-H, S6E-G, I, mean of three independent biological replicates are shown unless otherwise specified. For dose response assay shown in Figs. 3J-L and S7A-I, representative mean of two technical replicates are shown and each experiment was performed in three biological independent experiments. For live cell imaging experiment in Fig. 3A representative micrographs are shown from two biological independent experiments. For IF experiment in Figs. S6A and S10A, representative micrographs are shown from three biological independent experiments. For IHC experiments in Figs. 6C-D and 6K-L, representative micrographs are shown from at least five independent tumors per treatment group and the data are quantified in Figs. 6E-F and 6M-N.

For *in vivo* xenograft experiments in Figs. 6A-B and I-J, 2-way ANOVA was used to compare drug treatment vs. vehicle. For all other experiments, statistical significance was calculated using unpaired, two-tailed Students t-test. *p*-values were considered statistically significant if the *p*-value was <0.05. For all figures, * indicates *p*-value <0.05, ** indicates *p*-value <0.01, *** indicates and *p*-value <0.001, **** indicates and *p*-value <0.0001. Error bars represent SD unless otherwise indicated.

