## Supplementary data for "Cyclin A/B RxL Macrocyclic Inhibitors to Treat Cancers with High E2F Activity"

**Singh et al. Supplementary Data**

Figure S1

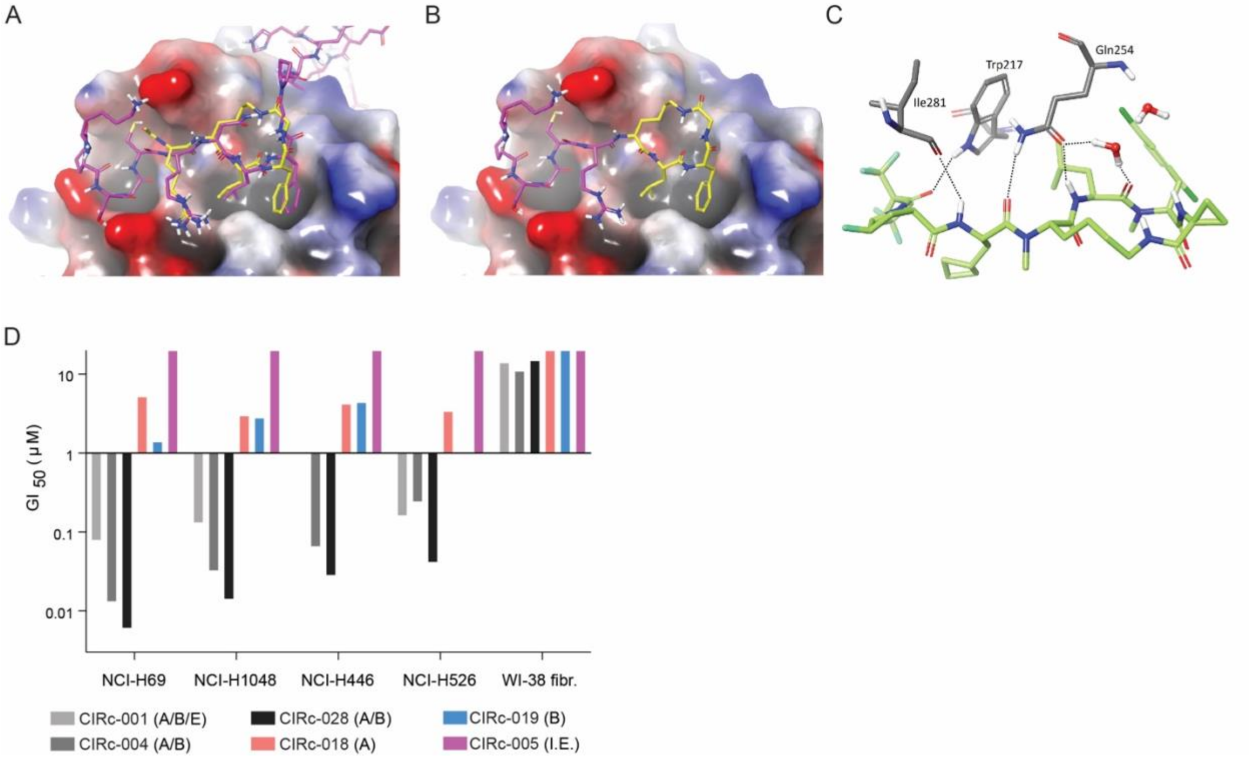


**Figure S1, related to Figure 1: A**) Overlay of cyclin A/Cdk2 complexes shown in surface representation with bound ligand at the RxL binding site, including a macrocycle (yellow, PDB: 1URC) and p27Kip1 (purple, PDB: 1JSU). **B)** Computational model of a lariat decapeptide macrocycle bound at the RxL binding site, based on the bound structures of p27Kip1 (purple) and lariat macrocycle (yellow), as a ligand alignment template for binding prediction. **C)** Detailed representation of modeled hydrogen bonds of CIRc-004 to Ile281, Trp217 and Gln 254 on cyclin A. **D)** Waterfall plot of GI50s for four SCLC and the non-transformed WI-38 fibroblast cell lines for compounds listed in **1C**.

Figure S2

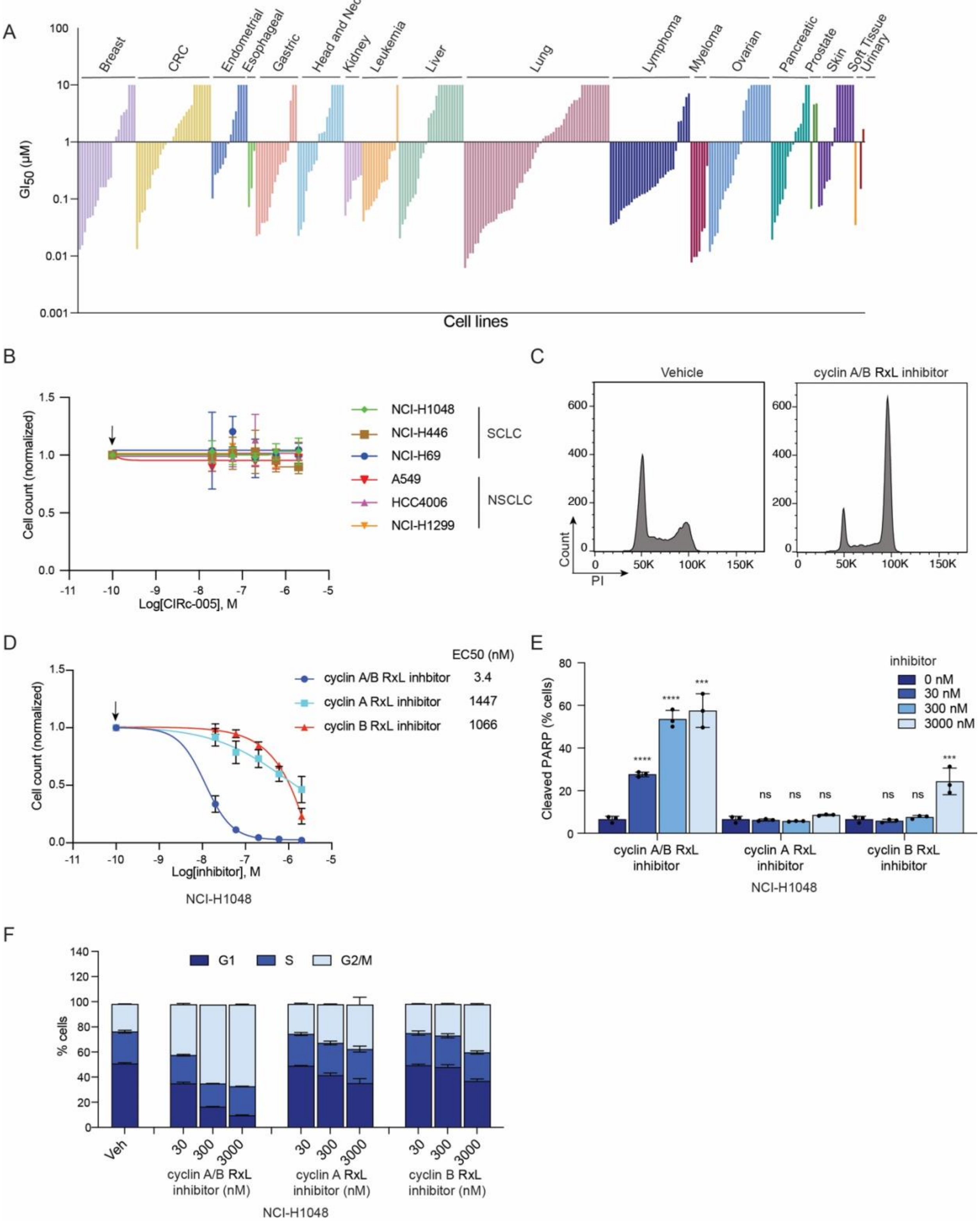
A
Breast
CRC
Endometrial
Esophageal
Gastric
Head and Neck
Kidney
Leukemia
Liver
Lung
Lymphoma
Myeloma
Ovarian
Pancreatic
Prostate
Skin
Soft Tissue
Urinary
GI50 (μM)
Cell lines
B
Cell count (normalized)
Log[CIRc-005], M
NCI-H1048
NCI-H446
NCI-H69
A549
HCC4006
NCI-H1299
SCLC
NSCLC
C
Vehicle
cyclin A/B RxL inhibitor
Count
PI
D
EC50 (nM)
cyclin A/B RxL inhbitor 3.4
cyclin A RxL inhibitor 1447
cyclin B RxL inhibitor 1066
Cell count (normalized)
Log[inhibitor], M
NCI-H1048
E
Cleaved PARP (% cells)
inhibitor
0 nM
30 nM
300 nM
3000 nM
cyclin A/B RxL inhibitor
cyclin A RxL inhibitor
cyclin B RxL inhibitor
NCI-H1048
F
G1
S
G2/M
% cells
Veh
cyclin A/B RxL inhibitor (nM)
cyclin A RxL inhibitor (nM)
cyclin B RxL inhibitor (nM)
NCI-H1048

**Figure S2, related to Figure 1: A**) $GI_{50}$ waterfall plot of cyclin RxL A/B/E inhibitor (CIRc-001) tested against Horizon Discovery cancer cell line panel. n=300 cell lines. **B)** Dose response assays of the indicated human SCLC cell lines (NCI-H1048, NCI-H446, and NCI-H69) and insensitive human NSCLC cell lines (A549, HCC4006, and NCI-H1299) treated for 6 days with increasing doses of the inactivate enantiomer of the cyclin A/B RxL inhibitor (CIRc-005). n=3 biological independent experiments. **C**) Representative histograms of flow cytometric analysis of propidium iodide (PI) stained NCI-H1048 cells treated with CIRc-004 (200 nM) or DMSO (vehicle) for 24 hours. **D)** Dose response assays of NCI-H1048 cells treated for 6 days with increasing doses of the selective cyclin A RxL inhibitor (CIRc-018), the selective cyclin B RxL inhibitor (CIRc-019), or the cyclin A/B RxL inhibitor (CIRc-004). **E)** Quantitation of cleaved PARP positive cells analyzed by flow cytometric analysis in NCI-H1048 cells treated for 3 days with the indicated doses of CIRc-018, CIRc-019, CIRc-004 or DMSO. n=3 biological independent experiments. **F)** Representative histograms of cell cycle distribution of NCI-H1048 cells treated with CIRc-018, CIRc-019, CIRc-004 or DMSO and then stained with PI. For **B**, **D**-**F**, n=3 biological independent experiments. Data are mean +/- SD. Arrows in **B, D** indicates DMSO-treated sample which was used for normalization. Statistical significance was calculated using unpaired, two-tailed students *t*-test. Where indicated, *=p<0,05, **=p<0.01, ***=p<0.001, ****=p<0.0001.

Figure S3

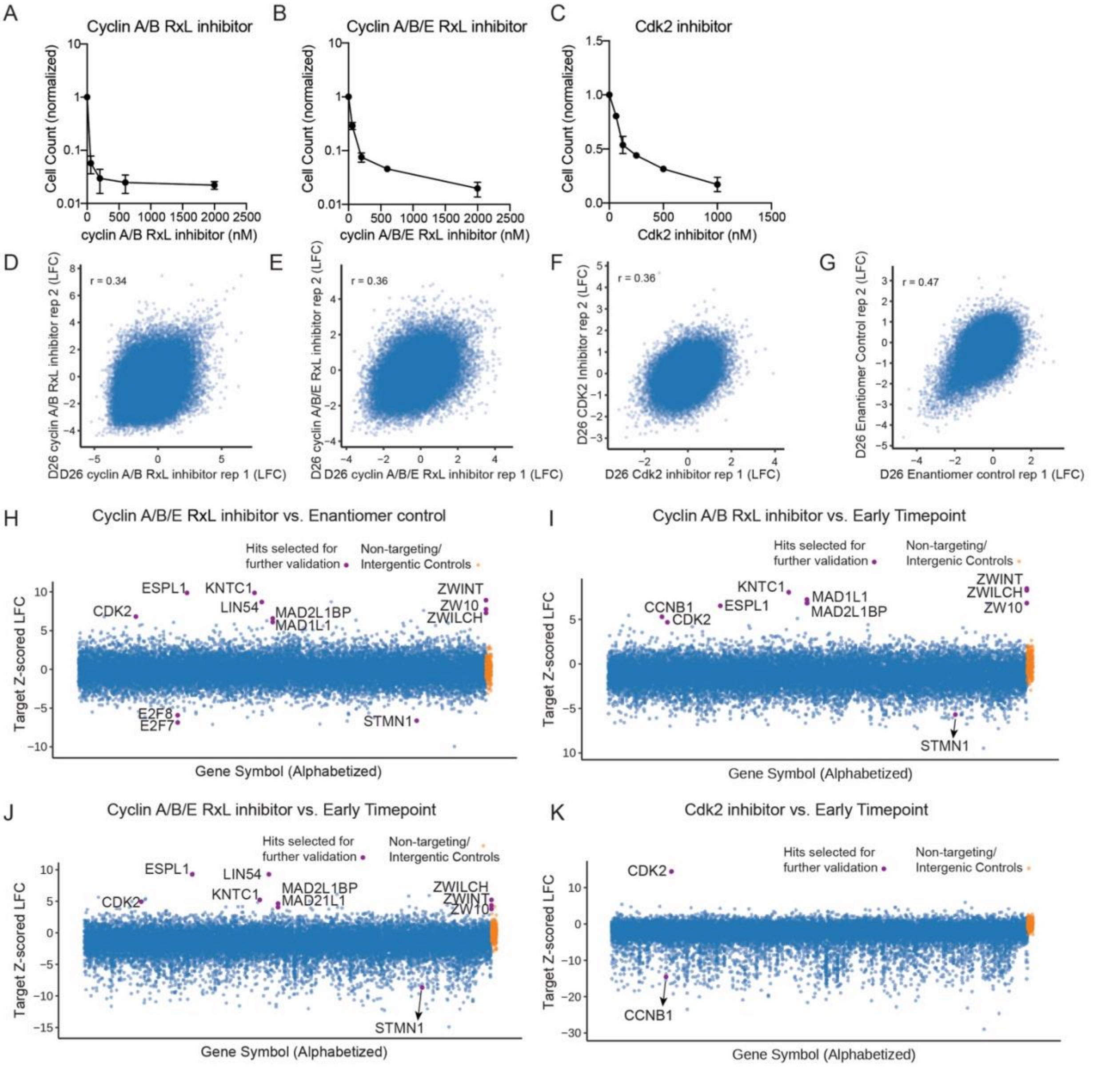
A
Cyclin A/B RxL inhibitor
Cell Count (normalized)
cyclin A/B RxL inhibitor (nM)
B
Cyclin A/B/E RxL inhibitor
Cell Count (normalized)
cyclin A/B/E RxL inhibitor (nM)
C
Cdk2 inhibitor
Cell Count (normalized)
Cdk2 inhibitor (nM)
D
r = 0.34
D26 cyclin A/B RxL inhibitor rep 2 (LFC)
D26 cyclin A/B RxL inhibitor rep 1 (LFC)
E
r = 0.36
D26 cyclin A/B/E RxL inhibitor rep 2 (LFC)
D26 cyclin A/B/E RxL inhibitor rep 1 (LFC)
F
r = 0.36
D26 CDK2 Inhibitor rep 2 (LFC)
D26 Cdk2 inhibitor rep 1 (LFC)
G
r = 0.47
D26 Enantiomer Control rep 2 (LFC)
D26 Enantiomer control rep 1 (LFC)
H
Cyclin A/B/E RxL inhibitor vs. Enantiomer control
Hits selected for further validation
Non-targeting/ Intergentic Controls
ESPL1
KNTC1
LIN54
CDK2
MAD2L1BP
MAD1L1
ZWINT
ZW10
ZWILCH
E2F8
E2F7
STMN1
Target Z-scored LFC
Gene Symbol (Alphabetized)
I
Cyclin A/B RxL inhibitor vs. Early Timepoint
Hits selected for further validation
Non-targeting/ Intergentic Controls
KNTC1
CCNB1
ESPL1
CDK2
MAD1L1
MAD2L1BP
ZWINT
ZWILCH
ZW10
STMN1
Target Z-scored LFC
Gene Symbol (Alphabetized)
J
Cyclin A/B/E RxL inhibitor vs. Early Timepoint
Hits selected for further validation
Non-targeting/ Intergentic Controls
ESPL1
LIN54
KNTC1
CDK2
MAD2L1BP
MAD21L1
ZWILCH
ZWINT
ZW10
STMN1
Target Z-scored LFC
Gene Symbol (Alphabetized)
K
Cdk2 inhibitor vs. Early Timepoint
Hits selected for further validation
Non-targeting/ Intergentic Controls
CDK2
CCNB1
Target Z-scored LFC
Gene Symbol (Alphabetized)

**Figure S3, related to Figure 2: A-C)** Dose response assays of NCI-H1048 cells treated with the indicated concentrations of the cyclin A/B RxL inhibitor (CIRc-004) (**A**), the cyclin A/B/E RxL inhibitor (CIRc-001) (**B**), the orthosteric Cdk2 inhibitor (PF-07104091) (**C**) for 6 days. **D-G**) Scatter plot of sgRNA abundance represented as Log Fold Change (LFC) comparing the 2 biological replicates for the **D)** cyclin A/B RxL inhibitor (CIRc-004), **E)** cyclin A/B/E RxL inhibitor (CIRc-001), **F)** orthosteric Cdk2 inhibitor (PF-07104091) and **G)** inactive enantiomer (CIRc-005). **H-K**) Top enriched and depleted hits determined by Apron analysis (Broad Institute) of the cyclin A/B/E RxL inhibitor (CIRc-001) at the end timepoint (day 26) relative to the CIRc-005 enantiomer negative control at the end timepoint (day 26) (**H**) and cyclin A/B RxL inhibitor (CIRc-004) (**I**), cyclin A/B/E RxL inhibitor (CIRc-001) (**J**), or the Cdk2 (PF-07104091) (**K**) at the end timepoint (day 26) all relative to the early timepoint prior to drug treatment (day 10). n=2 biological independent experiments.

Figure S4

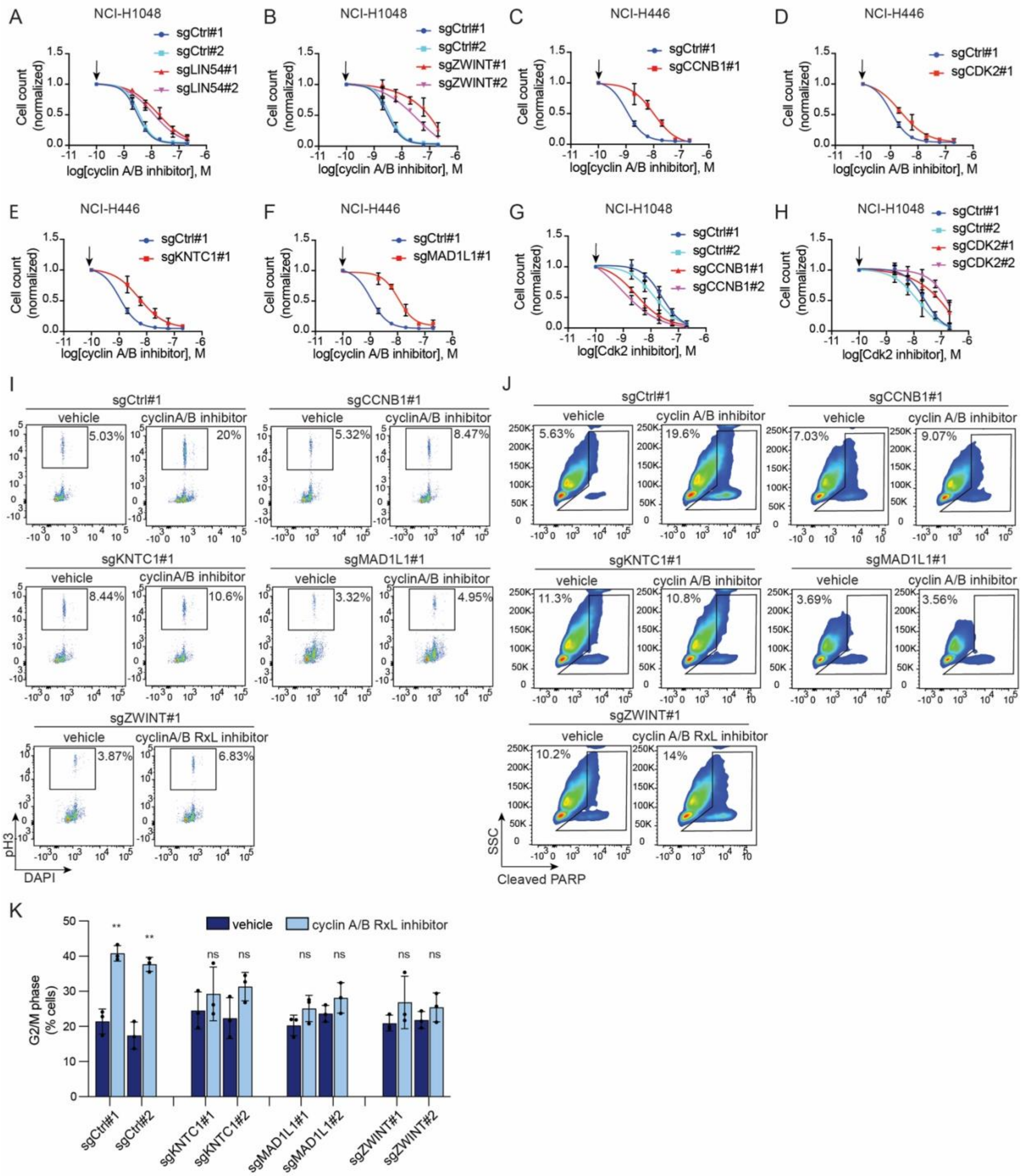

A
NCI-H1048
sgCtrl#1
sgCtrl#2
sgLIN54#1
sgLIN54#2
Cell count (normalized)
log[cyclin A/B inhibitor], M
B
NCI-H1048
sgCtrl#1
sgCtrl#2
sgZWINT#1
sgZWINT#2
C
NCI-H446
sgCtrl#1
sgCCNB1#1
D
NCI-H446
sgCtrl#1
sgCDK2#1
E
NCI-H446
sgCtrl#1
sgKNTC1#1
F
NCI-H446
sgCtrl#1
sgMAD1L1#1
G
NCI-H1048
sgCtrl#1
sgCtrl#2
sgCCNB1#1
sgCCNB1#2
log[Cdk2 inhibitor], M
H
NCI-H1048
sgCtrl#1
sgCtrl#2
sgCDK2#1
sgCDK2#2
I
sgCtrl#1
vehicle
cyclinA/B inhibitor
5.03%
20%
sgCCNB1#1
5.32%
8.47%
sgKNTC1#1
8.44%
10.6%
sgMAD1L1#1
3.32%
4.95%
sgZWINT#1
cyclinA/B RxL inhibitor
3.87%
6.83%
pH3
DAPI
J
sgCtrl#1
cyclin A/B inhibitor
5.63%
19.6%
sgCCNB1#1
7.03%
9.07%
sgKNTC1#1
11.3%
10.8%
sgMAD1L1#1
3.69%
3.56%
sgZWINT#1
cyclin A/B RxL inhibitor
10.2%
14%
SSC
Cleaved PARP
K
vehicle
cyclin A/B RxL inhibitor
G2/M phase (% cells)
sgCtrl#1
sgCtrl#2
sgKNTC1#1
sgKNTC1#2
sgMAD1L1#1
sgMAD1L1#2
sgZWINT#1
sgZWINT#2

**Figure S4, related to Figure 2:** Dose response assays of NCI-H1048 cells infected with two independent non-targeting sgRNAs (sgCtrl) or two independent sgRNAs against **A**) *LIN54*, **B**) *ZWINT* treated for 6 days with increasing doses of the cyclin A/B RxL inhibitor (CIRc-004). Dose response assays of NCI-H446 cells infected with sgCtrl or sgRNAs against **C**) *CCNB1*, **D**) *CDK*2, **E**) *KNTC1*, **F**) *MAD1L1*, treated for 6 days with increasing doses of CIRc-004. Dose response assays of NCI-H1048 cells infected with two non-targeting sgRNAs (sgCtrl) or two independent sgRNAs against **G**) *CCNB1*, **H**) *CDK2*, treated with treated for 6 days with increasing doses of the Cdk2 inhibitor (PF-07104091). **I**) Representative flow cytometric analysis from Fig. **2I** of phospho-histone H3 in NCI-H1048 cells infected with the indicated sgRNAs and treated with CIRc-004 at 20 nM or DMSO for 24 hours. **J**) Representative flow cytometric analysis from Fig. **2J** of cleaved PARP in NCI-H1048 cells infected with the indicated sgRNAs and treated with CIRc-004 at 20 nM or DMSO for 3 days. **K)** Quantitation of cells in G2/M phase of the cell cycle determined using PI staining of NCI-H1048 cells infected with indicated sgRNAs and then treated with CIRc-004 at 20 nM or DMSO for 24 hours. For **A-H, K**, n=3 biological independent experiments and data are mean +/– SD. Arrows in **A-H** indicates DMSO-treated sample which was used for normalization. Statistical significance in **K** was calculated using unpaired, two-tailed students *t*-test. Where indicated, *=p<0,05, **=p<0.01, ***=p<0.001, ****=p<0.0001.

### Figure S5

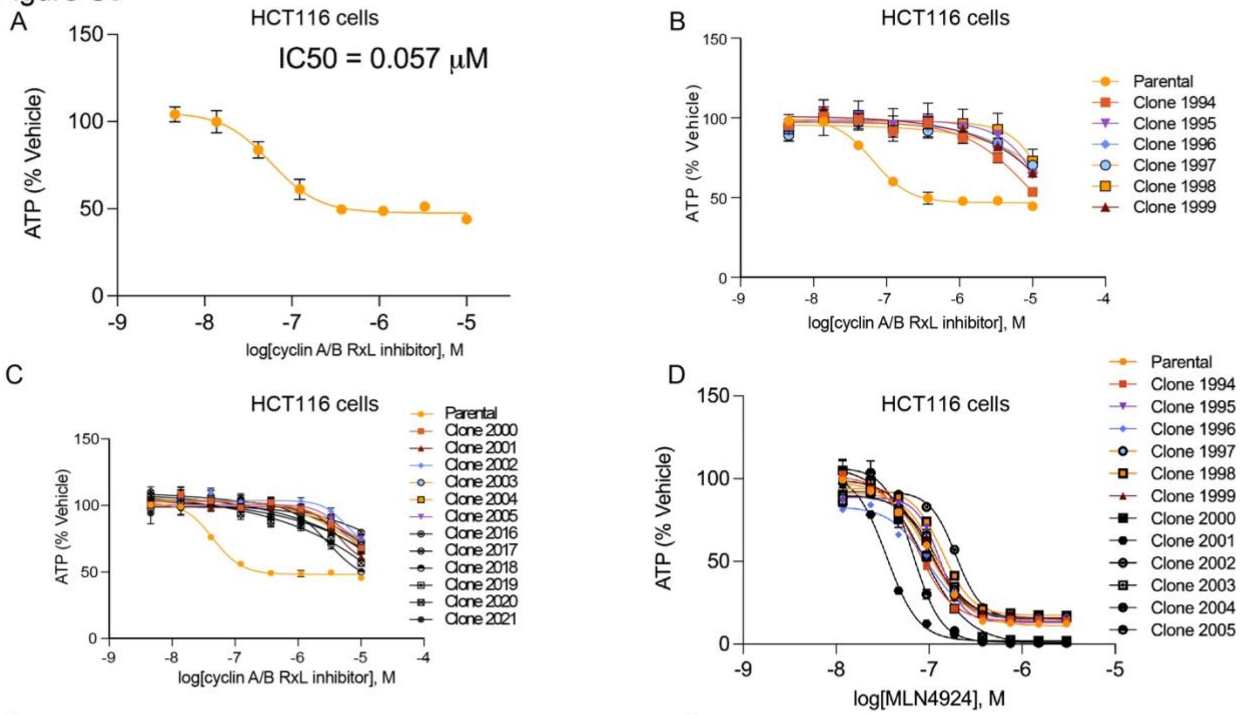


E

| Clone # | IAA | Barcode sequence |
|---|---|---|
| 1994 | - | GTGTCTGTCTGAGAGAGTGACACA |
| 1995 | - | GAGAGTGTGTGTGAGTCTCAGT |
| 1996 | - | CAGTCAGTCTCTGTCACAGAGACA |
| 1997 | - | CAGTCAGTCTCTGTCACAGAGACA |
| 1998 | - | GTGTGTGAGAGTGAGTGTCTGAGT |
| 1999 | - | GACACTGACACTCAGTGAGTCAGA |
| 2000 | + | CTGACAGTGACTCAGACTGTGAGT |
| 2001 | + | GTCTGTGAGTGAGTGTGAGTCACT |
| 2002 | + | CTGACAGAGTCTGTGAGACACACT |
| 2003 | + | GTGAGTGTGACTGTGAGTCAGTCA |
| 2004 | + | GTGTCTGTCTCTCAGTCTCACTCA |
| 2005 | + | GACAGTCAGTCAGAGTCTCAGTCT |
| 2016 | + | GTGTCTGACACTCAGTCTGAGAGT |
| 2017 | + | GTGTGTCTGTCTCTGTGAGTCACT |
| 2018 | + | GAGACAGTCACTGAGTGACTGACT |
| 2019 | + | CAGTCAGAGACAGACAGTCAGTCT |
| 2020 | + | CTGTCACTGAGTCTGTGTGACTGT |
| 2021 | + | CAGAGTCAGAGAGTCAGAGAGACA |

F

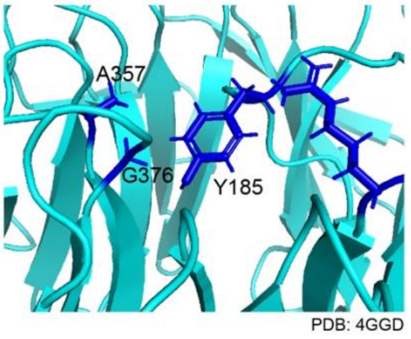


G

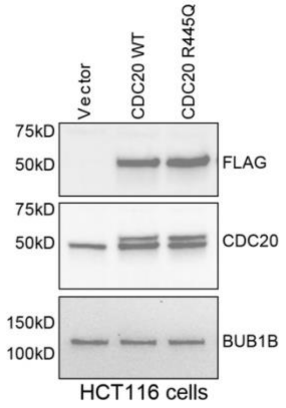


H

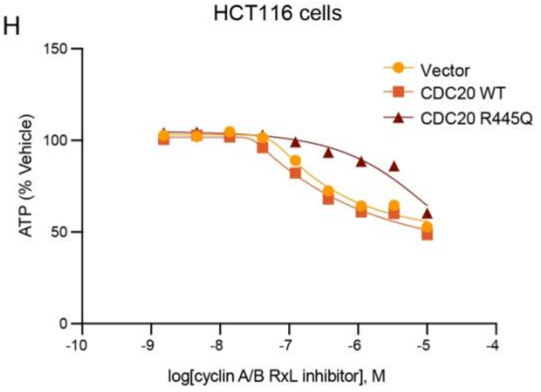

**Figure S5, related to Figure 2: A**) Dose response curve of iHCT116 cells treated with increasing doses of cyclin A/B RxL inhibitor (CIRc-004). **B**) Dose-response curves for six clones isolated from Mut-low iHCT116 cells and **C**) 12 clones from Mut-high iHCT116 cells treated with increasing doses of CIRc-004 for 3 days. **D**) Dose response curve of iHCT116 and different CIRc-004 resistant clones to MLN4924. For **A-D** n=2 technical replicates. Data are mean +/- SEM. **E**) Barcode sequences identified in 18 different CIRc-04 resistant clones, common sequences are marked in blue text. **F**) Mutated residues (blue) in CDC20 (PDB 4GGD) present in CIRc-004 resistant clones. **G**) Immunoblot analysis of iHCT116 cells stably expressing vector, Flag-CDC20 WT, or Flag-CDC20 R445Q mutant. **H**) Dose response assay iHCT-116 cells from **G** treated with increasing doses of CIRc-004. For **H**, n=3 technical replicates. Data are mean +/- SEM.

Figure S6

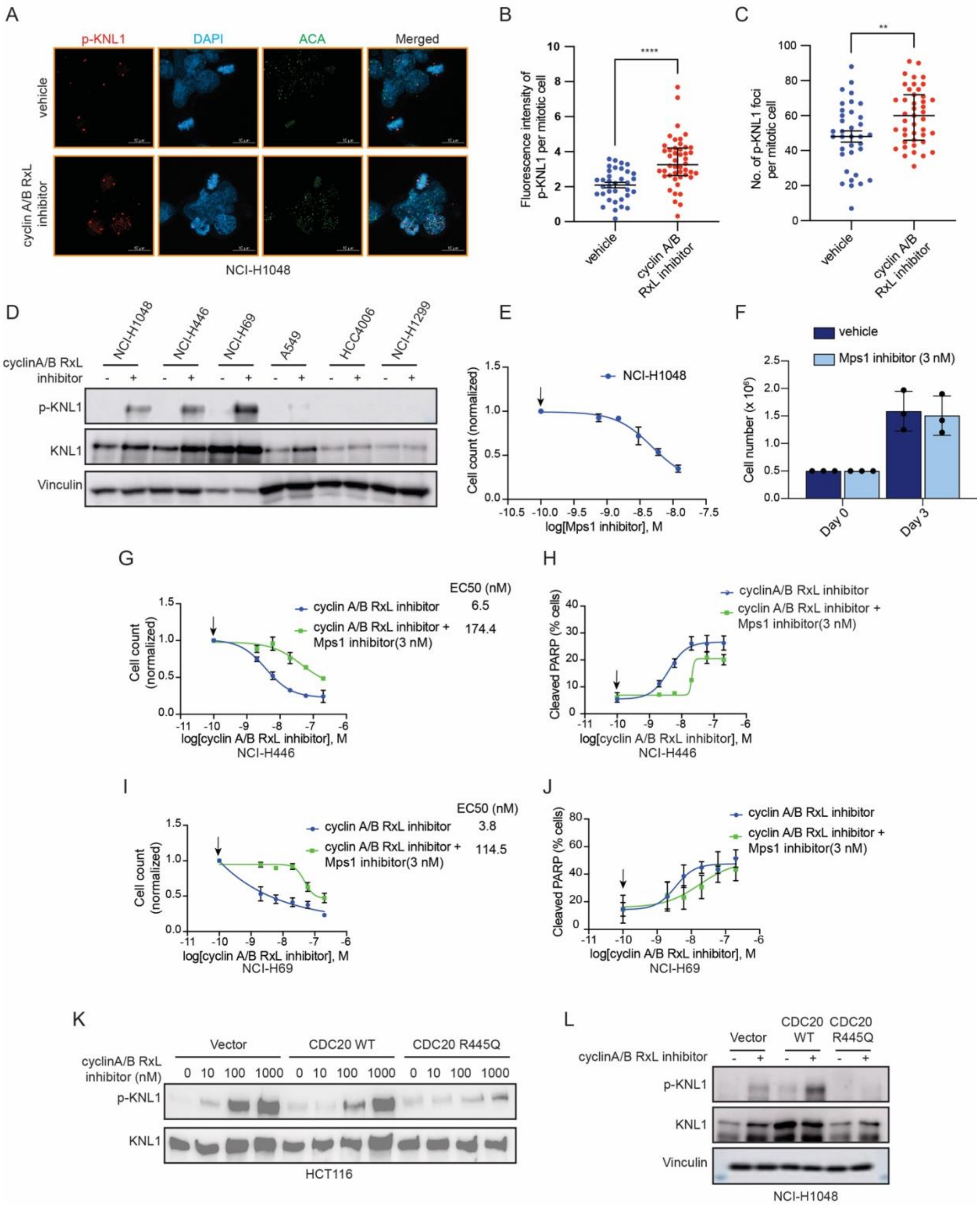
A
p-KNL1
DAPI
ACA
Merged
vehicle
cyclin A/B RxL inhibitor
NCI-H1048
B
Fluorescence intensity of p-KNL1 per mitotic cell
vehicle
cyclin A/B RxL inhibitor
C
No. of p-KNL1 foci per mitotic cell
D
NCI-H1048
NCI-H446
NCI-H69
A549
HCC4006
NCI-H1299
cyclinA/B RxL inhibitor
p-KNL1
KNL1
Vinculin
E
NCI-H1048
Cell count (normalized)
log[Mps1 inhibitor], M
F
vehicle
Mps1 inhibitor (3 nM)
Day 0
Day 3
G
EC50 (nM)
cyclin A/B RxL inhibitor 6.5
cyclin A/B RxL inhibitor + Mps1 inhibitor(3 nM) 174.4
log[cyclin A/B RxL inhibitor], M
NCI-H446
H
cyclinA/B RxL inhibitor
cyclin A/B RxL inhibitor + Mps1 inhibitor(3 nM)
Cleaved PARP (% cells)
I
EC50 (nM)
cyclin A/B RxL inhibitor 3.8
cyclin A/B RxL inhibitor + Mps1 inhibitor(3 nM) 114.5
NCI-H69
J
K
Vector
CDC20 WT
CDC20 R445Q
cyclinA/B RxL inhibitor (nM)
0 10 100 1000
HCT116
L
NCI-H1048

**Figure S6, related to Figure 3: A)** Representative phospho-KNL1, DAPI and anti-centromere antibody (ACA) confocal microscopy images of NCI-H1048 cells treated with cyclin A/B RxL inhibitor (CIRc-004) at 20 nM or DMSO for 24 hours. Magnification=63x, scale bar=10 µm. Dot plot measuring fluorescence intensity of phospho-KNL1 per mitotic cell (**B**) or number of phospho-KNL1 foci per mitotic cell (**C**) of NCI-H1048 cells from A. n=45 mitotic cells from 3 independent biological experiments. Statistical significance in **B-C** was calculated using unpaired, two-tailed students *t*-test. Where indicated, *=p<0,05, **=p<0.01, ***=p<0.001, ****=p<0.0001.**D)** Immunoblot analysis of the indicated human SCLC cell lines (NCI-H1048, NCI-H446, and NCI-H69) and insensitive human NSCLC cell lines (A549, HCC4006, and NCI-H1299) treated for 24 hours with CIRc-004 at 200 nM or DMSO. Note, this immunoblot contains the same lysates from Fig. **3C** and is included to ensure that the differences in phospho-KNL1 levels after CIRc-004 treatment between cell lines in Fig. **3C** are not a consequence of being loaded on independent immunoblots. **E)** Dose response assay of NCI-H1048 cells treated with increasing concentrations (0 nM, 1.5 nM, 3 nM, 6 nM, 12 nM) of the Mps1 inhibitor (BAY-1217389). n=2 biological independent experiments. Note that 3 nM of BAY-1217389 effectively blocked phosphorylation of the MPS substrate KNL1 (see Fig. **3D**) without blocking cellular proliferation and hence 3 nM of BAY-1217389 was used for all rescue experiments in Figure **3**. **F)** The raw cell counts of the dose response with BAY-1217389 from **E** showing that NCI-H1048 cells treated with BAY-1217389 at 3 nM proliferated over the 3-day dose response assay similar to the DMSO untreated control. **G-J)** Dose response assays of **G**) NCI-H446 or **I**) NCI-H69 cells and non-linear regression curves of cleaved PARP FACS analysis of **H**) NCI-H446 or **J**) NCI-H69 cells treated with increasing concentrations of the cyclin A/B RxL inhibitor (CIRc-004) in presence or absence of the Mps1 inhibitor (BAY-1217389) at 3 nM for 3 days. For **E-J** data are mean +/–SD. Arrows in **E**, **G-J** indicates DMSO-treated sample which was used for normalization. **K,L**) Immunoblot analysis of **K**) HCT116 or **L**) NCI-H1048 cells stably expressing vector, Flag-CDC20 WT, or Flag-CDC20 R445Q mutant treated with CIRc-004 with indicated concentrations for 24 hours. For **F, H-L**, n=3 biological independent experiments. For **E** and **G**, n=2 biological independent experiments.

Figure S7

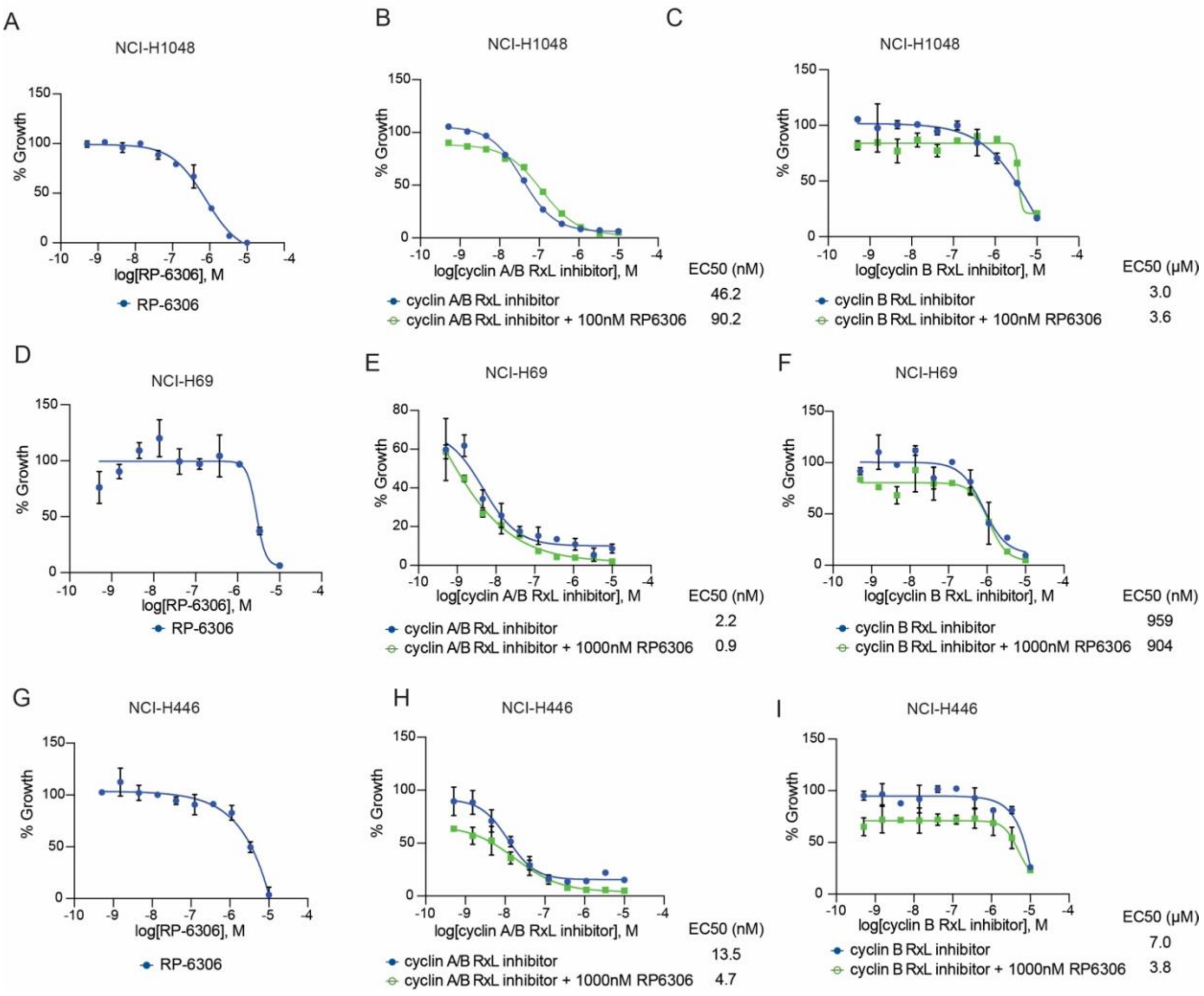


**Figure S7, related to Figure 3:** Dose response assays of NCI-H1048 (**A-C**), NCI-H69 (**D-F**) or NCI-H446 (**G-I**) cells treated with RP-6306 (Myt1 inhibitor) alone (**A, D, G**) or in combination with the cyclin A/B RxL inhibitor (CIRc-004) (**B, E, H**) or the cyclin B RxL inhibitor (CIRc-019) (**C, F, I**) for 5 days. For **A**-**I**, data are mean +/- SD of two technical replicates. n=3 biological independent experiments.

Figure S8

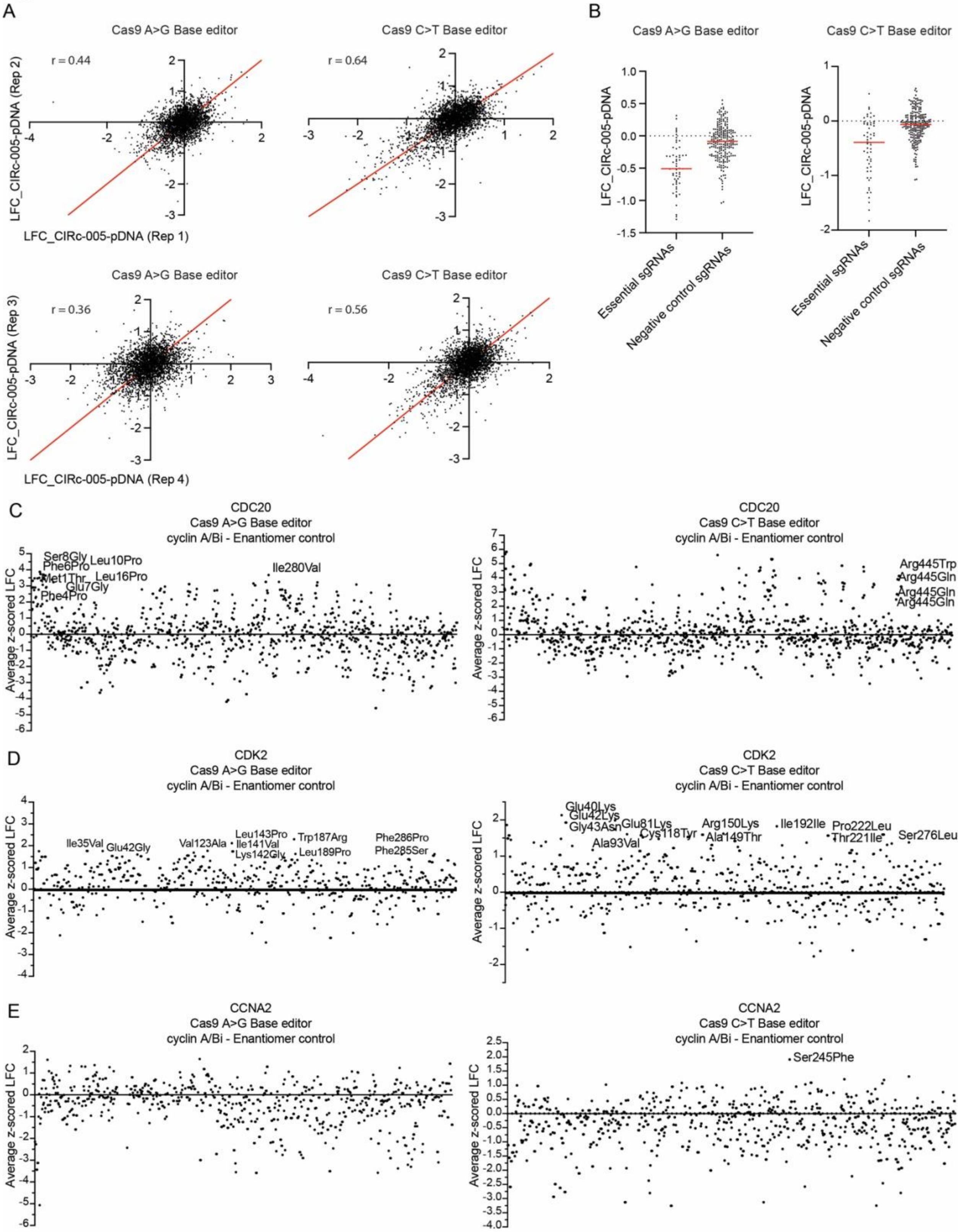
A
Cas9 A>G Base editor
Cas9 C>T Base editor
r = 0.44
r = 0.64
r = 0.36
r = 0.56
LFC_CIRc-005-pDNA (Rep 2)
LFC_CIRc-005-pDNA (Rep 1)
LFC_CIRc-005-pDNA (Rep 3)
LFC_CIRc-005-pDNA (Rep 4)
B
LFC_CIRc-005-pDNA
Essential sgRNAs
Negative control sgRNAs
C
CDC20
Cas9 A>G Base editor
cyclin A/Bi - Enantiomer control
Average z-scored LFC
Ser8Gly
Phe6Pro
Leu10Pro
Met1Thr
Leu16Pro
Glu7Gly
Phe4Pro
Ile280Val
CDC20
Cas9 C>T Base editor
cyclin A/Bi - Enantiomer control
Arg445Trp
Arg445Gln
Arg445Gln
Arg445Gln
D
CDK2
Cas9 A>G Base editor
cyclin A/Bi - Enantiomer control
Ile35Val
Glu42Gly
Val123Ala
Leu143Pro
Ile141Val
Lys142Gly
Trp187Arg
Leu189Pro
Phe286Pro
Phe285Ser
CDK2
Cas9 C>T Base editor
cyclin A/Bi - Enantiomer control
Glu40Lys
Glu42Lys
Gly43Asn
Glu81Lys
Ala93Val
Cys118Tyr
Arg150Lys
Ala149Thr
Ile192Ile
Pro222Leu
Thr221Ile
Ser276Leu
E
CCNA2
Cas9 A>G Base editor
cyclin A/Bi - Enantiomer control
CCNA2
Cas9 C>T Base editor
cyclin A/Bi - Enantiomer control
Ser245Phe

**Figure S8, related to Figure 4: A)** Replicate reproducibility plot of the base editor screen from Fig. 4B showing Log Fold Change (LFC) of each individual sgRNA at day 26 in cells treated with CIRc-005 (inactivate enantiomer of CIRc-004) relative to plasmid DNA (pDNA). Top panels show replicate 1 vs. replicate 2 and bottom panels show replicate 3 vs. replicate 4 for the indicated Cas9 base-editor cell lines. Pearson correlation coefficients (r) are indicated. **B)** Dot plot of Log Fold Change (LFC) of cells from **A** at day 26 treated with CIRc-005 relative to pDNA showing sgRNA depletion/enrichment of positive control sgRNAs targeting essential genes relative to negative control sgRNAs that are either non-targeting or targeting intergenic genomic DNA (negative control sgRNAs). n=4 biological replicates. **C-E**) Dot plot of average z-scored log-fold change (LFC) for each sgRNA tiling *CDC20* (C)*, CDK2* (D)*, and CCNA2* (E) in cells treated with CIRc-004 vs. CIRc-005 at day 26. The x-axis indicates the position of each sgRNA along each protein coding sequence. Variants of interest are labeled with the predicted amino acid position and mutational change. A>G base-editor is shown on left and C>T base-editor is shown on right. n=4 biological replicates.

Figure S9

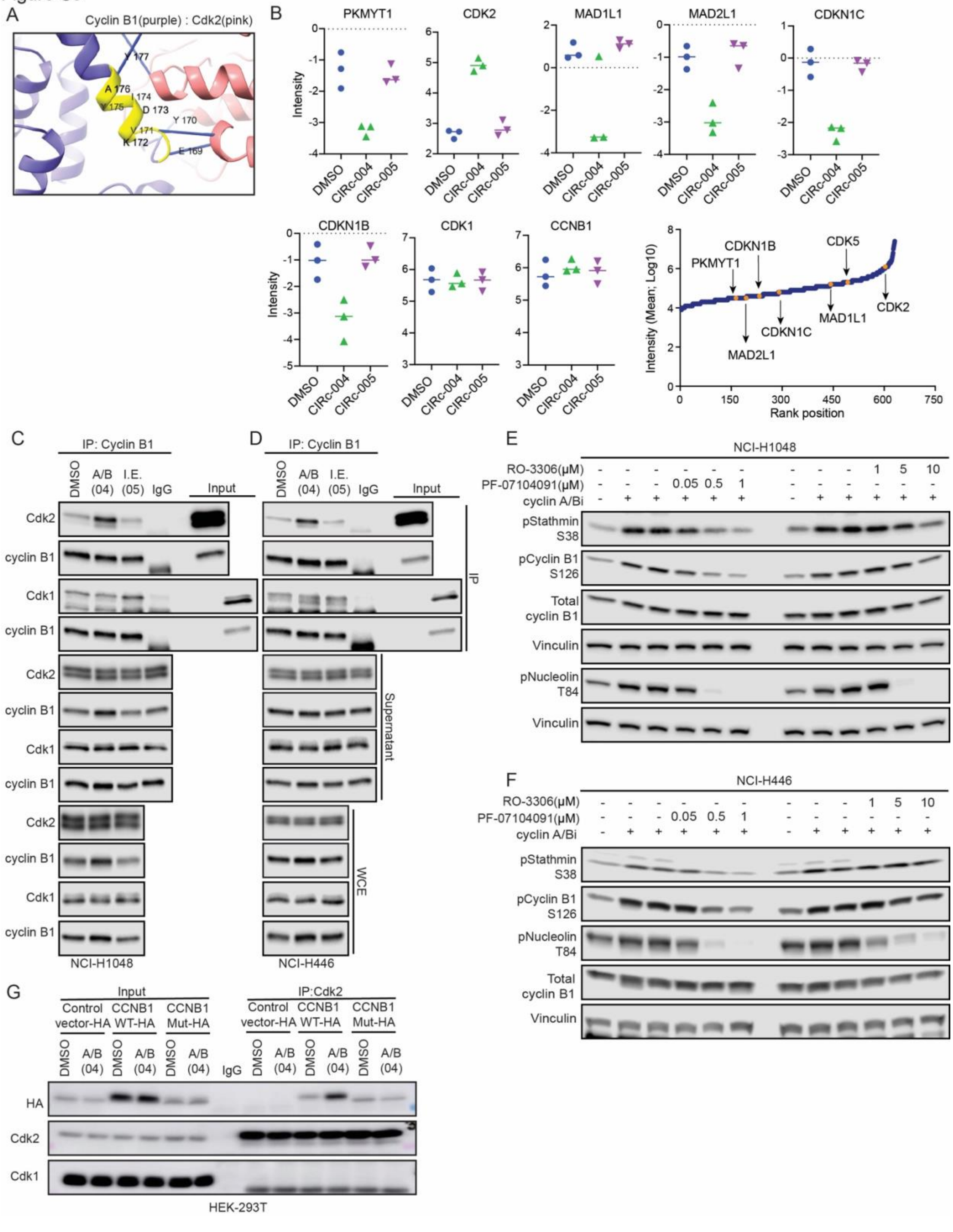
A
Cyclin B1(purple) : Cdk2(pink)
B
PKMYT1
CDK2
MAD1L1
MAD2L1
CDKN1C
CDKN1B
CDK1
CCNB1
Intensity
DMSO
CIRc-004
CIRc-005
Intensity (Mean; Log10)
Rank position
CDK5
CDK2
C
D
IP: Cyclin B1
A/B (04)
I.E. (05)
IgG
Input
Cdk2
cyclin B1
Cdk1
IP
Supernatant
WCE
NCI-H1048
NCI-H446
E
F
RO-3306(μM)
PF-07104091(μM)
cyclin A/Bi
pStathmin S38
pCyclin B1 S126
Total cyclin B1
Vinculin
pNucleolin T84
G
Input
IP:Cdk2
Control vector-HA
CCNB1 WT-HA
CCNB1 Mut-HA
HA
HEK-293T

**Figure S9, related to Figure 4: A)** AlphaFold2 model of cyclin b1:Cdk2 complex is shown. Amino acids 169-177 of cyclin b1 is highlighted in yellow showing Cdk interacting region (pink). Solid lines indicate hydrogen bonds between the two proteins with interacting amino acid residues of cyclin b1 labelled. **B**) Mass spectrometry analysis after IP of endogenous Cyclin B in NCI-H1048 cells treated with CIRc-004 (50nM), CIRc-005 (50nM) (inactive enantiomer of CIRc-004) or DMSO for 2 hours. Plots show the relative abundance of the indicated proteins as Intensity (MaxLFQ, 3VN 1 group, log 2 Median centered, missing values imputed). Data points indicate values obtained from n=3 independent biological experiments. Bottom right panel: Ranking plot of the log10 average of the 3 groups (CIRc-004, CIRc-005, DMSO) of raw protein abundance where proteins of interest (POI) are indicated within the total dataset. **C**) Immunoblot analysis after IP of endogenous cyclin B1 in NCI-H1048 cells treated with CIRc-004 (300 nM), CIRc-005 (inactive enantiomer of CIRc-004, I.E), or DMSO for 2 hours. Note that this is the same immunoblot from Fig. 4H but now includes the supernatant and whole cell lysate (WCL). **D**) Immunoblot analysis after IP of endogenous cyclin B1 in NCI-H446 cells treated with with CIRc-004 (300 nM), CIRc-005 (inactive enantiomer of CIRc-004, I.E), or DMSO for 2 hours. n=2 biological independent replicates. **E-F**) Immunoblot analysis of **E**) NCI-H1048 or **F**) NCI-H446 cells treated cyclin A/B RxL inhibitor (CIRc-004) at 200 nM, Cdk1 inhibitor (RO-3306), Cdk2 inhibitor (PF-07104091) at the concentrations indicated or DMSO for 4 hours. NCI-H1048 n=3; NCI-H446 n=2 biological independent experiments. **G)** Immunoblot analysis after IP of endogenous Cdk2 in HEK-293T cells expressing CCNB1 WT-HA, CCNB1 triple mutant-(E169K/Y170H/Y177C)-HA, or a negative control vector, and treated with CIRc-004 (300 nM) or DMSO for 2 hours. n=3 biological independent experiments.

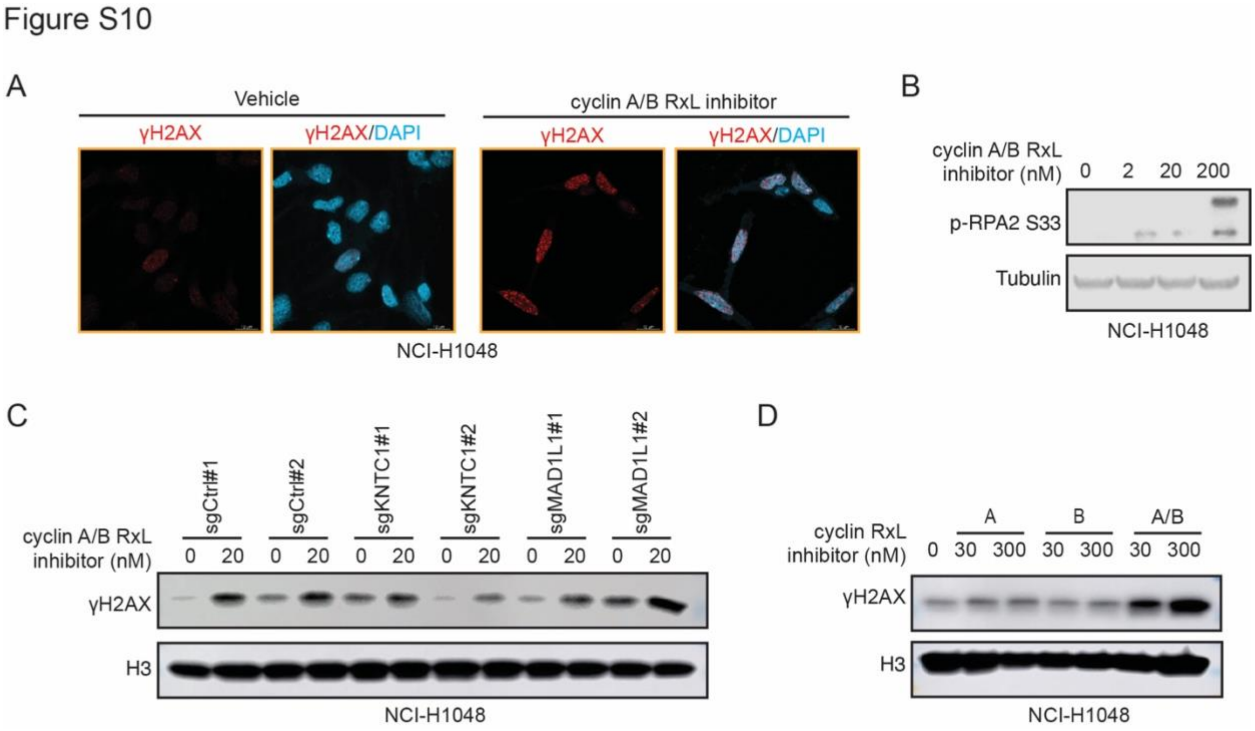


**Figure S10, related to Figure 5: A)** Representative γ-H2AX and DAPI confocal microscopic images of NCI-H1048 cells treated with the cyclin A/B RxL inhibitor (CIRc-004) at 20 nM or DMSO for 72 hours. Magnification=63x, scale bar=10 µm. **B)** Immunoblot analysis of NCI-H1048 treated with CIRc-004 at indicated concentrations for 24 hours. n=3 biological independent experiments. **C)** Immunoblot analysis of NCI-H1048 cells infected with indicated sgRNAs and then treated CIRc-004 at 20 nM or DMSO for 72 hours. **D)** Immunoblot analysis of histone lysates from NCI-H1048 cells treated with the selective cyclin A RxL inhibitor (CIRc-018), the selective cyclin B RxL inhibitor (CIRc-019), the cyclin A/B RxL inhibitor (CIRc-004), or DMSO (vehicle) for 72 hours. For **A**, **C-D**, n=3 biological independent experiments.

Figure S11

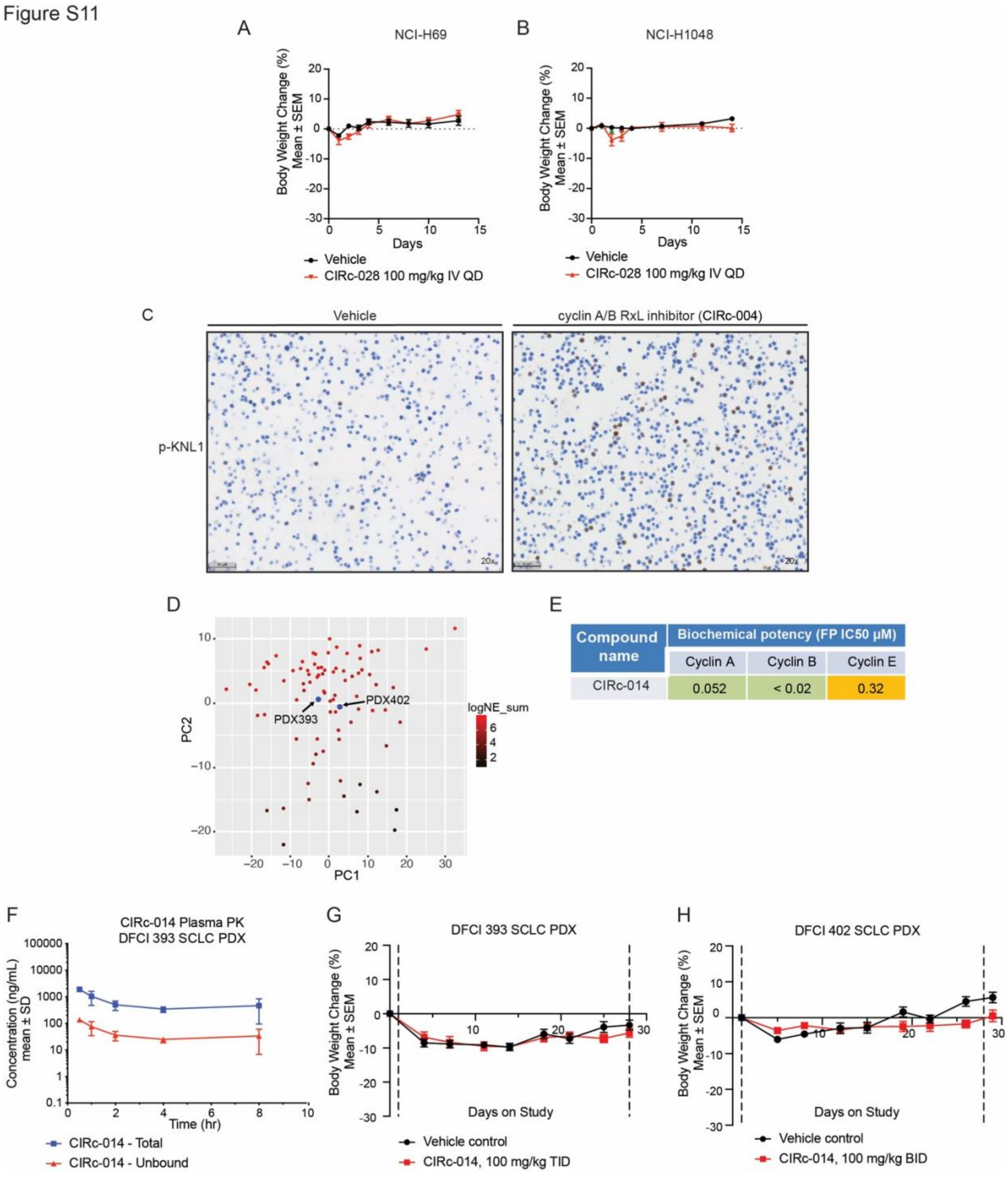
A
NCI-H69
Body Weight Change (%) Mean ± SEM
Days
Vehicle
CIRc-028 100 mg/kg IV QD
B
NCI-H1048
Body Weight Change (%) Mean ± SEM
Days
Vehicle
CIRc-028 100 mg/kg IV QD
C
Vehicle
cyclin A/B RxL inhibitor (CIRc-004)
p-KNL1
20x
20x
D
PDX393
PDX402
logNE_sum
PC2
PC1
E
Compound name
Biochemical potency (FP IC50 μM)
Cyclin A
Cyclin B
Cyclin E
CIRc-014
0.052
< 0.02
0.32
F
CIRc-014 Plasma PK
DFCI 393 SCLC PDX
Concentration (ng/mL) mean ± SD
Time (hr)
CIRc-014 - Total
CIRc-014 - Unbound
G
DFCI 393 SCLC PDX
Body Weight Change (%) Mean ± SEM
Days on Study
Vehicle control
CIRc-014, 100 mg/kg TID
H
DFCI 402 SCLC PDX
Body Weight Change (%) Mean ± SEM
Days on Study
Vehicle control
CIRc-014, 100 mg/kg BID

**Figure S11, related to Figure 6:** Body weights of mice enrolled in the **A**) NCI-H69 or **B**) NCI-H1048 xenografts efficacy treatment studies treated for 14 days with vehicle, cyclin A/B RxI inhibitor (CIRc-028, 100 mg/kg IV QD) or paclitaxel (20 mg/kg IV QOD x 5). For **A,** n=10 independent mice per arm. For **B**, n=10 independent mice for vehicle and n=8 independent mice for CIRc-028 arm. QD=Every day. **C**) Representative IHC micrographs of cell pellets from NCI-H1048 cells treated with CIRc-004 at 200 nM or DMSO (vehicle) overnight and then stained for phospho-KNL1 to validate the phospho-KNL1 antibody for IHC. Magnification=20x, scale bar=50 µm. **D**) Principal component analysis (PCA) from RNA-seq data integrating RNA-seq data from DFCI 393 and DFCI 402 PDX models with 81 primary SCLC human samples from George et al. Nature 2015 (PMID 26168399). Bar scale shows neuroendocrine score (see Methods). **E**) Biochemical activity of the orally bioavailable cyclin A/B RxL macrocyclic inhibitor (CIRc-014) for cyclin A1/Cdk2, cyclinE1/Cdk2 and cyclin B/CDK1 complexes measured by Fluorescence Polarization. CIRc-014 was used for *in vivo* efficacy studies in SCLC PDX models in Fig. 6G-N. **F**) Pharmacokinetic (PK) study in NSG mice treated with CIRc-014 for 30 min, 1, 2, 4 and 8 hours post oral dosing. Data are mean +/- SEM. Study run in NSG mice and NOD SCID % plasma protein binding (PPB) used to estimate unbound fraction. n=2 for 1-hour time point; n=3 independent mice for 30-min, 2 and 4-hour timepoints; and n=6 mice for 8-hour timepoint. **G-H**) Body weights of mice enrolled in the **G**) DFCI 393 SCLC PDX or **H**) DFCI 402 SCLC PDX efficacy treatment studies treated for 28 days with CIRc-014 (100 mg/kg PO TID for DFCI 393 and 100 mg/kg PO BID for DFCI 402) or vehicle. For DFCI 393 PDX in **I**, n=10 independent mice for vehicle and n=8 independent mice for CIRc-014 arm. For DFCI 402 PDX in **J**, n=10 independent mice per arm for both DFCI 393 and DFCI 402. TID=three times a day; BID=two times a day.

Figure S12

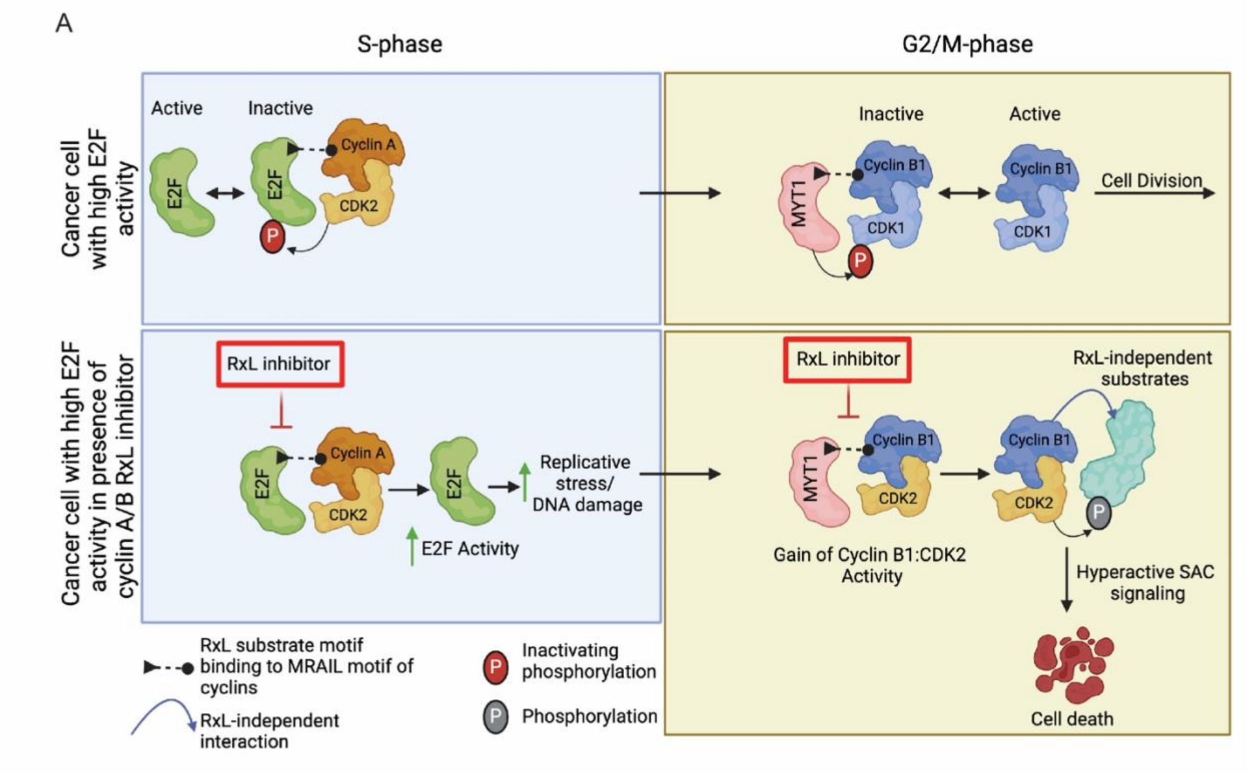


**Figure S12: Proposed mechanism of action for cyclin A/B RxL macrocyclic peptide inhibitors in cancers with high E2F activity. Top:** In a cancer cell with high baseline E2F activity, cyclin A binds to activating E2Fs by recognizing an RxL-motif on E2F facilitating cyclin A-Cdk2 to phosphorylate E2F, which decreases E2F activity allowing cells to progress from S-phase to G2/M-phase of the cell cycle. At the beginning of G2, cyclin B binds to Myt1 by recognizing an RxL-motif on Myt1 and Myt1 phosphorylates Cdk1, which inhibits cyclin B-Cdk1 activity until cells progress to M-phase where cyclin B-Cdk1 is active and controls mitotic fidelity. **Bottom**: When a cancer cell with baseline high E2F activity is treated with a cyclin A/B RxL inhibitor to block these RxL-dependent interactions, the cyclin A-E2F1 RxL-dependent interaction is disrupted leading to increased E2F activity which increases replication stress as the cancer cell moves from S-phase to G2/M-phase of the cell cycle. In G2/M, the cyclin B-Myt1 RxL-dependent interaction is also disrupted and Cdk2 is redirected to cyclin B to form a neomorphic cyclin B-Cdk2 complex with gain of function activity leading to phosphorylation of RxL-independent substrates, prolonged spindle assembly checkpoint (SAC) activation, and mitotic cell death. This mechanism of action of cyclin A/B RxL inhibitors requires high baseline activity of E2F observed in cancer cells with genetic alterations that compromise the negative regulation of E2F, such as *RB1* inactivation, often in the setting of concurrent *TP53* inactivation, which is observed in almost all SCLCs. This figure was created with Biorender.
